# Ideal efficacy photoswitches for TRPC4/5 channels harness high potency for spatiotemporally-resolved control of TRPC function in live tissues

**DOI:** 10.1101/2024.07.12.602451

**Authors:** Markus Müller, Konstantin Niemeyer, Navin K. Ojha, Sebastian A. Porav, Deivanayagabarathy Vinayagam, Nicole Urban, Fanny Büchau, Katharina Oleinikov, Mazen Makke, Claudia C. Bauer, Aidan V. Johnson, Stephen P. Muench, Frank Zufall, Dieter Bruns, Yvonne Schwarz, Stefan Raunser, Trese Leinders-Zufall, Robin S. Bon, Michael Schaefer, Oliver Thorn-Seshold

## Abstract

Directly probing the endogenous biological roles of target proteins with high spatial and temporal resolution, as non-invasively and reproducibly as possible, is a shared conceptual goal for research across many fields, as well as for targeted therapies. Here we describe the rational conceptual design and test-case practical implementation of a photopharmacological paradigm to empower high-performance photomodulation studies *in vivo*. TRPC4/5 ion channels are involved in many spatiotemporally resolved circuits, from pain and anxiety, to reproductive signaling, digestion, and obesity. To unpick their biology requires spatiotemporally precise tools, which were lacking. We developed “ideal efficacy photoswitch” ligands to control their diverse functions *in situ*. These *E*⇆*Z-*photoswitchable ligands bias TRPC[4]/5 channel activity with exquisite photocontrol, from strong agonism under 360 nm, to low agonism at 385 nm, to strong antagonism at 410-460 nm. Cryo-EM structures of both TRPC4 and TRPC5 with both *Z-*agonists and *E-*antagonists support the rationale for efficacy switching through competitive *E/Z* isomer binding. Crucially, since the *E/Z* ratio is exclusively determined by the light wavelength applied, **their channel photocontrol is exclusively wavelength-dependent, yet drug-concentration-independent: so is reproducible from cell culture to >millimetre-depth tissues**. Indeed, we were able to photocontrol both direct and downstream TRPC4/5 biology in cell lines or primary cells in culture, from calcium flux, to primary neuron excitability and adrenaline release; and even in tissues, photoswitching small intestine motility and peristalsis. The TRPC4/5 ligands we develop will thus unlock a range of high-precision investigations in TRP biology. More broadly, we propose that the success of this efficacy photoswitch program, from concept to tissue level translation, is mainly a consequence of how biology has evolved proteins for efficacy control. We therefore foresee that a variety of functionally responsive protein targets, not only sensory and signaling ion channels and receptors, will be amenable to similarly high-performance photocontrol even *in vivo*, if a new generation of reagent development adopts this paradigm of **ideal efficacy photoswitching**.

**Table of Contents Graphic:** 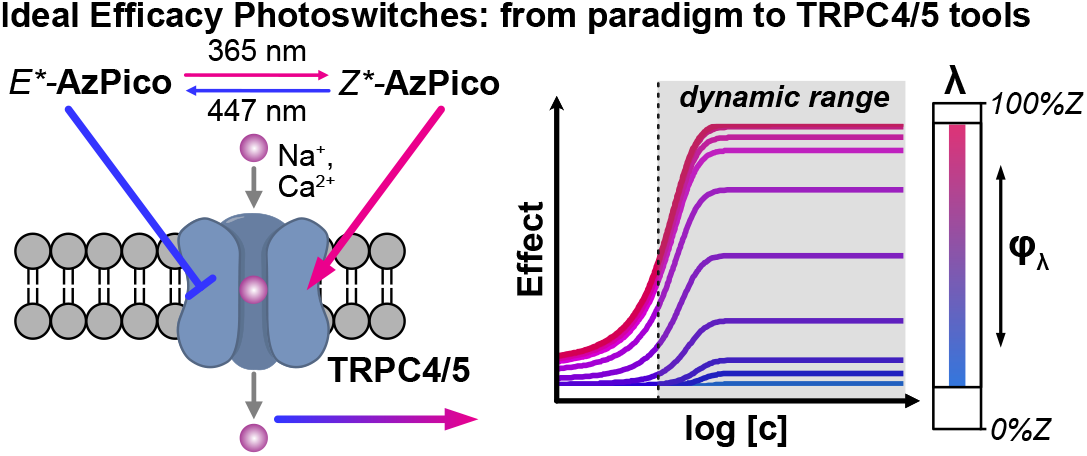

## Introduction

The 27 human Transient Receptor Potential (TRP) proteins can assemble to form tetrameric TRP ion channels, which have varied and highly tissue-dependent roles in cellular physiology: from sensing and signaling to mineralostasis.^1,2^ Several environment-sensing TRP channels such as TRPV1 (heat), TRPM8 (cold), or TRPA1 (electrophiles), are well-studied. For many other TRP channels their roles and significance remain unclear, even if their links with diseases make them hotly pursued as therapeutic targets.^3–5^ TRP canonicals (TRPC) are a subfamily of non-selective Na^+^/Ca^2+^-permeable cation channels; including the structurally similar TRPC4 and TRPC5.^6–8^ TRPC4/5 are mainly expressed in the central nervous system (CNS), gut, and kidneys, and are functionally linked with pain,^9–11^ reproductive signaling,^12^ anxiety and depression,^13,14^ kidney disease,^15^ and digestion;^16^ and striking discoveries are still ongoing e.g. linking TRPC5 loss to postpartum depression and obesity.^17^ TRPC4/5 form both homotetrameric, and heterotetrameric functional channels that can also include TRPC1.^18^

Insights into TRPC4/5 relevance were initially derived from knockout mice because potent and selective modulators were lacking.^19^ Recently, small-molecule modulators of TRPC4/5 have become available as tool compounds,^20^ starting with the nanomolar-potency natural product (-)-**Englerin A** (**EA**),^21^ a TRPC1/4/5 activator,^22,23^ and complemented by TRPC4/5 inhibitors developed as drug candidates.^24,25^ However, TRP functionality varies according to tissue localisation (spatial), and TRPs are best studied if their activity is reversibly modulated on short timescales (temporal); therefore, photoresponsive chemical tools, whose activity can be spatiotemporally patterned by light, are particularly well-adapted to resolve TRP biology.^26^ In general, photoswitchable TRP ligands have been impactful: with azo-capsaicins for TRPV1,^27,28^ TRPswitch for TRPA1,^29^ OptoBI for TRPC3,^30^ and azo-diacylglycerols (PhoDAGs and Opto-DArGs) for TRPC2/3/6.^31–33^ However, there is no photoresponsive ligand for TRPC4, and the sole compound for Trpc5 (**BTDAzo**, a lipophilic photoswitchable agonist) has low potency and is almost inactive on human TRPC5^34^.

The highly potent and selective TRPC4/5 drug leads from industry, e.g., TRPC5-targeting pyridazinones GFB-8438 and GFB-887 developed by GoldfinchBio, and TRPC4/5-targeting xanthines **HC-070** (in clinical trials as BI-1358894) and **Pico145** (also named HC-608) developed by Hydra/Boehringer (**Figure 1a**), have already driven major progress in basic TRPC4/5 research (**Supporting Note 1**). In this work, we sought to create photoswitchable analogues of them, to add spatiotemporal resolution on top of their potency and selectivity, and so make powerful tools for TRP studies in complex biological systems. We particularly sought to address TRPC4, given its lack of photoswitchable tools. Crucially, we were drawn to the xanthines for conceptual reasons, since their structure-activity relationships (SAR) reveal activators and inhibitors with high structural similarity,^35,36^ which in turn suggested the rare opportunity to harness an efficacy tipping point to obtain high-performance *efficacy switch* tools. We now report the design and experimental validation of this concept, use these tools for light-controlled high-spatiotemporal-precision elucidation of the separate roles of TRPC4 and/or TRPC5 from cell culture through to endogenous organ sections, and argue more broadly for the adoption of this efficacy switch paradigm as a way to unlock high precision *in vivo* photocontrol of a range of biological systems.

**Fig 1.**
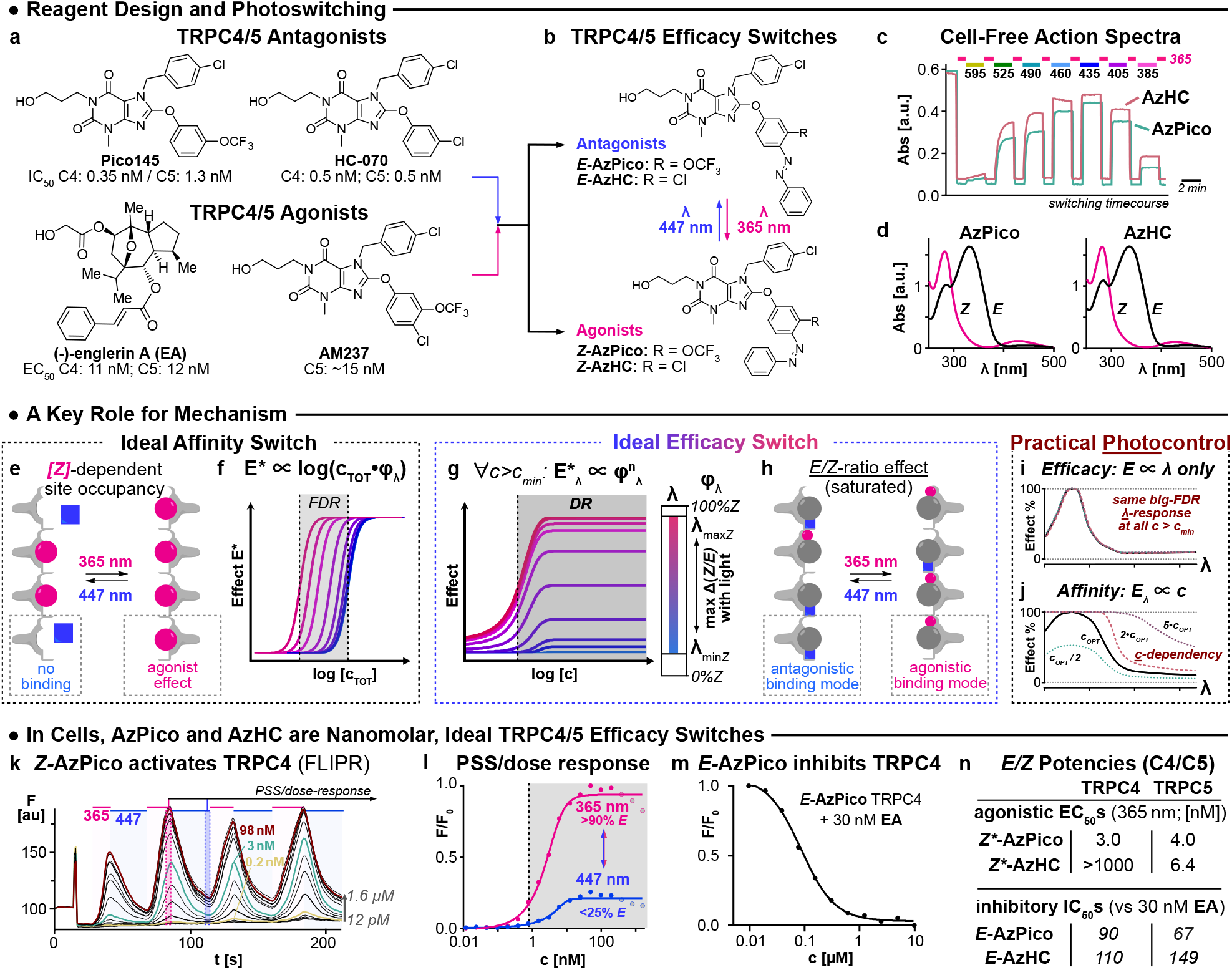
Efficacy photoswitches for TRPC4/5. **a**, Known TRPC4/5 modulators. **b**, Photoswitchable TRPC4/5 modulators **AzPico** and **AzHC. c-d**, Photoisomerisation action spectra and *E/Z* isomer absorption spectra of **AzPico** and **AzHC. e**,**f**, For an ideal affinity switch, only one isomer binds the target. Within the FDR window, binding site occupancy and thus biological effect **E\*** depend on the total switch concentration **c**_**TOT**_ and the PSS fraction of active isomer **f**_**λ**_. **g**,**h**, For an ideal efficacy switch, both isomers bind, with similar affinities, but with different efficacies. The dynamic range (DR) where the biological effect E*_λ_ is PSS-dependent but concentration-independent covers all c > c_min_. **i**,**j**, Photocontrol in practice: for an efficacy switch, small variations in PSS(λ) sensitively control performance (whereas in affinity switches they are unimportant); but even large variations in concentration, which would ruin the performance of an affinity switch, are irrelevant. **k**,**l**, Reversible Ca^2+^ influx modulation with **AzPico** under 365/447 nm cycles, as timecourse and peak amplitudes. **m**, *E*-**AzPico** binds competitively to **EA. n**, EC_50_ and IC_50_ values of *E/Z*-**AzPico & AzHC** on TRPC4 and TRPC5 (**k-n**, Fluo-4-loaded HEK_mTRPC4β_ cells).

## Results

### Design Concept: Efficacy Switching (Figure 1e-j)

Azobenzenes are the best-explored chemical scaffolds for fully reversible structural photoswitching (*E*⇆*Z* isomerisation),^37^ and we determined to employ them in ligands to photoswitch TRPC4/5 bioactivity. However, azobenzene *E*⇆*Z* photoswitching is never complete in both directions: it operates between mostly-*Z* and mostly-*E* photostationary states (PSSs, typically ca. 90%*Z* for the most-*Z*-PSS [*Z\**], and 80%*E* for the most-*E*-PSS [*E\**]).

**Affinity Switching** is the typical rational design concept for photopharmacology (ca. >95% of all photoswitches reported), where differences in binding affinity based on steric clash/fit drive a difference in potency of *E/Z-*isomers; so net *E*⇆*Z* photoswitching modulates the biological activity applied. However, due to the typically 20% residual *Z* isomer in the mostly-*E*-PSS, even an *ideal* affinity switch that has a completely non-binding *E*-isomer but a high-agonist-potency *Z*-isomer (**Figure 1e**) suffers substantial background activity after *Z*→*E* photo-switching-off. Conceptually, this is covered in the general result that bidirectional photocontrol of an affinity switch can only deliver a narrow *functional dynamic range of bioactivity* (FDR),^38,39^ i.e. the bioactivity window between best-photoswitched-on and best-photo-switched-off states (**Figure 1f**; the FDR is limited by its PSSs in both directions). Worse still, the applied bioactivity of an affinity switch under a fixed wavelength is *extremely* sensitive to its concentration (**Figure 1j**), such that both dose and wavelength must be dynamically balanced to deliver a given effect. This is problematic or impossible for tissue or *in vivo* work, due to highly variable reagent concentrations (ADME-PK, distance from blood vessels, inter-animal variability, etc). We conjecture that this explains why affinity switches’ applicability for bidirectional photocontrol has been mainly limited to highly controlled cell culture settings^40^.

**Efficacy Switching** is the design concept we harness in this work. We formulate the goal^41^ of an efficacy photoswitch to be, that *E/Z* photoisomers have different modes of bioactivity on the target, while still binding competitively with similar affinity. Ideally, *E*/*Z* isomers have either (i) opposing activity (e.g., activator/inhibitor), or (ii) one isomer is biologically innocent, but prevents the other from acting by its competitive binding (**Figure 1h**). Here, the novelty of installing such a nanomolar, ideal efficacy switch on TRPC4/5-binding scaffolds will be the chemical highlight of this paper, and is the key feature that allows their *in vivo* biological applications. First however, it will be crucial to trace the logic of efficacy switching, to see why its opportunities and advantages arise, since this conceptual issue of *how* a compound exerts its photocontrol has not received proper formal attention (especially compared to e.g. how much literature is devoted to easily-measured but less critical aspects such as incremental tuning of photoresponse wavelengths and PSSs).

For an ideal efficacy switch, above a threshold concentration where its target nears saturation by an *E/Z* mixture, there is no more concentration-dependency of bioactivity. The only variable controlling bioactivity is the *E/Z* ratio of the competitively-binding isomers: i.e., (1) the PSS entirely controls the bioactivity. Setting the PSS by choosing the wavelength of light applied is far easier and more reproducible than controlling drug biodistribution or concentration: making efficacy switches ideal to tackle complex environments despite variable biodistribution, and suiting them to easy translation between model systems. Also, (2) efficacy switches thus access an *open-ended dynamic range* (DR), where switching between ON and full baseline states is identically possible at *any* concentration above the threshold, as long as light is applied efficiently (**Figure 1gi**). These features should make efficacy photoswitches much better-suited for robust use in biology, from cell culture through to *in vivo* (**Figures 3-6**). However, only about ten cases of efficacy photoswitching have been published (chemokine, adenosine, cannabinoid, and serotonin receptors). Key contributions include those of Decker, Leurs, Gorostiza, and coworkers (discussed in **Supporting Note 2**).^42–51^ With the exception of Leurs’ micromolar chemokine photoswitch ligand VUF16216,^43,48^ efficacy switching on the other known targets can be considered “non-ideal” in that **(a)** the ligand efficacies were switched between more- and less-activating, rather than activating and inhibiting, and **(b)** the isomers’ affinity was rather different, which compromises the concentration-independence of ideal efficacy switching. It is also certain that other efficacy switches have been created, but simply not reported as such; e.g., we also published a reagent with excellent dynamic range of bioactivity photocontrol despite its *E*⇆*Z* photoswitching incompleteness, but had not understood the full consequences of that performance at the time^34^ (while re-parsing recent photopharmacology papers, we believe we have found many more instances of unsuspected efficacy switches; **Supporting Note 2** discusses this further with some examples from Fuchter, Groschner, Pepperberg, Trauner, etc^52–56^).

We give this level of detail since we believe that efficacy photoswitching is *the* mode needed to reach the naively popular picture of “solely wavelength-controlled bioactivity” that has motivated much of photopharmacology, but, we did not find it collected accessibly in the literature elsewhere. The conceptual importance of this framework goes deeper than pharmacology though. As one example, we highlight that there are *target-driven* reasons to choose efficacy switching for proteins which are natively poised for steeply nonlinear dose-response switching between metastable states, such as receptors and ion channels with multiple binding sites. These ought to be ideal platforms where competitive binding of similar-affinity *E/Z* isomers with opposing modes of action ought to deliver a concentration-independent, *true binary photoswitch for protein activity*, even though ligand photoswitching is never complete. A separate theoretical paper will treat these aspects in detail^57^; but for now, we set out to test this concept in practice, by creating such “ideal efficacy switch” reagents for TRPC4/5.

### Xanthine Efficacy Switches: Design and Synthesis (Figure S3)

Xanthines **Pico145** (also called **HC-608**) and **HC-070**^58^ are TRPC1/4/5 antagonists with picomolar potency^59^ and remarkable selectivity against hundreds of enzymes, receptors, transporters and other ion channels (including other TRP channels).^24^ Excitingly, the very similar **AM237**^58^ is instead a nanomolar *agonist* of homotetrameric TRPC5, despite also being a nanomolar antagonist of homotetrameric TRPC4 (for full details of the pharmacology, see **Supporting Note 1**).^35^ The structures of these different-efficacy compounds are nearly identical (**Pico145**: *m-*OCF_3_, **HC-070**: *m-*Cl; **AM237**: *m*-OCF_3_, *p*-Cl; **Figure 1a**). This suggests that the *meta/para* positions are suitable as an “ideal efficacy tipping point”: small modifications may flip the efficacy mode (activator/inhibitor) *without* changing the binding affinity. Thus, the xanthines seemed an outstanding starting point for ideal efficacy switches that are *also* highly potent, and so can be reliably applied *in vivo*.

In brief, we synthesised a series of xanthines “extended” with bidirectionally switchable azobenzenes (**Figure S3**). Noteworthily, with a simple -NNPh motif in *para* (where **AM237** has a -Cl), we obtained **AzPico** (*m-*OCF_3_) and **AzHC** (*m-*Cl; **Figure 1b**) which were soon identified as the most biologically useful candidates in our panel of eight. From here onwards we will focus only on them, leaving the others to **Supporting Note 3. AzPico/AzHC** could be reversibly photoswitched between PSSs of ca. 82%*E* around 410 nm and 95%*Z* around 360 nm (**Figure 1cd** and **Table S3**).

### Parallel-Throughput Photocontrol Assessment in Cells by FLIPR (Figure 1k-n)

We initially screened for photoswitchability of activity in cells, using a fluorometric imaging plate reader (FLIPR) calcium flux assay, with HEK293 cells stably expressing mouse TRPC4β or mouse TRPC5. The parallel-throughput FLIPR setup is limited to use fixed LEDs at 365 nm (good-*Z*) and 447 nm (suboptimal-*E*), during imaging at 490 nm (so imaging counteracts both PSSs). Therefore, these FLIPR results will underestimate the true photocontrol the reagents can access, as shown later.

*E-***AzPico** from 12 pM to 1.6 µM gave no effect with TRPC4 or TRPC5; upon 365 nm illumination, strong agonism was evident with Ca^2+^ influx rapidly evoked at low nanomolar concentrations; and this was rapidly photoreversible with 447 nm illumination, over many cycles (**Figure 1k**). The remarkable observation that the maximum and minimum calcium signals are dose-independent over a concentration range of >100-fold (from nM to µM), even though the concentration of the agonistic *Z* isomer likewise increases by >100-fold in this range, confirms it as an *ideal efficacy switch* (TRPC4 in **Figure 1gl**; TRPC5 in **Figure S2** (*E*) and **Figure S13** (*Z*)). Pleasingly, it inherits the high potency of its parent **Pico145**, with 365 nm EC_50_ of just 3.0 nM on TRPC4 and 4.0 nM on TRPC5 (**Figure 1n**). **AzHC** was also a *Z-*agonistic ideal efficacy switch for TRPC5 with a 365 nm EC_50_ of 6.4 nM. Excitingly though, *Z-***AzHC** was completely inactive on TRPC4 into the µM range (**Figure S13**). Competition assays supported that these compounds are indeed efficacy switches (**Figure 1m** and **Supporting Note 1**).

Taken together, this pair of reagents are uniquely potent efficacy switches; **AzPico** is the first photoswitchable tool to address TRPC4; and although **AzPico** acts on TRPC5 as well, it can be applied comparatively to the TRPC5-selective **AzHC** to test the cellular role of TRPC4: making them an outstanding reagent pair for probing these otherwise difficult to resolve channels. Their retention of the nanomolar potency of their optimised parent compounds is also a rarity among photopharmaceuticals, because the extra moiety needed for photoisomerisation usually sacrifices potency.

**Photomodulated electrophysiology (Figure 2)**

**Fig 2.**
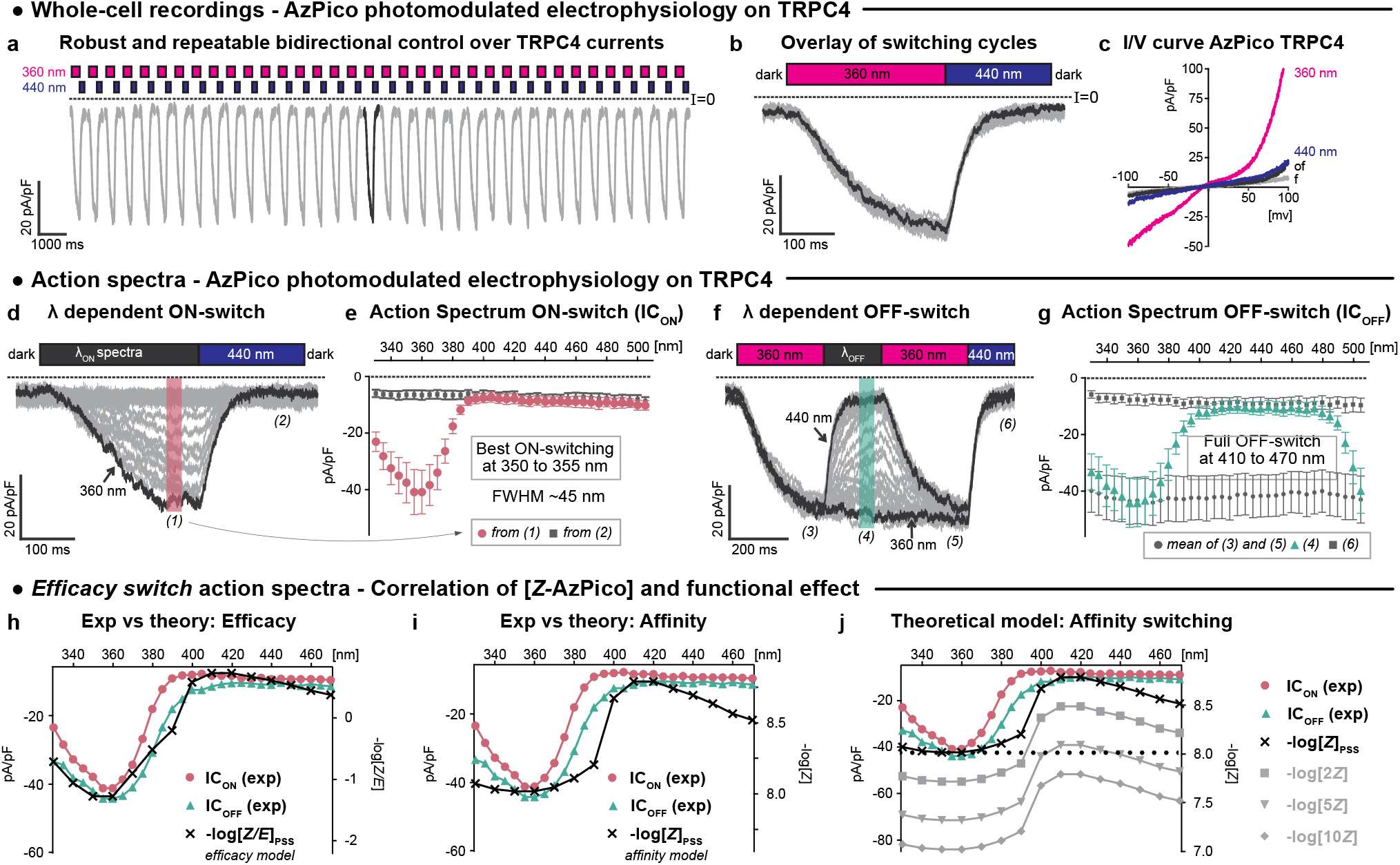
AzPico-photocontrolled electrophysiology of TRPC4. **a-g**, Electrophysiological whole-cell recordings of TRPC4 currents in voltage clamp mode (**a**,**b**,**d-g**: *V*h = -80 mV; **c**: *V*h scan) in HEK293 cells with 10 nM **AzPico** during photoswitching. **a-b**, Reproducibility of 36 consecutive photoswitching cycles of 360/440 nm (**a**: time-course; **b**: overlay of all cycles). **c**, *I*/*V* curves show that 440 nm drives almost full return to baseline currents throughout the applied voltage range. **d-g**, Spectral scans to extract the wavelength dependency of channel current photoswitch-on (**d-e**, cycles of λ_ON_/440 nm) and photoswitch-off (**f-g**, 360 nm/λ_OFF_). **h-j**: Ephys action spectra of **AzPico** match PSS-informed expectations for an efficacy switch (**h**), not an affinity switch (**i**): important, since an affinity switch would have severe concentration dependency (**j**). Full legend at **Figure S24**.

**Fig 3.**
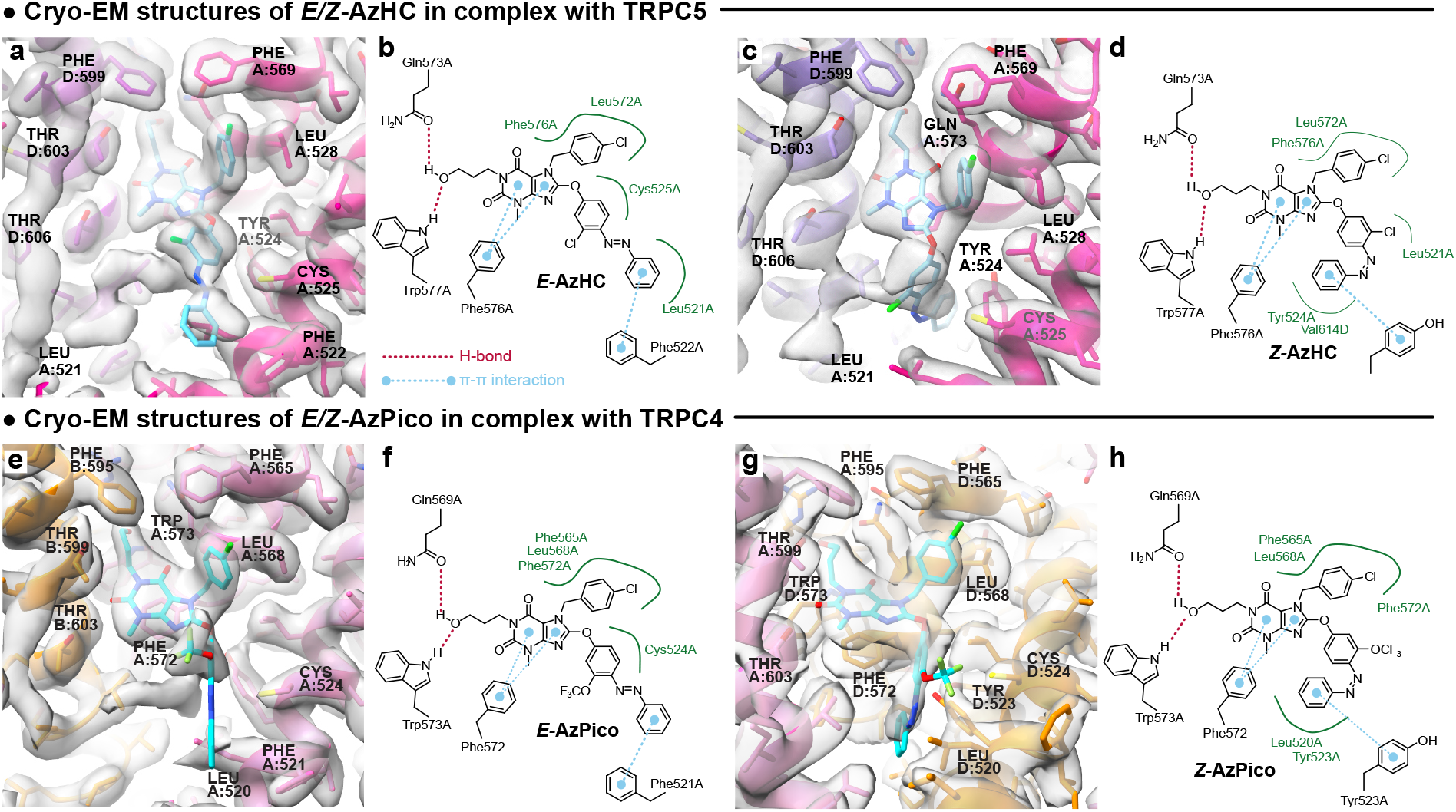
Structures of TRPC4/5 in complex with *E/Z-*isomers of efficacy photoswitches. **a-b**, hTRPC5:*E*-**AzHC** (2.6 Å, PDB xx, EMDB xx), with 2D map of ligand-protein interactions. **c-d**, hTRPC5:*Z*-**AzHC** (2.9 Å, PDB xx, EMDB xx). Note the flip of the distal azobenzene ring. **e-g**, TRPC4_DR_:*E/Z*-**AzPico** (*E:* 3.0 Å, PDB 9FXL, EMDB 50850; *Z:* 3.1 Å, PDB 9FXM, EMDB: 50851). Full legend at **Figure S25**.

We next performed electrophysiological patch clamp experiments (ephys), to characterise the action and specificity of channel modulation in more detail. Though its throughput is lower, ephys is more powerful than FLIPR in several respects: (i) the non-optical ephys readout does not cause unwanted photoswitching, unlike FLIPR; (ii) ephys readouts linearly and temporally resolve ionic currents through the activated channels, unlike the delayed and attenuated fluorometric Ca^2+^ influx analyses in FLIPR; (iii) the ephys setup can use narrow-bandwidth monochromated light at any wavelength. These features allow measuring the full power of the photoswitch reagents.

We thus recorded wavelength-dependent action spectra of **AzPico** and **AzHC** under fast photoswitching in ephys in TRPC4 / TRPC5-expressing cells (**Figure 2, Figure S14**). They now fulfilled the potential of ideal efficacy switches. For example, **AzPico** reversibly photomodulated TRPC4 currents over >36 consecutive cycles with a fully constant activation profile, without fatigue (**Figure 2ab**). Current/voltage (I/V) curves show a strong activation of ion flux at best-*Z* (360 nm) PSS; yet, only at nonphysiological voltages (<-80 mV or >+60 mV) could any small differences between basal activity and good-*E* (440 nm) PSS be detected (**Figure 2c**): making them truly off/on TRPC modulators.

The repeatability of on/off photocycling allowed us to extract action spectra for both (1) photoactivation (**Figure 2de**) and (2) photodeactivation (**Figure 2fg**) *in situ* in live cells. Channel currents were optimally photoswitched-on in a sharp wavelength range of 340-370 nm; and *full* photoswitch-off was triggered over the broad range of 400-480 nm (best: 430 nm). Such action spectra conclusively match the highly wavelength-sensitive model expected from cell-free PSS measurements for an ideal efficacy switch (bioactivity ∝ -log_10_([*Z*]/[*E*])), but mismatch the model for an affinity switch (bioactivity ∝ -log_10_([*Z*]) (match/mismatch is visible even for single concentration data: **Figure 2hi**).

Importantly, we recall that only the efficacy switch mechanism allows these reagents to deliver binary off/on bioactivity, that is identically wavelength-dependent at any concentration (**Figure 1ghi, Figure 2h**); e.g., if they were classical affinity switches, even doubling their concentration would prevent photo-switching-off cell currents (**Figure 2j**; **Supporting Note 1**). Thus, the concentration-scan, fixed-wavelength FLIPR plus the fixed-concentration, wavelength-scan ephys (**Figures 1kl, 2egh, S14**) show that **AzHC** and **AzPico** are ideal efficacy switches, whose *E/Z-*competitive binding controls TRPC4/5 currents with exquisite wavelength-sensitivity and concentration independence.

### Paired Pairs of Cryo-EM TRPC4/5:*E/Z* Structures Indicate Activation Mechanism (Figure 3)

As **AzHC** and **AzPico** should bind TRPC4/5 with high affinity in both *E-* and *Z-*forms, this offers a rare opportunity for structural photopharmacology to elucidate the basis of efficacy switching (to date, there are only two protein-ligand structure pairs with an azobenzene reagent bound as both *E-* and *Z-*isomers^49,60^); and potentially to understand the remarkable differential of TRPC4 activity induced by the -Cl/-OCF_3_ swap. Single-particle cryo–electron microscopy (cryo-EM) studies of TRPC4^61–63^ and TRPC5^64–67^ have already given important insights into the structures of these channels and their complexes with lipids, metals, proteins, and small-molecule modulators. These include TRPC5 structures with **Pico145**^66^ and **HC-070**,^64^ binding in near-identical pose to the same lipid binding pocket, adjacent to the pore helices. However, no channel-open structures are known, and all reported ligand structures are with inhibitors.^36^

We now studied TRPC5:*E/Z-***AzHC** and TRPC4:*E/Z-***AzPico** complexes by cryo-EM, hoping to acquire the “paired pair” of both inhibited and activated structures for both channels. This work was run fully independently for TRPC4 at one site, and TRPC5 in another; yet the results aligned, giving confidence in their interpretations. We could determine all four structures at high resolution: **hTRPC5:***E*-**AzHC** (2.6 Å), **hTRPC5:***Z*-**AzHC** (2.9 Å), **TRPC4**_DR_: *E*-**AzPico** (3.0 Å), and **TRPC4**_DR_: *Z*-**AzPico** (3.1 Å), without imposing symmetry during data processing (**Figure 3**; **Figures S15-S19**; **Tables S4-S7**; hTRPC5 = human TRPC5, TRPC4_DR_ = zebrafish TRPC4). We used 365 nm for *Z-*ligand structures, and either dark or 440 nm for *E-*structures, and excluded DTT from buffers to avoid diazene reduction.

**Supporting Note 4** contains full discussions of the structural biology results, but in brief, *E/Z-***AzHC**/**AzPico** could be built into the expected lipid/xanthine binding site, with near-identical positions as for **Pico145**^68^ for the conserved ligand portion, and differences between the *E* and *Z* isomers only at the azobenzene. E.g., the distal ring of *E*-**AzHC** is projected outwards to make a π-π interaction with Phe522 of TRPC5, while in *Z*-**AzHC**, it folds deeply inwards to make a π-π interaction with Tyr524; **AzPico** behaves similarly on TRPC4 (**Figure 3**; **Figures S4, S6**). Most protein residues in the ligand binding site are in similar positions in the *E/Z* structures; the exceptions are that for hTRPC5, Phe520 is flipped “in” or “out” depending on whether the antagonistic *E* or agonistic *Z* isomer is bound (**Figure S4b**), while in TRPC4_DR_ the cognate Phe (521) is much less shifted, though its neighbouring Leu520 is significantly displaced. All *E* and *Z* structures had the channel pore closed (see **Figures S5** and **S7**, which suggest potential reasons for this), so caution in interpreting the agonist structures is needed; nevertheless, these data offer the first structural insight into how closely related xanthines can have opposite effects on TRPC4/5 function (i.e. inhibition vs. activation).

With these binding modes confirmed, we tested whether the selective activation of TRPC5 but not TRPC4 by *Z-***AzHC** could be the result of their binding sites’ single amino acid difference (TRPC5 has Val579 where TRPC4 has Ile575 at the cognate position). However, neither the TRPC5→C4 mutation (V579I) nor the TRPC4→C5 mutation (I575V) changed their activity profiles for *E/Z-***AzHC** or for control activator **AM237**. This suggests that the basis for *Z-***AzHC**’s TRPC5-selective activation is more complex than the immediate residues it contacts (**Supporting Note 4**).

### Photoswitching endogenous TRPC4/5 in primary cells to photoreversibly actuate cell function (Figure 4)

We next moved to test whether **AzPico** can directly photocontrol endogenous TRPC4/5, using autaptic hippocampal neurons (neurons cultured in isolation which only make synapses back onto themselves, as a model for simultaneously monitoring pre- and post-synaptic responses). 365/460 nm cycles reversibly activated inward currents in wildtype (**wt**) neurons, just as was seen in heterologously TRPC4/5-[over]expressing HEK cells (**Figure 2**). These **wt** neurons consist of a fraction of cells expressing TRPC channels that can be directly activated by **AzPico**, plus about 50% of neurons that do not express TRPC channels.^69^ Matching expectations, average currents were larger when acquired exclusively from TRPC5-bearing cells (**5ki**); strongly depressed with TRPC5 single knockout (**5ko**); and almost abolished with TRPC1/C4/C5 triple knockout (**145tko**); supporting that TRPC[1]4/5 channels are the central contributor to **AzPico**’s photomodulation of membrane conductance (**Figure 4a**).^70^ This shows that **AzPico** can optically control the activity of native TRPC[1/4/]5 channels in primary nerve cells, without channel over-expression.

**Fig 4.**
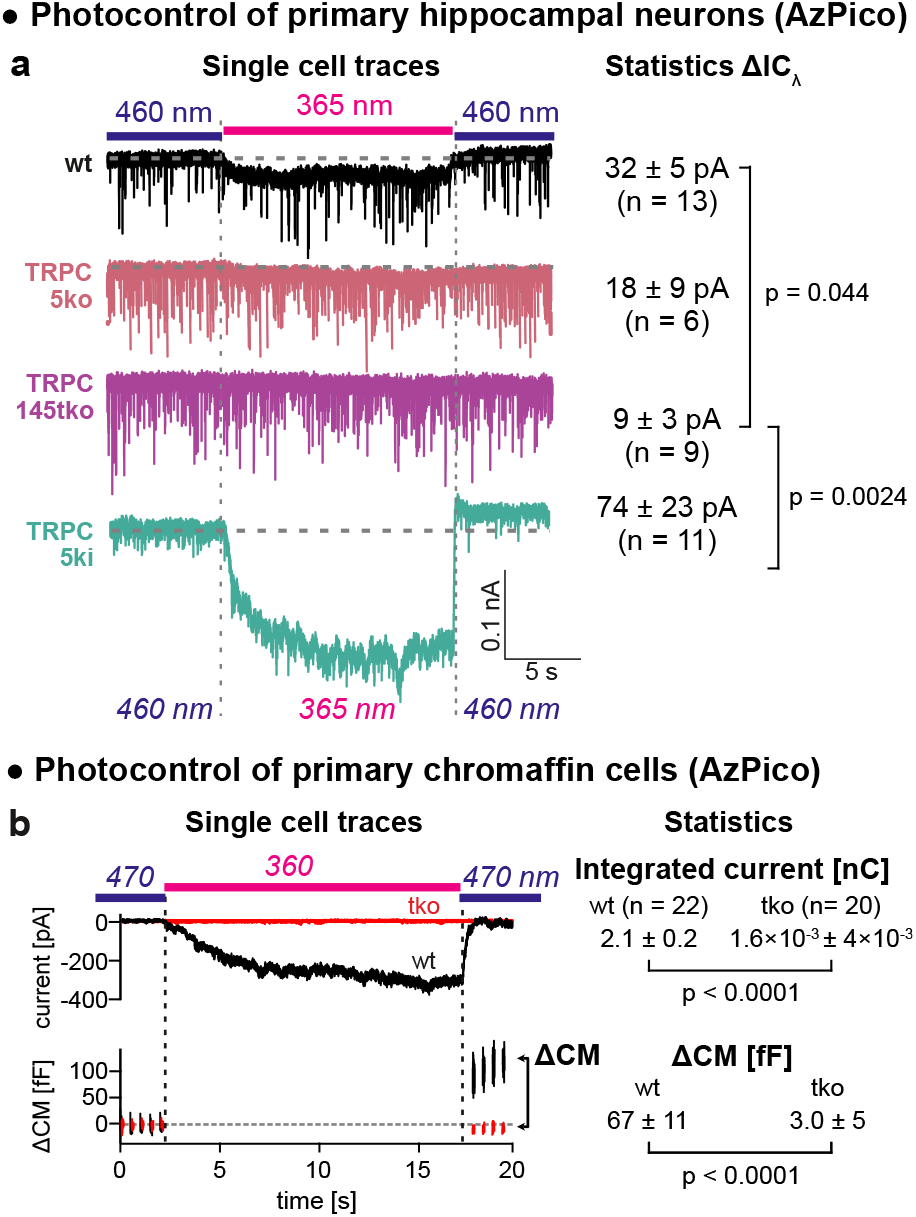
AzPico. (30 nM) photoswitchably evokes currents in primary neuronal and neuroendocrine cells with endogenous TRPC levels (single cell traces at left, group statistics at right). **a**, photoswitching-based current differentials **ΔIC**_**λ**_ in hippocampal neurons (currents at 365 nm relative to 460 nm). **b**, photoswitch-based currents and net charge transfer (top) correlate to membrane capacitance changes **ΔCM** (bottom), indicating that phototriggering of TRPC[1]/4/5 leads to exocytosis (details at **Figure S20**).

We then tested the photopharmacology of **AzPico** in isolated primary chromaffin cells (the neuroendocrine cells in the adrenal gland that secrete adrenaline by exocytosis in response to electrical activation; these functionally express TRPC1/4/5 channels).^71^ 360 nm illumination of **AzPico** photoreversibly triggered robust inward currents that caused an increase in membrane capacitance (CM) in **wt** cells, indicative of exocytosis (**Figure 4b**). Neither response was observed in **145tko** cells, confirming the TRPC-specificity of this photopharmacology. By supporting that direct activation of endogenous TRPC channels in chromaffin cells can trigger exocytosis, this again suggests how **AzPico** may be used to exert functional control over endogenous biology: here, for optically-controlled release of adrenaline.

### TRPC4/5 photoswitches are effective in tissue slices and can reveal channel-specific biology (Figure 5)

We believed that **AzHC** and **AzPico**’s efficacy switch mechanism and high potency should make them effective photocontrol reagents for deeper tissues, and moved to test it. The hypothalamic arcuate nucleus (ARC; **Figure 5a**) is a signaling centre in the brain where most dopamine (Th+) neurons are known to express TRPC5, which contributes to spontaneous oscillatory burst-firing activity and Ca^2+^ burst responses,^34^ but also to sustained activation following stimulation with the hormone prolactin (a mechanism of reproductive signaling that has been conserved for >300 million years).^12,34^ The temporal distinction between these activation modes is striking, and suggests using either **AzHC** or **AzPico** as a time-resolved TRPC5 probe. However, we remain ignorant even of whether the most closely related congener TRPC4 plays a role in this circuitry, so we were particularly drawn to apply these reagents *comparatively*, hoping that their intersecting C4/C5 selectivities could deliver new information about TRPC4 biology in endogenous systems.

**Fig 5.**
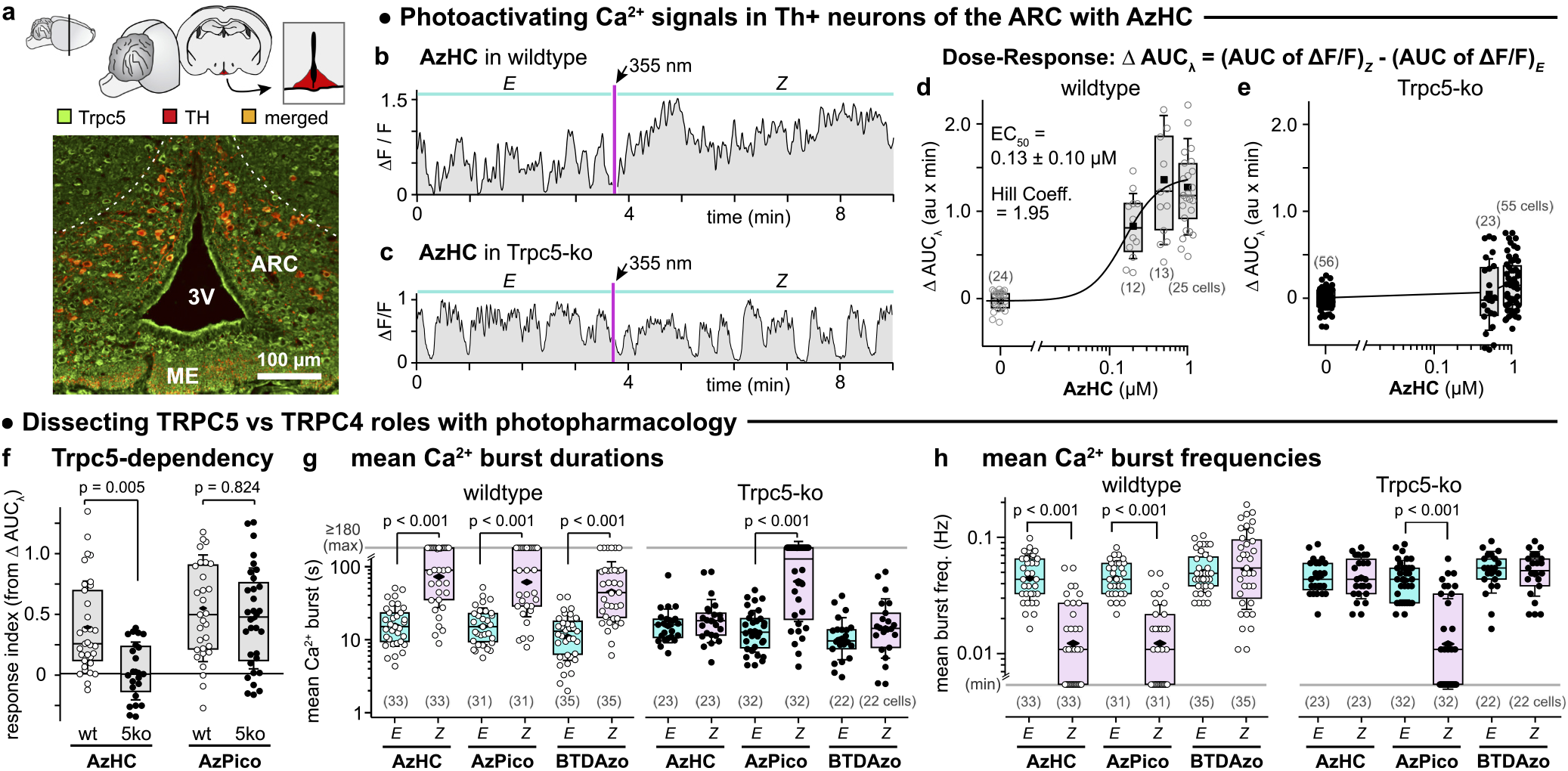
AzHC and AzPico are potent photoswitchable activators of TRPC-dependent Ca^2+^ responses in mouse hypothalamus. **a**, Coronal brain slice: cartoon with ARC region in red, and microscopy image with Trpc5 immunostained in green, Th+ neurons in red. **b-h**, Ca^2+^ responses in Th+ neurons of Th-GCaMP6f (wildtype or wt) or Th-GCaMP6f-?Trpc5 (Trpc5-ko or 5ko) mice. **b**,**c**, Single-cell Ca^2+^ traces before (“*E*”) and after (“*Z*”) a 355 nm pulse. **d**,**e**, ΔAUC_λ_: light-dependency of the area under the curves as acquired in (**b**,**c**). **f, AzHC** only photocontrols TRPC5-dependent Ca^2+^ responses; **AzPico** can photocontrol Ca^2+^ responses by another route (likely TRPC4). **g**,**h**, mean Ca^2+^ burst durations and frequencies in wt and 5ko upon *E→Z* photoswitching. (**AzHC**/**AzPico** at 500 nM except in dose-response; **BTDazo** at 10 µM; **d-h**: each point is 1 cell, (n) cells per group; box plots show interquartile ranges, median (line), mean (black rhombus), and SD whiskers; **f-h**: Kruskal-Wallis ANOVA, Dunn’s *p* values. For min/max values and all other details, see **Figure S21**; expanded legend at **Figure S27**.)

We therefore took 275 µm thick mouse brain slices through the dorsomedial ARC (**Figure 5a**), and imaged the Ca^2+^ indicator GCaMP6f in Th+ neurons while photoswitching **AzHC** or **AzPico**, to study TRPC[4]/5 photoactuation *in situ*. Ca^2+^ influx should be seen across several independent parameters: from time-resolved aspects such as longer Ca^2+^ burst durations and lower Ca^2+^ burst frequency, to simply larger areas under the curve of the fluorescent Ca^2+^ signal (AUC), which translate into a “response index” above 1 (details at **Figure S21**). Pleasingly, we again found that **AzHC** and **AzPico** can strongly photoswitch Ca^2+^ responses at endogenous TRPC levels by all these metrics. A single ≥23 ms pulse of 355 nm light (*E→Z*) induced dramatically sustained high-Ca^2+^ signals lasting up to ≥3 min (42% of cells; **Figure 5bdgh**). TRPC5 is sufficient for this signal, since TRPC5 knockout slices do not respond to **AzHC** photoswitching (**Figure 5ce**). However, *Z*-**AzPico** Ca^2+^ photoresponses are maintained despite TRPC5 knockout, providing the first functional evidence for a role of TRPC4 or TRPC1/C4 in the Ca^2+^ response in dorsomedial Th+ neurons of the ARC (**Figure 5f-h, Figure S21cd**), which can now be further explored^12^.

This discovery underscores the utility of this pair of photoswitches for deconvoluting the roles of TRPC4 from TRPC5. We also stress how important it was for these studies, that the reagents are high-potency as well as ideal-efficacy photoswitches. This allowed internally baselining signals after compound application (to overcome expression heterogeneity), then photoswitching activity on from zero background at a precisely defined time in a fully reproducible wavelength-dependent manner (all needed for reliable statistics). Temporally-modulated studies beyond the scope of this report are already underway, motivated by the xanthines’ sustained (low frequency) Ca^2+^ bursts, that contrast to the photoswitchable TRPC5 activator **BTDazo** (burst frequency barely affected, **Figure 5h**). This points to a rich interplay of pharmacology and spatiotemporally-resolved biology in complex tissues, that photoswitchable reagents are uniquely poised to tackle.

### Photoswitching tissue-level physiology: TRPC4-based photocontrol of intestinal motility (Figure 6)

After observing these ideal efficacy switch reagents photocontrol channel currents in thin endogenous tissue slices by microscopic imaging, we next aimed to test their capabilities by photoactuating macroscopic downstream processes in thick tissue sections. Landmark papers by Freichel^72^ and Zholos^16^ argued by irreversible suppression experiments that TRPC4 activation should be a critical component controlling peristalsis of the small intestine, occupying a downstream position relative to muscarinic acetylcholine receptors (mAChR, the target of atropine). Intestinal segment contractions are macroscopically coordinated oscillatory motions overlaid on a “tonic” baseline contractile force. A sub-threshold slow oscillatory pacemaker potential is amplified by mAChR via phospholipase C (PLC) and TRPC4 activation to surpass the threshold potential of voltage-gated Ca^2+^ channels, leading to peristaltic contractions (**Figure 6a**). We therefore expected that direct TRPC4 activation by *Z-***AzPico** might hijack the pacemaker signal to drive oscillatory contractility, even if upstream signaling by mAChR were blocked, and determined to test this and thus to directly elucidate the role of TRPC4 activation in intestinal contractility in endogenous tissues for the first time.

**Fig 6.**
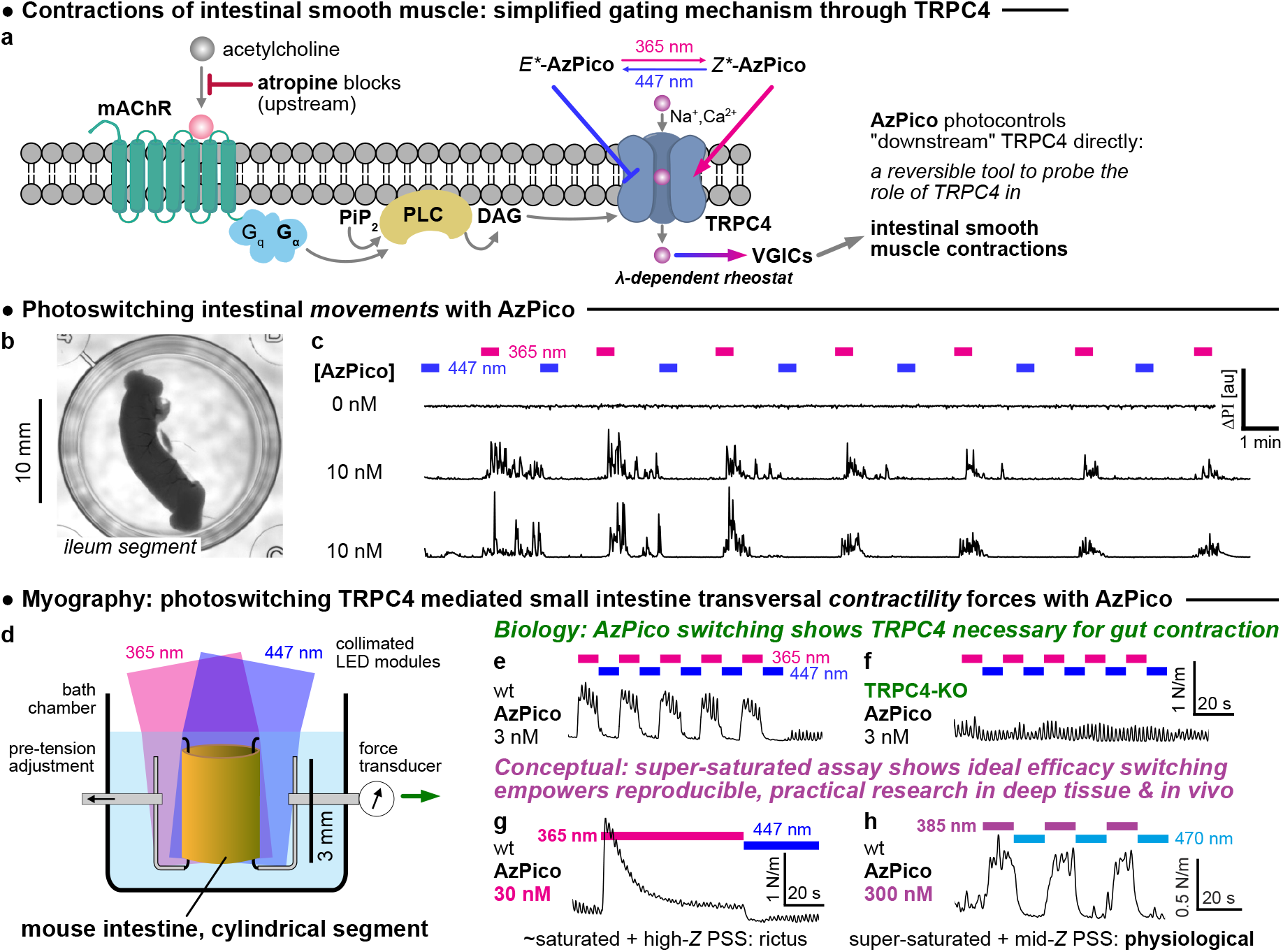
AzPico photocontrol of intestinal contractility shows the key role of TRPC4, and power of ideal efficacy switching. **a**, simplified molecular mechanism for TRPC4-dependent intestinal contractility^75^, that was now directly testable using **AzPico** (*E\**/*Z\** indicate *E/Z-*rich-PSSs). **b**,**c** (see also **Movie S1**), intestinal segments (mouse ileum) whose motility was blocked by atropine but were treated with **AzPico**, were driven into phases of fast macroscopic motions by UV photoswitching (longitudinal as well as ring muscle contractility), then returned to immobility by blue light, reversibly over many cycles. **d-f**, physiological-like ring contractility is reversibly stimulated and suppressed by alternating UV/blue illumination of segments treated with **AzPico**, in a TRPC4-dependent manner. **g**,**h**, the ideal efficacy switch paradigm allows fully reproducible control of deep tissue bioactivity, by leveraging the *saturation* of dose-and-photon-flux (hard to titrate) but the *selection* of wavelength (easy to choose) (see also **Figure S22**; full legend at **Figure S28**).

We took fresh 8-12 mm long segments of mouse small intestine (jejuneal and ileal), added low atropine concentrations (300 nM) to paralyse their spontaneous motility, and monitored their contractility by macroscopic video imaging while photoswitching **AzPico** (**Figure 6b**). Indeed, even with just 10 nM **AzPico**, UV illumination initiated vigorous motions, that were stopped rapidly by applying 447 nm light, then restarted with UV, and the on↔off photocontrol could be cycled many times (**Figure 6c, Movie S1**). No intestinal light responses were evident without **AzPico**; and, highlighting the tissue-specificity involved, **AzPico**-treated colon segments also had no photoresponse.

We next analysed intestinal contractility quantitatively, monitoring contractile muscle force in 3 mm long ileal or jejuneal segments by myography during **AzPico** photoswitching with UV and blue light (**Figure 6d**). With moderate **AzPico** concentrations (e.g. 3 nM), 365 nm UV not only increased tonic forces, but also doubled oscillatory forces (1.9×) yet *without* changing oscillatory frequency: indicating that *Z-***AzPico** “rides” the circuit driving typical contractility. Subsequent 447 nm blue light immediately reversed this and returned to basal tension; and 365/447 nm cycling could be repeated many times (**Figure 6e**). We assign this intestinal photocontrol to TRPC4 since (i) segments of TRPC4-deficient mice^73^ never responded to *E/Z-***AzPico** (**Figure 6f**), while (ii) tonic and oscillatory force photoresponses in TRPC5-deficient mice were indistinguishable from those in matched control mice with the same genetic background.

Our findings thus support that TRPC4 activation is not only necessary, but also sufficient, to convert sub-threshold oscillatory signals into effective peristaltic motility in the small intestine. On an applied level, we note that the **AzPico**-induced, optically tunable re-activation of atropine-paralysed intestinal segments also bears the potential of designing experimental therapies to treat medical conditions of insufficient intestinal motility, such as toxic or post-operative paralytic ileus. On a more conceptual level, we feel it is the full and rapid photoreversibility of both gross motility and myographic readouts, assessed on the tissue level in wildtype samples, which gives such high confidence of the molecular TRPC4-specificity of intestinal contractility. We stress however that this simple and direct result is *only* possible for the ideal efficacy photoswitch, whereas classical high-affinity compounds such as **(-)-Englerin A** or **Pico145** practically cannot be washed out, so internally-baselined, reversible experimentation without fatigue-based rundown or gross toxicity^74^ has been impossible before now. We also highlight that this is the first instance of a small molecule photoswitch being used to elucidate a target by hypothesis-driven research, instead of being created and used *after* target identification.

However, given the *in vivo* potential of ideal efficacy photoswitches, we are certain this will not be the last such case. To motivate this, we highlight a rare but practical example of true *light rather than dose control* that such photoswitches finally enable. Our protocols had converged to use 365 nm for high-completion *E→Z* switching and best channel activation; however, with moderate to high concentrations of **AzPico** (30 nM), this resulted in rictus-like intestinal tensioning and rundown, presumably via overstimulation (**Figure 6g**). On reflection, this seems to be a regular biological situation: that a desired, physiological effect is only reached when a target protein is *partially but not fully stimulated*. This could be avoided in a classical way by titrating down the **AzPico** concentration to 3 nM (**Supporting Section 7.1.1**): but the power of the efficacy switch mechanism allows a much simpler workaround. To stringently test the ideal model shown in **Figure 1g-i**, we now fully saturated tissues using *300 nM* **AzPico**, but adopted 385 nm instead for UV illuminations. This 20 nm wavelength shift perfectly reproducibly drove only partial channel activation (even under saturating illumination), and so returned the physiology-like oscillatory intestinal contractions we sought, by a fully reproducible *doubly saturating* compound+light treatment protocol (**Figure 6h**). No affinity photoswitch can work when both light and compound dosage are maximised: at least one if not both of them are always limiting for biological performance. Now, we show that adopting this paradigm shift to *ideal efficacy photoswitching* can empower deep tissue or *in vivo* research, while overcoming the traditionally assumed irreproducibility of compound-and-light-dosing. Demonstrating this even on TRPC4/5 as non-linearly-responsive targets with time-and-dose-dependent bio-activity, gives an additional impression of the power that ideal efficacy photoswitching can bring to biology: whether on simple spatiotemporally-differentiated targets, or (here) as integrated in increasingly complex circuitries of biological sensing, signaling, and control.

### Summary

We rationally developed **AzHC** and **AzPico** as ideal efficacy photoswitches for TRPC[4]/5. Their photoisomerisation flips them between *E* inverse agonists and *Z* agonists, with both forms having excellent binding affinity: resulting in a pair of lit-active low-nanomolar-potency tools that can be used together to elucidate TRPC5-selective or TRPC4-selective biology. To date, there have only been two reports of protein-ligand structures where both *E-* and *Z-*isomers of the same azobenzene reagent were bound;^49,60^ we now report two more, with both TRPC4 and TRPC5 *E/Z* structure pairs, which will help progress design rules for efficacy switches on other targets. Unlike nearly all photopharmaceuticals, the bioactivity of **AzHC**/**AzPico** is fully determined by the illumination wavelength used, not their concentration. This suits them to reliable and reproducible operation across diverse model systems, from over-expression in HEK cells, to endogenous expression in primary neurons or primary adrenalin-secreting cells, to mouse brain slices, and finally intact segments of intestine, where they elegantly and directly substantiated a long-unsolved hypothesis for TRPC4-dependent contractility.

Importantly for the TRPC field, the demonstrations of highly effective *tissue-level photoresponse* under exposure to low concentrations of **AzPico** / **AzHC** mark these reagents as exceptionally effective optical tools for precisely and reversibly manipulating endogenous TRPC-dependent^17^ biology *in situ*. Their likely applications thus stretch far beyond the muscular and neuronal applications shown here, towards elucidating TRPC4/5’s rich and largely still cryptic biology. Finally, the rapid macroscopic photoswitchability of **AzPico**’s downstream secondary and tertiary effects (i.e. not only channel opening and ion flux, but also of the integration and control of the native cascade which ultimately controls tissue-scale muscle movement) is highly unusual, if not unique, in photopharmacology (**Figure 6**).

For biology as well as chemistry in general, we have thus shown that the paradigm of efficacy photoswitching can be rationally chemically designed as well as rationally biologically exploited, and that it at last achieves a >20-year-old dream of photopharmacology: delivering robust pharmacological control of endogenous systems, that is *purely* and effectively controlled by light without needing to bow to the vagaries of drug dosing and distribution. This is the feature combination that suits it so excellently to the challenges of deep tissue and *in vivo* work. We have introduced some general biological target considerations as well as general chemical design concepts to ground the widespread introduction of efficacy photoswitching.^57^ In brief, our perception has been that the targets which are of the highest possible *importance* for studying *in vivo* with the resolution that efficacy switches allow, are also the targets which are the most *likely* to be addressable by this spatiotemporally precise paradigm. We refer to these as “poised targets”, such as ion channels, receptors, sensory or signaling integrators, etc: covering many of the proteins that need to rapidly and spatiotemporally modulate their functions, in order to support life. That match of need to opportunity is of course not accidental; it simply reflects how biology has evolved and perfected its own ligand-(or protein-)based regulatory mechanisms to operate in complex environments.

We particularly encourage chemists to take up an awareness of this possibility, to start rigorously testing for it even where it was not a design aim,^52,53,55,56^ and to work towards rationally using the efficacy photoswitching paradigm to generate a cornucopia of *in vivo-*competent reagents, rather than remaining locked to affinity photoswitching approaches. We foresee that such efficacy photoswitches can unlock a new era for photocontrolled biology, in ways that deeply impact not only chemical or cell culture proof of principle studies, but translate seamlessly across to basic physiology and medical research, to noninvasively probe and modulate endogenous pathways in deep tissues and *in vivo*.

## Caption to Movie S1

**Movie S1:** Photocontrol over gross motility of intestinal segments by alternating 365/447 nm illumination in the presence of 10 nM **AzPico**.

## Author Contributions

As related to **Figure 1a-j, 2h-j**: M.Müller designed targets and performed synthesis and photochemical evaluation, as supervised by O.T.-S.; M.Müller and O.T.-S. performed mechanistic analysis and modelling.

As related to **Figures 1k-n, 2a-g, 6**: K.N. and F.B. performed all experiments as designed and supervised by N.U. and M.S.. M.S. analysed data and drafted the corresponding manuscript sections and figures.

As related to **Figure 3a-d**: S.A.P. performed protein expression/purification, cryoEM data collection, processing, and model building and data analysis; C.C.B. designed and cloned TRPC4 and TRPC5 plasmids; A.V.J. performed intracellular calcium recordings and analysed data; S.A.P. and A.V.J. made figures; S.P.M and R.S.B designed and supervised experiments and obtained funding; R.S.B. wrote sections of the manuscript with input from S.A.P., C.C.B, A.V.J. and S.P.M..As related to **Figure 3e-h**: D.V. performed protein expression/purification, cryoEM data collection and processing, under the supervision of S.R.

As related to **Figure 4**: K.O. and M.Makke performed all experiments related to chromaffin cells and hippocampal neurons, that were designed, supervised, and analysed by Y.S. and D.B., who drafted the corresponding manuscript sections and figures.

As related to **Figure 5**: N.K.O., F.Z., and T.L.Z. designed, performed, analysed, and drafted the manuscript for all experiments related to mouse hypothalamus.

M.S. and O.T.-S. designed the study.

M.Müller and O.T.-S. performed overall data and figure assembly, and wrote the manuscript with input from all authors.

## Acknowledgements

M.Müller thanks the Joachim Herz Foundation for fellowship support. A.V.J. thanks the BBSRC for a White Rose DTP PhD studentship. Professor Eric Gouaux (Vollum Institute) is kindly acknowledged for providing the BacMam vector. Marc Freichel (Heidelberg) is kindly acknowledged for providing the TRPC4- and TRPC5-KO mice. O.T.-S. thanks Rob Leurs, Michael Decker, Pau Gorostiza, Wiktor Szymanski, and James Frank, for feedback on the prior reports of efficacy switches.

## Data and materials availability

All data needed to evaluate the conclusions in the paper are present in the paper and/or the Supplementary Materials. Data are also deposited and freely available on BioRxiv and Figshare (*dois upon paper acceptance*).

Cryo-EM structures are deposited as:

TRPC4_DR_:*E*-**AzPico**, 3.0 Å, PDB 9FXL, EMDB 50850; TRPC4_DR_:*Z*-**AzPico**, 3.1 Å, PDB 9FXM, EMDB 50851; hTRPC5:*E*-**AzHC**, 2.6 Å, PDB xxxx, EMDB xxxx; hTRPC5:*Z*-**AzHC**, 2.9 Å, PDB xxxx, EMDB xxxx.

