## Supporting Information for "Ideal efficacy photoswitches for TRPC4/5 channels harness high potency for spatiotemporally-resolved control of TRPC function in live tissues"

### 1 Supporting Notes

#### 1.1 Supporting Note 1: Pharmacology for Efficacy Switching

Xanthines **Pico145** and **HC-070**[1] are low nanomolar/picomolar potency inhibitors of homomeric and heteromeric TRPC1/4/5 channels.[2,3] The very similar **AM237**[1] (single-atom replacement from **Pico145**: chlorine instead of hydrogen at C'4) is instead a nanomolar partial *agonist* of homomeric TRPC5, despite being a nanomolar *inhibitor* of the full agonist (-)-englerin A (**EA)** in the context of homomeric TRPC4 and heteromeric TRPC1/5 and TRPC1/4 (structures: **Figure S1**; potencies: **Table S1**).[4] Structures of TRPC5 in complex with **Pico145** and **HC-070** were recently obtained, showing their conserved binding site at the transmembrane domain between two monomers.[5,6] A docking study of **AM237** into the structure of **Pico145** bound to TRPC5 indicated that avoiding a steric clash of the chlorine with a single amino acid residue of the protein requires rearranging the structure (either the protein or the ligand) resulting in a plausible mechanism for the switch from inhibitory **Pico145** to agonistic **AM237**. A similarly dramatic efficacy change resulting from a tiny structural modification was also observed for **E54**, formally a deoxygenated ether derivative of **EA**, which also antagonises **EA**'s activation of homomeric TRPC4 (weakly), and homomeric TRPC5 and heteromeric TRPC1:4 (strongly), presumably by competitive binding with suppressed channel activation[4].

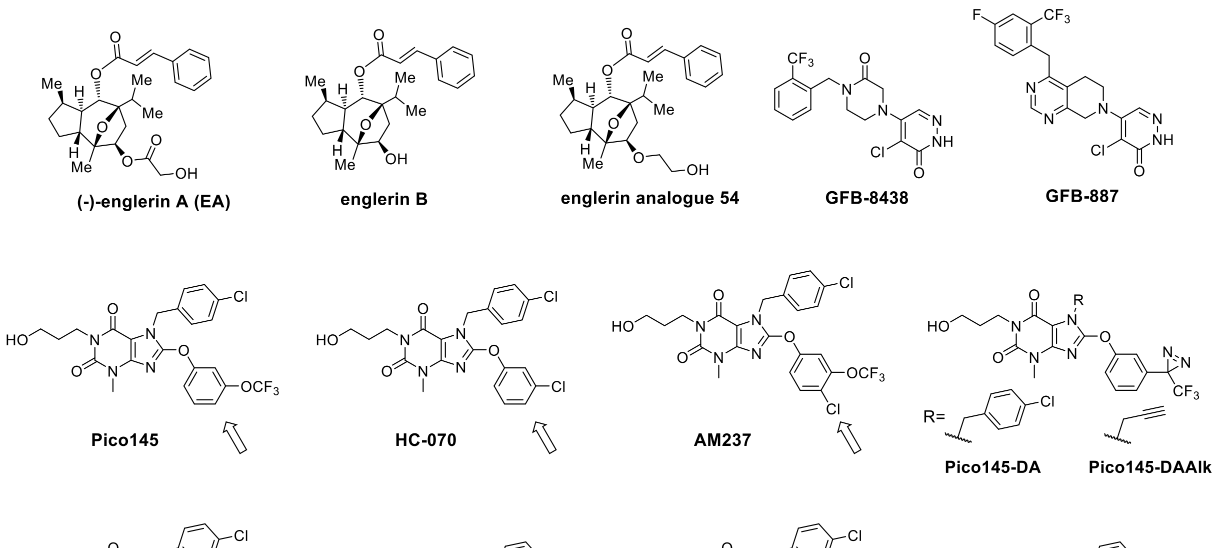

**Figure S1.** Structures of lead compounds incl. **Pico145**, **HC-070**, **AM237** (xanthines), xanthine derived photoaffinity probes **Pico145-DA** and **Pico145-DAAlk**, and **(-)-englerin A** and **E54** (englerins supposedly share the same binding site with xanthines), as well as pyridazinones **GFB-8438** and **GFB887** (different binding site); alongside this work's star photoswitchable modulators **AzPico** and **AzHC**.

For interpreting the $EC_{50}$s reported in **Fig 1n** and **Table S1**, two notes should be kept in mind: (1) **comparing $EC_{50}$/$IC_{50}$s for agonist *Z* and antagonist *E* does not deliver any information about relative binding affinity,** since (a) the experiment types are so different in nature (agonists: sole-ligand assay; antagonists: out-competing 10-30 nM of the high-affinity **EA**), and (b) there are many additional unknown (and complicating) factors related to the multiple binding sites per channel, including potential binding cooperativities. A discussion of these effects is beyond the scope of this paper; but it is plausible that the binding affinity of *E*-**AzHC** and *E*-**AzPico** isomers is similar to that of the ca. 1 nM potent **EA**, given that their $EC_{50}$s for antagonism in the competition

assay are only ca. 3-5 times the applied concentration of **EA**. This is a handwaving explanation, but it can justify why the affinity of the *E/Z*-isomers may be similar.
(2) The $EC_{50}$ values are reported as the midpoints of *unconstrained* sigmoidal fits to the FLIPR data (**Fig 1l** and **Figure S2**): with different levels of channel activation / blocking and very different Hill slopes across different compounds, isomers, and the two channels. This argues that **the midpoint values only capture one of the *several* important factors describing the compounds' activity**.

We believe that it is possible to derive indicative relative values for binding affinity comparing the *E* and *Z* isomers of the same compound using only the combination of FLIPR and ephys data as reported in this paper; this analysis is being developed in a separate paper (see also below).

**Table S1.** Potencies on TRPC4/5 depicted as either simple agonistic potency (blue) or else inhibitory potency against activation by 10-30 nM **EA** (orange). *Z**-values refer to mostly-*Z*-isomer reached at PSS at 365 nm, *E**-values refer to experiments that initially applied 100% *E*-isomer (thermally relaxed), though in both cases the $Ca^{2+}$ imaging in FLIPR with Fluo4-AM uses 470 nm excitation, which may over time promote photoswitching towards a PSS that is intermediate between those of pure-365 nm and all-*E*; this is controlled for in the more reliable ephys measurements). *(Table indices: a, data from ref[7]; b, data from ref[2]; c, data from ref[3]; d, data from ref[8].)*

<table>
<tr><th></th><th colspan="2">$EC_{50}$ [nM] (for simple agonism or EA-competitive inhibition)</th></tr>
<tr><td></td><td>TRPC4</td><td>TRPC5</td></tr>
<tr><td>EA</td><td>11 ag.[a]</td><td>7.6 ag [a]</td></tr>
<tr><td>Pico145</td><td>0.35 (vs 10 nM EA) [b]</td><td>1.3 (vs 10 nM EA) [b]</td></tr>
<tr><td>HC-070</td><td>0.5 (vs 10 µM carbachol) [c]</td><td>2.0 (vs 20 µM carbachol) [c]</td></tr>
<tr><td>AM237</td><td>7 (vs 30 nM EA) [d]</td><td>20 ag [d] | 13 (vs 30 nM EA) [d]</td></tr>
<tr><td>E-AzPico</td><td>90 (vs 30 nM EA)</td><td>67 (vs 30 nM EA)</td></tr>
<tr><td>Z*-AzPico</td><td>3.0 ag.</td><td>4.0 ag.</td></tr>
<tr><td>E-AzHC</td><td>110 (vs 30 nM EA)</td><td>149 (vs 30 nM EA)</td></tr>
<tr><td>Z*-AzHC</td><td>>1000</td><td>6.4 ag.</td></tr>
<tr><td></td><td>orange: inhibitory effect</td><td>ag.: agonistic effect</td></tr>
</table>

##### 1.1.1 From always-active antagonists and agonists towards efficacy photoswitches

In this paper, we will show that this *"chemical efficacy switch"* behaviour can be mimicked by photoisomerising an azobenzene that is merged in at the C'4 position yielding the compounds **AzPico** and **AzHC** which show a strong lit active agonistic effect for the *Z*-isomers and a strong inhibitory effect for the *E*-isomers (**Table S1**). Essentially, the effects of **Pico145** and **AM237** are latent within a single (two-state reversibly photoswitchable) molecule. This combination is what we call an *efficacy switch* (also see main text **Figure 1e-j**).

##### 1.1.2 Mechanistic test for efficacy photoswitching

The concentration-independent-but-PSS-determined channel activation that **AzHC** and **AzHC** deliver (**Figures 1-2**) is a powerful support for an efficacy switch interpretation. To indirectly support this further, we also measured the effects of *E*-**AzPico** and *E*-**AzHC** against **EA**-activated TRPC4/5. Both *E*-isomers bind to TRPC4/5 and inhibit **EA**-evoked currents with nanomolar potencies (**Figure S2**, **Table S2**). This supports the hypothesis that *E*-**AzHC/AzPico** are **EA**-competitive tight binders on both channels (much as **Pico145** was proposed to be an **EA-**competitive binder[5,9,10]), so their efficacy as single agent photoswitches results from E/Z-competitive binding that is determined by their PSS.

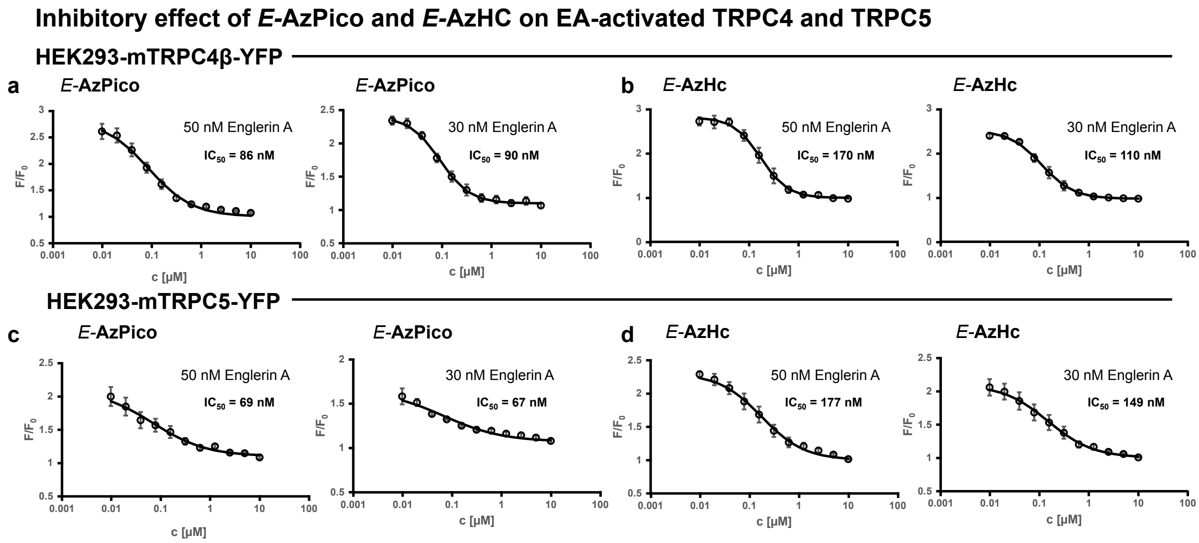

**Figure S2.** Dose response curves for *E*-**AzPico** and *E*-**AzHC** silencing the activation of TRPC4 and TRPC5 that is driven by 30-50 nM of **EA**. **(a-b)** 3 µM Fluo-4-loaded HEK293-mTRPC4β-YFP; **(c-d)** 3 µM Fluo-4-loaded HEK293-mTRPC5-YFP. **(a)** *E*-**AzPico** TRPC4 activated by 30 nM or 50 nM **EA**. **(b)** *E*-**AzHC** TRPC4 activated by 30 nM or 50 nM **EA**. **(c)** *E*-**AzPico** TRPC4 activated by 30 nM or 50 nM **EA**. **(d)** *E*-**AzHC** TRPC4 activated by 30 nM or 50 nM **EA**.

**Table S2.** Potencies on TRPC4/5 depicted as inhibitory against **EA** activation. ($^a$ Inhibitory effect measured against 30 nM **EA**; $^b$ Inhibitory effect measured against 50 nM **EA**).

| | $IC_{50}$ TRPC4 [nM] | $IC_{50}$ TRPC5 [nM] |
|---|---|---|
| *E*-**AzPico** | 90 $^a$ / 86 $^b$ | 110 $^a$ / 170 $^b$ |
| *E*-**AzHC** | 67 $^a$ / 69 $^b$ | 149 $^a$ / 177 $^b$ |

#### 1.2 Supporting Note 2: Prior Art in Efficacy Photoswitching

##### 1.2.1 Reported Prior Art in Efficacy Photoswitching

In general, ligand development with confirmed efficacy switching has been rather thinly represented in the literature; and there are several examples of claimed efficacy switches that do not hold up to scrutiny, or which are not as useful as might be hoped (small relative changes in activity without a change in the sign of activity; and/or low potency; and/or were not photoisomerised or perhaps not usefully photoisomerisable *in situ*; etc). We recommend Leurs' 2018 Angewandte[11] and particularly Leurs' 2019 Beilstein JOC[12] as being the clearest demonstrations and discussions of efficacy switches that were well-characterised and were used as photoswitches. Leurs' later review article[13] also explicitly contains some literature statistics on efficacy switches (see that paper's Figure 7f), and Szymanski and Feringa's review[14] collected and commented the references to the efficacy switches that we have also cited in this work.[12,15–23]

While these precedenting cases have been published, we believe that our discussion goes beyond those reports in the sense of interpreting efficacy switching as a systematic feature that is fundamentally different and advantageous for photopharmacology: hence our explicit call to develop high-potency competitive binding efficacy switches *as a means to succeed in reproducible photoswitching despite variable biodistribution* (expected *in vivo* and in deep tissue) *by fixing selected wavelengths to drive fixed activity profiles by fixed PSS ratios* (the opposite of typical "chromodosing" wherein both wavelength and applied concentration are tuned to drive fixed activity profiles by fixed concentration of an active photoisomer).

##### 1.2.2 Unsuspected Prior Art in (Ideal) Efficacy Photoswitching

Probably very many more compounds that have been reported without explicit commentary, or even reported implicitly as though they were affinity switches, actually *are* efficacy photoswitches, just that they have not been tested or proven to act as such; and of these, we believe a significant number are "ideal" in the sense of near-identical *E/Z*-affinities. One hallmark we expect in such cases is that the wavelength-dose-response profiles should plateau, at different heights, under different wavelengths: as depicted in **Figure 1l** for non-ideal-wavelengths (non-ideal-efficacy-switches will generate similar plots), or **Figure 1g** for ideal wavelengths on an ideal efficacy switch.

Such plot shapes *can* however be confused with drugs whose solubility reaches a limit beyond which no further increase in concentration-dependent bioactivity is possible (plateau), if only two wavelengths are used for activity determination.

Our suggestion for unambiguous identification and interpretation of efficacy switches is thus to plot action spectrum based data instead, at multiple concentrations if needed (**Figure 2h-j**). Form fitting to log([*Z*]) or log([*E*]) traces derived from PSS values (in an appropriate solvent!) will identify affinity switches (**Figure 2ij**), whereas form fitting to log([*Z/E*]) traces marks efficacy switches (**Figure 2h**).

We have re-parsed some of our favourite photopharmacology literature to search for unsuspected ~ideal efficacy switching. A more comprehensive theoretical treatment that traces the target-based and pharmacology-based *drivers* of efficacy switching is in preparation as a separate paper; below we list just some of the reported examples that to us seem very likely to have been ideal efficacy switches. We note that several of these were translated from cell culture into complex settings and even *in vivo* photoswitching (e.g. Trauner 2019[24], Fuchter 2020[25]) with success that would have been unexplainable for an affinity switch. <u>For the photopharmacology field beyond TRP channels, this is probably the key message of our paper: the rewards and importance of designing and recognising ideal efficacy switches, in order to optimise and exploit them as such</u>.

***Five of the more recent examples are:***

##### 1.2.3 Tangential Remarks & Outlook for Efficacy Photoswitches

(1) In the main text, we argued that the major reason why affinity switches' applicability for bidirectional photocontrol has been mainly limited to highly controlled cell culture settings is that the bioactivity applied by an affinity switch is so sensitive to its concentration (**Figure 1j**), yet linearly-reproducible dose-fixing is problematic or impossible for deep tissue or *in vivo* work. We note though, that if only unidirectional photo-turn-on is used ("one-shot"), this limitation diminishes somewhat since light dose can be titrated at a fixed wavelength. Then again, such cases also do not require a *photoswitch* per se, and can be (probably better achieved) by photocages.

(2) In addition to the features outlined in the main text, we suggest that another special feature of a high-potency efficacy photoswitch with near-equal-affinity isomers is that, when binding site saturation is ensured, the PSS dependence of activity can give insights into cooperativity that would not otherwise be accessible with always-active ligand/s. For example, at 400 nm the PSS *E:Z* ratio is ca. 3:1 (**Table S1**), and there is no detectable channel activation (**Figure 2eg**). Of course, even at full saturation, there will always be a statistical distribution of $E_n/Z_{4-n}$:TRP complexes (and even that C2-symmetric $E_2/Z_2$:TRP may have a different activity profile than its C1-symmetric $E_2/Z_2$:TRP isomer): nonetheless, this finding is quite suggestive that occupying just one of the four TRPC4/5 binding sites with the xanthine agonist (*Z*-**AzPico**) is insufficient for channel activity, if the other three sites are occupied by an antagonist (*E*-**AzPico**; further discussion at **Figure S14**).

(3) As hinted in several locations, it seems clear to us that there are many levels of target-based features which can be used to pre-select potential targets for developing efficacy photoswitches even when no ligand SAR data is known that would predict an efficacy cliff. The most significant level, in our opinion, is that "**poised targets**" that natively must be able to swap back and forth between (at least) two metastable states to balance and deliver their physiological function, such as ligand-gated channels and receptors, seem ideal candidates for efficacy photoswitch operation in general.[29] This as well as the other levels of target selection, as well as the general features and consequences of efficacy photoswitching for chemists and for biologists, will be addressed in more detail in our upcoming conceptual Perspective article that accompanies and comments on this paper's experimental results.

#### Supporting Note 3: Results from Other Compounds in our Panel

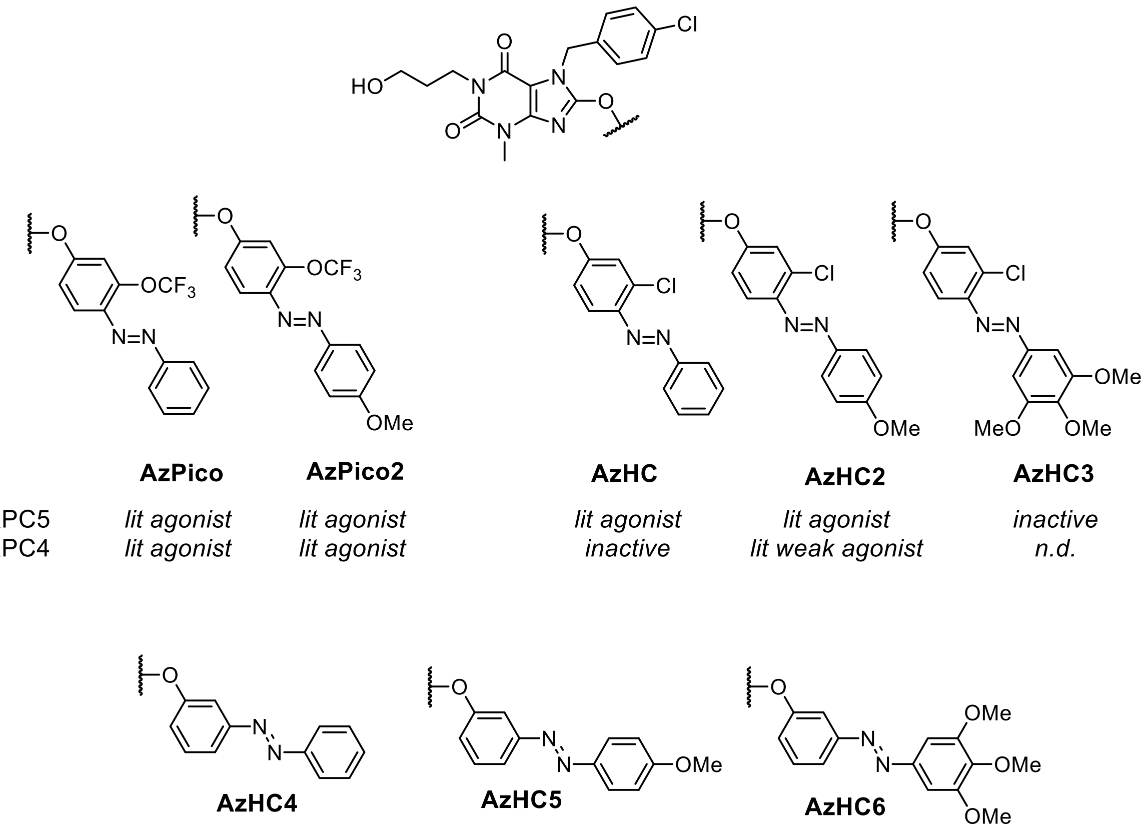

| | AzPico | AzPico2 | AzHC | AzHC2 | AzHC3 |
|---|---|---|---|---|---|
| TRPC5 | *lit agonist* | *lit agonist* | *lit agonist* | *lit agonist* | *inactive* |
| TRPC4 | *lit agonist* | *lit agonist* | *inactive* | *lit weak agonist* | *n.d.* |

| | AzHC4 | AzHC5 | AzHC6 |
|---|---|---|---|
| TRPC5 | *inactive* | *inactive* | *lit weak agonist* |
| TRPC4 | *n.d.* | *n.d.* | *n.d.* |

**Figure S3.** structures of all the compounds made and tested in this work; and short summary of biological activity that guided the selection of **AzHC** and **AzPico** as the best candidates to progress.

The initial FLIPR screening was carried out with all compounds shown in **Figure S3** on TRPC5; only hits **AzPico**, **AzPico2**, **AzHC** and **AzHC2** were further analysed on TRPC4. It is interesting to note that the structural changes which ruined agonistic potency in the abandoned compounds are comparatively small, e.g. 3,4,5-trimethoxy for **AzHC3**, a motif that tends to improve photoswitching completion and replacement of the C'3 residue with a photoswitch did not result in any biological relevant photoswitching performance and those compounds were not elucidated further. Among the hit compounds, **AzPico2** and **AzHC2** were discontinued even though their para-methoxy increases *E*→*Z* photoswitching completion (**Figure S9**), because their *Z*→*E* switching was less complete, their solubility was worse, and their total $Ca^{2+}$ signal was lower, as compared to **AzPico** and **AzHC** respectively. We thus chose to investigate both **AzPico** and **AzHC** because of their different selectivity profile on TRPC4/5 (continued in **Supporting Note 4**).

#### 1.3 Supporting Note 4: Structural Biology and TRP Selectivity

We prepared and acquired TRPC5:***E/Z*-AzHC** samples in one lab, and TRPC4:***E/Z*-AzPico** independently in another, which was useful for several plausibility checks. (1) There are additional lipid densities in the pockets that are present in both TRPC4 and TRPC5, and the changes to the lipid densities in the EM maps between *E* and *Z* isomers are similar in TRPC4 and TRPC5 as well: so these changes are not dependent on protein, nature of amphipol used, or lab in which the data were obtained/processed, but are rather inherent features of the system. (2) The positions of the distal rings which are much less-well-resolved than the rest of the ligand, were independently best aligned for the *E* and *Z* isomers, and in each case found to be *Z*-buried vs. *E*-projected, which gives more confidence in this structure-based analysis. (3) Identical deleterious effects of DTT were found in each lab, and removing it removed those problems.

##### 1.3.1 Detailed Commentary: hTRPC5:*E/Z*-AzHC

To start, we note that although no channel-open structures are known, and all reported ligand structures are with inhibitors, there has been docking work done on xanthine-based TRPC5 *activators* such as **AM237**, **Pico145-DA**, and **Pico145-DAAlk**, (**Figures S1**). Docking suggested that a slight twist of their *C*-8 aryloxy substituents, compared to those of the inhibitors **Pico145** and **HC-070**, is needed to avoid potential clashes with nearby residues V610, V579 and L521.[8,30] Because the functional data suggest **AzHC** is a TRPC5 efficacy switch (*E*-antagonist / *Z*-agonist), it would be expected that the *E/Z*-isomers both bind to TRPC5 but in different ways. To study TRPC5 complexes with both **AzHC** isomers by cryo-EM, we made several changes to our previously published cryo-EM pipeline[31] (see **Supporting Section 4**). In brief, **AzHC** samples were irradiated with blue light (440 nm; to access **(*E*)-AzHC**) or UV-A light (365 nm; to access **(*Z*)-AzHC**) prior to incubation with purified MBP-hTRPC5$_{\Delta 766-975}$ and during grid preparation; at all other times, samples were handled under red light (650 nm) to minimise photoisomerisation. We found that **AzHC** degrades in DTT-containing buffers, as confirmed by HPLC analysis; and solving initial "*E/Z*-**AzHC**:TRPC5" cryo-EM structures obtained with DTT-containing buffers either did not locate the terminal phenyl ring of the azobenzene and could have been assigned to reductive N=N bond cleavage, or else indicated an unusual ca. 120° CNNC torsion angle that could have been assigned to the diphenylhydrazine; we therefore discontinued analysis of these data sets and DTT was omitted from buffers used to prepare all remaining **TRPC5:AzHC** (and TRPC4:AzPico) samples.

We determined high-resolution structures of **hTRPC5:(*E*)-AzHC** (2.60 Å) and **hTRPC5:(*Z*)-AzHC** (2.90 Å) without applying symmetry (**Figure 3**; **Figures S4-S5 and S15-S18**; **Table S4**).

In **Figure 3ab**, TRPC5 is bound to *E*-**AzHC** (resolved to 2.60 Å in C1 symmetry); 4 **AzHC** molecules are bound and the best-resolved binding site is shown. There is high confidence in the binding pose (there are slightly different solutions for the *E*-azobenzene in the 4 sites but no major differences); and there is some lipid density still present in the map (likely: displaced phospholipid). In **Figure 3cd**, TRPC5 is bound to *Z*-**AzHC** (resolved to 2.90 Å in C1 symmetry); 4 **AzHC** molecules are bound, the best resolved is shown; EM map suggests flexibility of the *Z*-azobenzene as multiple poses could be docked by Glide-EM; lipid density is still present in map, but overlaps partly with the compound density (probably due to multiple conformations in the binding pocket in data set). Applying C4 symmetry led to higher overall resolution, but less well-defined densities for the azobenzene, potentially because of incomplete photoswitching of **AzHC** during sample preparation and/or incomplete displacement of the resident lipid from the binding sites. The **AzHC** *E* and *Z* isomers could be built into the lipid/xanthine binding site, with near-identical positions to **Pico145**[31] for the xanthine core, the 3-hydroxypropyl on *N*-1, and the 4-chlorobenzyl on *N*-7 (**Figure 3**). Clear differences between the **AzHC** *E* and *Z* isomers were seen for the azobenzene distal ring: in **(*E*)-AzHC** it makes a π-π interaction with Phe522, while in **(*Z*)-AzHC,** it makes a π-π interaction instead with Tyr524 (**Figure 3**; **Figure S4a**). Considering the protein, most residues in the ligand binding site are in similar positions in both structures, except for Phe520, which flips strongly when the agonistic *Z*-**AzHC** is bound (**Figure S4b**). These data further support the finding that **AzHC** acts as an efficacy switch and provide the first structural insights into how closely related xanthines can have opposite effects on TRPC5 function (i.e. inhibition vs. activation).

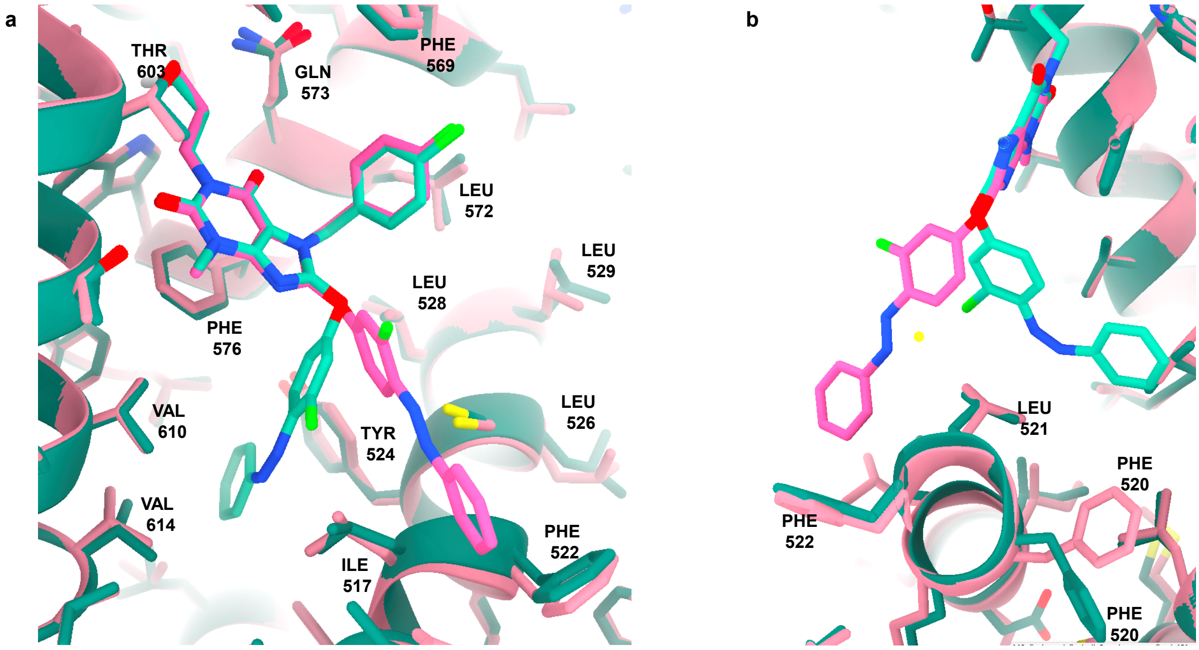

**Figure S4**. **Comparison of hTRPC5 binding modes of the AzHC isomers. (a)** The azobenzene distal phenyl ring interacts with Phe522 for **(*E*)-AzHC** (magenta ligand and protein), but with Tyr524 for **(*Z*)-AzHC** (teal ligand and protein). **(b)** *E/Z*-Photoisomers of **AzHC** feature a major difference in the conformation of Phe520 (at lower right).

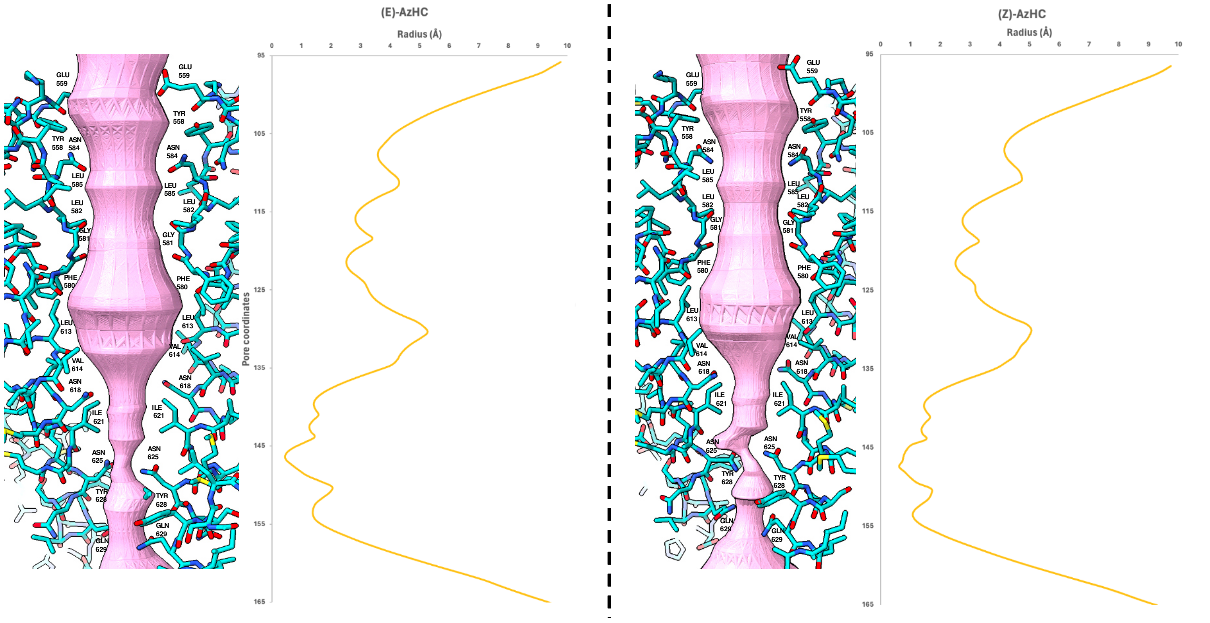

**Figure S5**. **Pore analysis of TRPC5 structures.** Asymmetric pore analysis of **TRPC5:(*E*)-AzHC** and **TRPC5:(*Z*)-AzHC** structures with PoreAnalyser[32] (default settings) suggests that both structures are in a closed state.

Acquiring open-pore TRP channel structures is highly challenging. PoreAnalyser[32] analysis suggests that both *E* and *Z* structures are channel-closed (**Figure S5**), which we believe is due to the channel's restraint in the amphipol stabiliser though it could also be due to low open state probability (especially without membrane potential), or the absence of cellular factors needed for opening (e.g., the membrane environment, $Ca^{2+}$, bound lipids, post-translational modifications, protein-protein interactions); incomplete switching in all binding sites or incomplete displacement of lipid might also contribute. Still, while needing cautious interpretation, the structural rearrangements upon binding of agonist ***Z*-AzHC** in the closed form may still be usefully indicative.

##### 1.3.2 Detailed Commentary: TRPC4$_{DR}$:*E/Z*-AzPico

We likewise determined the structure of DR-TRPC4:(*E*)-**AzPico** (3.0 Å) and DR-TRPC4:(*Z*)-**AzPico** (3.1 Å) without applying symmetry. The high-resolution structures enabled us to model the ligand into the density that was distinct from the annular lipid density reported from our earlier studies.[33,34] We could model the ligand xanthine core with high confidence which is similar to the previously reported structure of Pico145 bound with TRPC5.[5]

The difference in the *cis* (*Z*) and *trans* (*E*) isomers of the **AzPico** could be modelled after processing the data with C1 symmetry that gave a slightly better density for the less-resolved terminal benzene ring. The relative displacement of the stabilising residues Leu517, Leu520 and Phe521 was visible when comparing the *E*-**AzPico** and *Z*-**AzPico** densities, boosting our confidence in ligand modelling (**Figure S6**). The terminal benzene ring makes a π-π interaction with Phe521 in *E*-**AzPico** while in *Z*-**AzPico** the terminal ring flips to make a π-π interaction with Tyr523.

Surprisingly, the *Z*-**AzPico** that activates the channel, yielded a closed state (**Figure S7**) which could be attributed to the low open state probability of the channel or the absence of other cellular factors required for opening the channel as mimicked by TRPC5. Nevertheless, the *Z*-bound structure gives a plausible explanation for channel activation, in that the terminal benzene ring of *Z*-**AzPico** interacts with S6 helix residues closer to channel gating residues that might influence opening, whereas in the *E* isomer the terminal ring interacts with the S5 helix (**Figures S6-7**).

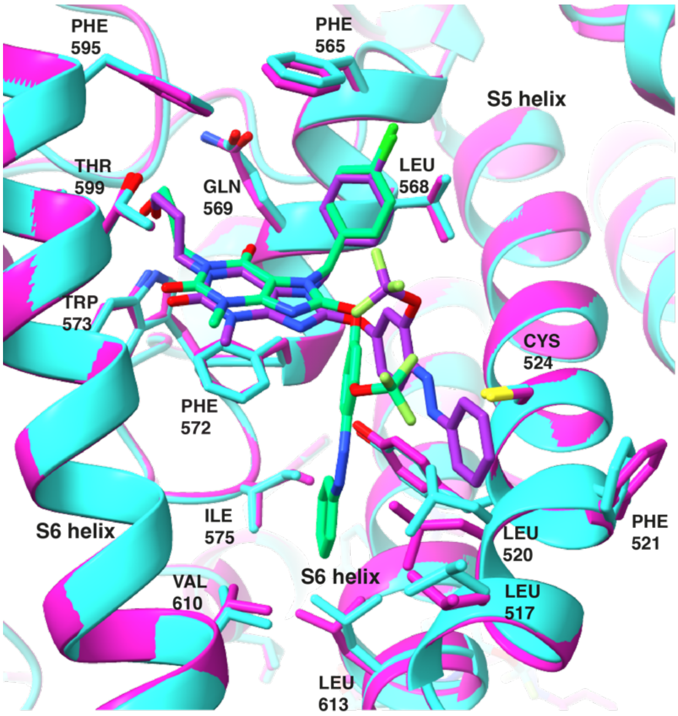

**Figure S6**. Comparison of the binding modes of *E/Z*-**AzPico** isomers to TRPC4. *Z*-**AzPico** bound TRPC4 is shown in cyan with the ligand in green. *E*-**AzPico** bound TRPC4 is shown in magenta with the ligand in dark purple. The residues interacting with the ligand are shown with sticks and labelled. The residues interacting with the xanthine core of the ligand show less relative displacement between *E* and *Z* structures than those interacting with the terminal benzene ring. The S6 helix involved in channel gating is labelled.

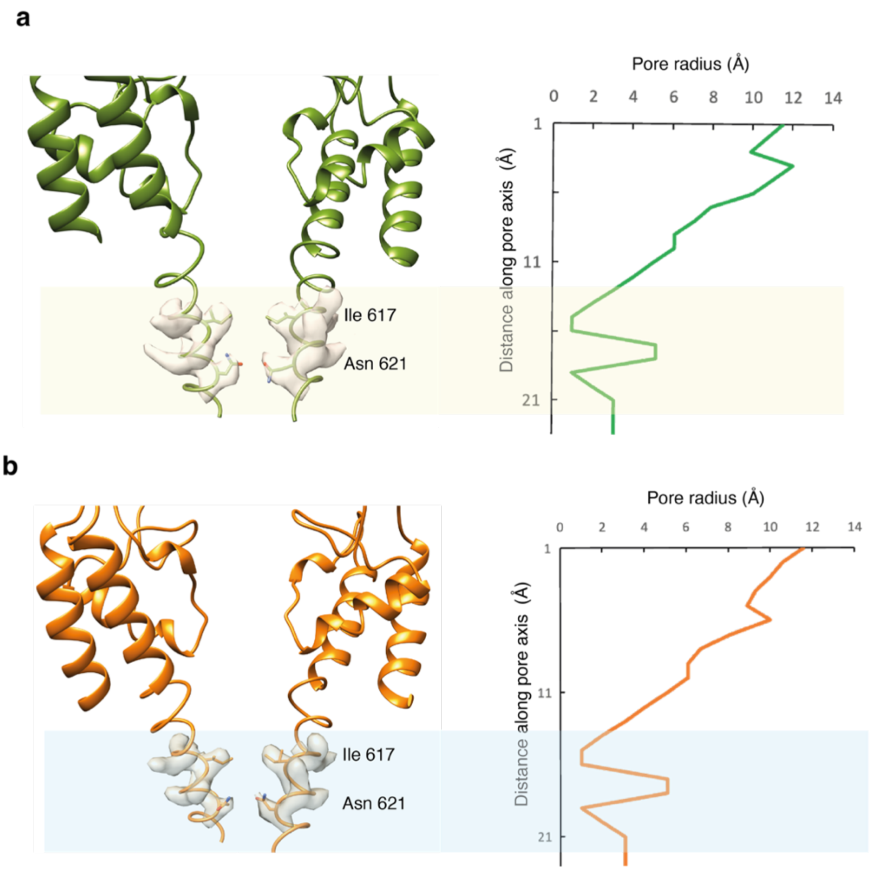

**Figure S7.** Pore analysis of TRPC4$_{DR}$ structures with bound *E/Z*-**AzPico**. Asymmetric pore radius calculation of the **(a)** TRPC4:(*E*)-**AzPico** and **(b)** TRPC4:(*Z*)-**AzPico** structures with PoreWalker[35] indicates the tight constriction at the lower gating residues Ile617 and Asn621 (transparent shaded region), indicating that both structures are in a closed state.

##### 1.3.3 Note on TRPC4 / TRPC5 Sequence Alignment

The key contact residues of interest in the ligand binding site are highly conserved between TRPC4 and TRPC5. The relevant sequences in the site are:

**TRPC4$_{DR}$, 501 - 529:**

```
HLGPLQISL  GRMLLDILKF  LFIYCLVLLA
```

**hTRPC5, residues 502 - 530:**

```
HLGPLQISL  GRMLLDILKF  LFIYCLVLLA
```

##### 1.3.4 Differences and selectivity between TRPC5 and TRPC4?

**AM237** activates TRPC5 but inhibits **EA** mediated activation of TRPC4; **Pico145** inhibits both TRPC4 and TRPC5. Thus, somewhat naively, it could be argued that it is the *para* substituent here which affects both the efficacy mode and the selectivity for TRPC4 and/or TRPC5.

Intriguingly, ***Z*-AzPico** activates TRPC4 and TRPC5 with low nanomolar potency whereas ***Z*-AzHC** only activates TRPC5 but not TRPC4. Therefore, the *meta* substituent is now responsible for the selectivity in this case. The cryo-EM structures of **Pico145** and **HC-070** in complex with TRPC5 indicated the same binding site but the southern phenyl ring bearing the $OCF_3$/Cl group is flipped and pointing in opposite directions. Our structural data with **AzHC**:TRPC5 and **AzPico**:TRPC4 does not reproduce the ring flip but shows the same orientation shift for the diazene motif for *E* and *Z* isomers, i.e. *Z*-buried vs. *E*-projected. Additionally, both ***E*-AzPico** and ***E*-AzHC** inhibit **EA** mediated activation on TRPC4 and TRPC5, hence both scaffolds can bind to either protein but the *efficacy switch* does not occur for **AzHC** on TRPC4.

It is not surprising that both ligands bind to either protein because there is a high sequence similarity among the TRPC family in the identified binding pocket for **Pico145** /**HC-070**, and in the binding region, human TRPC4/5 differ only by a single amino acid (TRPC5: V579, TRPC4: I579). We hypothesised that the selective activation of TRPC5 vs TRPC4 by **AzHC** (and **AM237**[8]) could be the result of differences in this residue. To test this hypothesis, we used $[Ca^{2+}]_i$ recordings to study the effect of **AzHC** and **AM237** on the variant TRPC5-V579I (in which V579, the only residue in the **AzHC** binding site that is different between TRPC4/5, is mutated to its TRPC4 counterpart), and the reverse variant TRPC4β-I575V (residues correspond exactly; the sequence numbering differs between the channels). These point mutations do not switch the effect of **AM237** and **AzHC** (**Figure S8**), suggesting that the molecular basis for **AzHC**'s TRPC5-selective activation is more complex than the immediate residues it contacts, and perhaps is regulated allosterically by other TRPC4/5 domains instead of direct binding within this pocket.

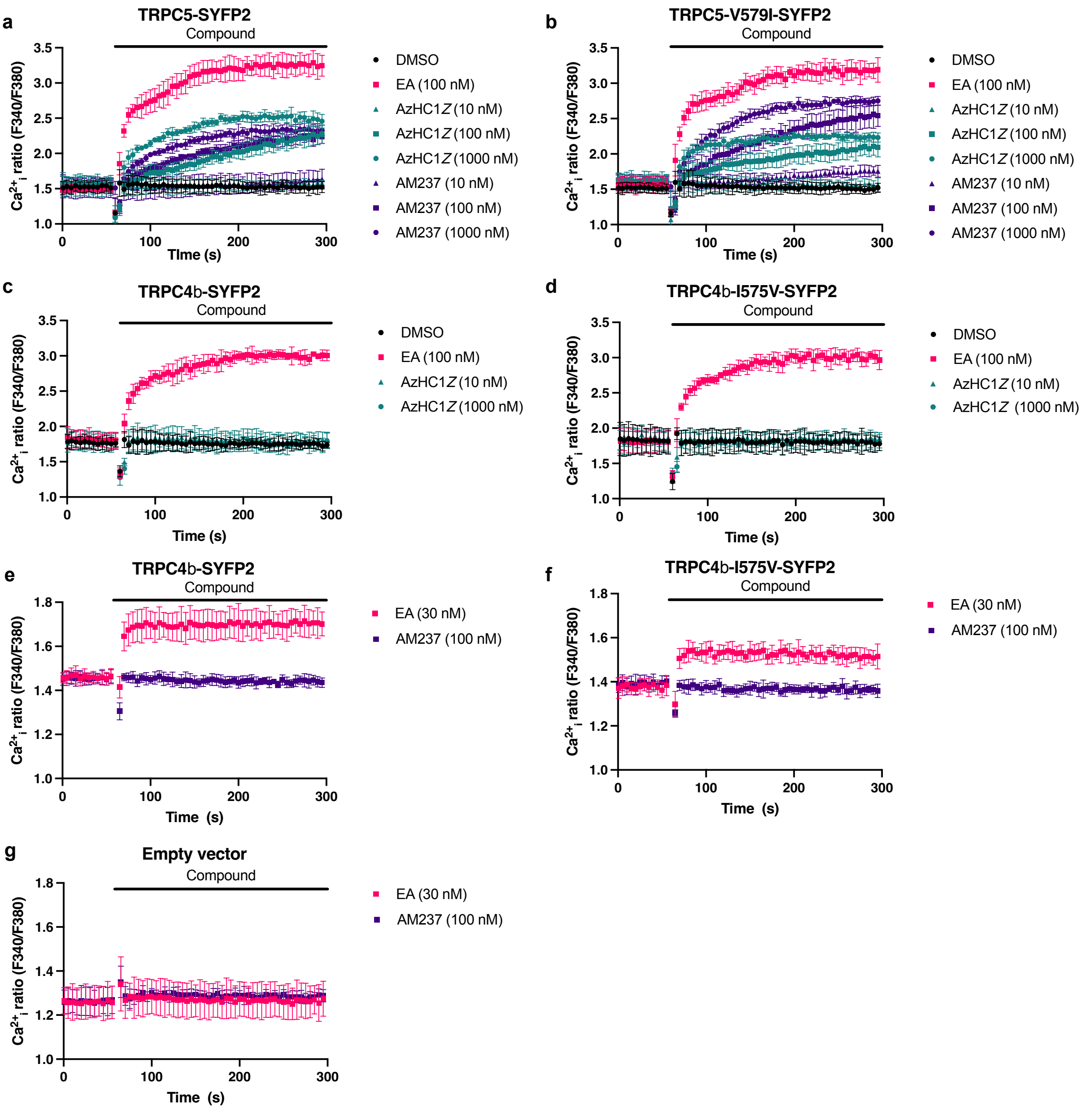

**Figure S8**. Selective TRPC5 activation by **AM237** and **AzHC** is not the result of differences in residues lining the AzHC binding site. $[Ca^{2+}]_i$ measurements were made from HEK 293 cells transiently expressing the indicated plasmid. Each panel displays data from a single 96-well plate (N = 6; mean ± SD over technical replicates) showing the baseline recording (0-60 s) followed by the effect of addition of the indicated compound (61-300 s). TRPC1/4/5 activator (-)-englerin A (**EA**) was used as a positive control to determine the maximum $[Ca^{2+}]_i$ response. **(a-b) AM237** and (*Z*)-**AzHC** are partial agonists of TRPC5 as well as $TRPC5_{V579I}$. **(c-f) AM237** and (*Z*)-**AzHC** do not activate TRPC4, nor $TRPC4_{I575V}$. **(g)** This control experiment shows that transfection with TRPC4/5 is necessary for detection of a $[Ca^{2+}]_i$ response to **EA** or **AM237**.

#### 1.4 Supporting Note 5: AzHC & TRPC5-dependent $Ca^{2+}$ in mouse hypothalamus

***Expanded from main text:***

TRPC5 is expressed in dopamine (i.e., tyrosine hydroxylase-positive, Th+) neurons of the dorsomedial aspect of the hypothalamic arcuate nucleus (ARC) (**Fig. 5a**), which has been confirmed by molecular biology and electrophysiological techniques.[36] The channel contributes to both spontaneous oscillatory activity and sustained activation following stimulation with the hormone prolactin, which is essential for normal prolactin homeostasis in the body.[36,37] Using brain slices through the ARC of mice expressing the $Ca^{2+}$ indicator GCaMP6f in Th+ neurons (Fig. 4), we investigated (1) whether $Ca^{2+}$ responses at endogenous TRPC5 expression levels can be photoswitched by UV illumination of **AzHC** or **AzPico**, (2) whether $Ca^{2+}$ responses depend on TRPC5 expression using TRPC5-deficient brain slices, (3) whether a dose dependence could be detected, and (4) whether there are properties in channel activity that can be distinguished between photoswitchable small TRPC modulators.

Th+ neurons are known for their spontaneous oscillatory burst-firing activity,[36] which is also evident in their spontaneous $Ca^{2+}$ responses (**Fig. 5b**). These spontaneous rhythmic $Ca^{2+}$ activities are characterized by periods of high and low $Ca^{2+}$ fluorescence.[37] Th+ neurons treated with 500 nM **AzHC** and stimulated with a 355 nm UV laser for ~60 ms inducing an E→Z isomerization (***Z*-AzHC**) induced a sustained high $Ca^{2+}$ fluorescence that lasted up to 3 min (14 of 33 cells, 42%). Genetic deletion of the TRPC5 channel in the Th-GCaMP6f-ΔTrpc5 mouse, prevented the increase in ***Z*-AzHC** induced $Ca^{2+}$ activity in Th+ neurons (**Fig. 5c**), indicating channel selectivity of **AzHC**. The area under the curve (AUC) of $Ca^{2+}$ signals quantifies that **AzHC** does not induce changes under 488-nm illumination alone, but requires both isomerization with 355-nm light ($p < 0.0001$) and TRPC5 expression to produce $Ca^{2+}$ increases ($p < 0.001$) (**Figure S21ab**).

To determine the sensitivity of ***Z*-AzHC** induced $Ca^{2+}$ responses in Th+ neurons, we performed concentration-response measurements (**Fig. 5d**). These results showed a dose dependence with a Hill coefficient of 1.95 and an EC50 of 0.13 ± 0.10 µM. At **AzHC** concentrations of 1 µM, no change in spontaneous $Ca^{2+}$ activity was observed in TRPC5-deficient Th+ neurons compared with before UV stimulation (**Fig. 5e**).

In wildtype slices, **AzPico** also induced long-lasting high $Ca^{2+}$ signals after 355 nm pulsing (13 out of 31 cells), with overall $Ca^{2+}$ signal properties that are not distinguishable from *Z*-**AzHC**. A comparison of both ***Z*-AzHC** and ***Z*-AzPico** illustrates that both photoswitchable modulators similarly prolonged the mean $Ca^{2+}$ burst duration and consequently lowered the mean $Ca^{2+}$ burst frequencies, as well as increased the mean amplitude of the $Ca^{2+}$ responses (**Fig. 5g,h; Figure S21d-l**).

The total duration of the $Ca^{2+}$ fluorescence increase after 355 nm UV illumination indicates that ***Z*-AzHC**, ***Z*-AzPico** and the photoswitchable agonist ***Z*-BTDAzo**[37] are very effective in elevating the $Ca^{2+}$ response in Th+ neurons ($p < 0.001$; **Figure S21g**). Comparing the properties of the TRPC5-dependent ***Z*-AzHC** and ***Z*-BTDAzo** the main difference is the higher frequency of the ***Z*-BTDAzo** $Ca^{2+}$ signals ($p < 0.0001$; **Fig. 5h**). Consequently, **BTDazo** had mainly shorter lasting $Ca^{2+}$ burst durations (29 out of 35 cells, 83 %) and only few mean $Ca^{2+}$ durations up to 3 min (6 out of 35 cells, 17 %). These results can suggest a different binding site or interaction sites in the same binding pocket for **AzHC** and **BTDAzo** in the TRPC5 channel.

### 2 Photocharacterization

**UV-VIS.** UV-Vis spectra were recorded on an Agilent Cary 60 UV-Vis spectrophotometer using 1 cm quartz or PMMA cuvettes. All photoisomerisations were performed at room temperature in non-degassed solvents unless stated differently. Samples were irradiated with either a CoolLED pE-4000 or Polycon V monochromator (TILL Photonics, Gräfelfing, Germany) by shining from the top of the cuvette, until the spectra did not change further (photoequilibrium). Unless stated differently, all measurements were performed at a default concentration of 20 µM. *Dark* state refers to stocks in DMSO kept at 60 °C for >14 h prior to measurements (all-*trans*). All other measurement methods have been detailed elsewhere (refs [37–39]). All-*E* and PSS spectra are shown in **Figure S9**.

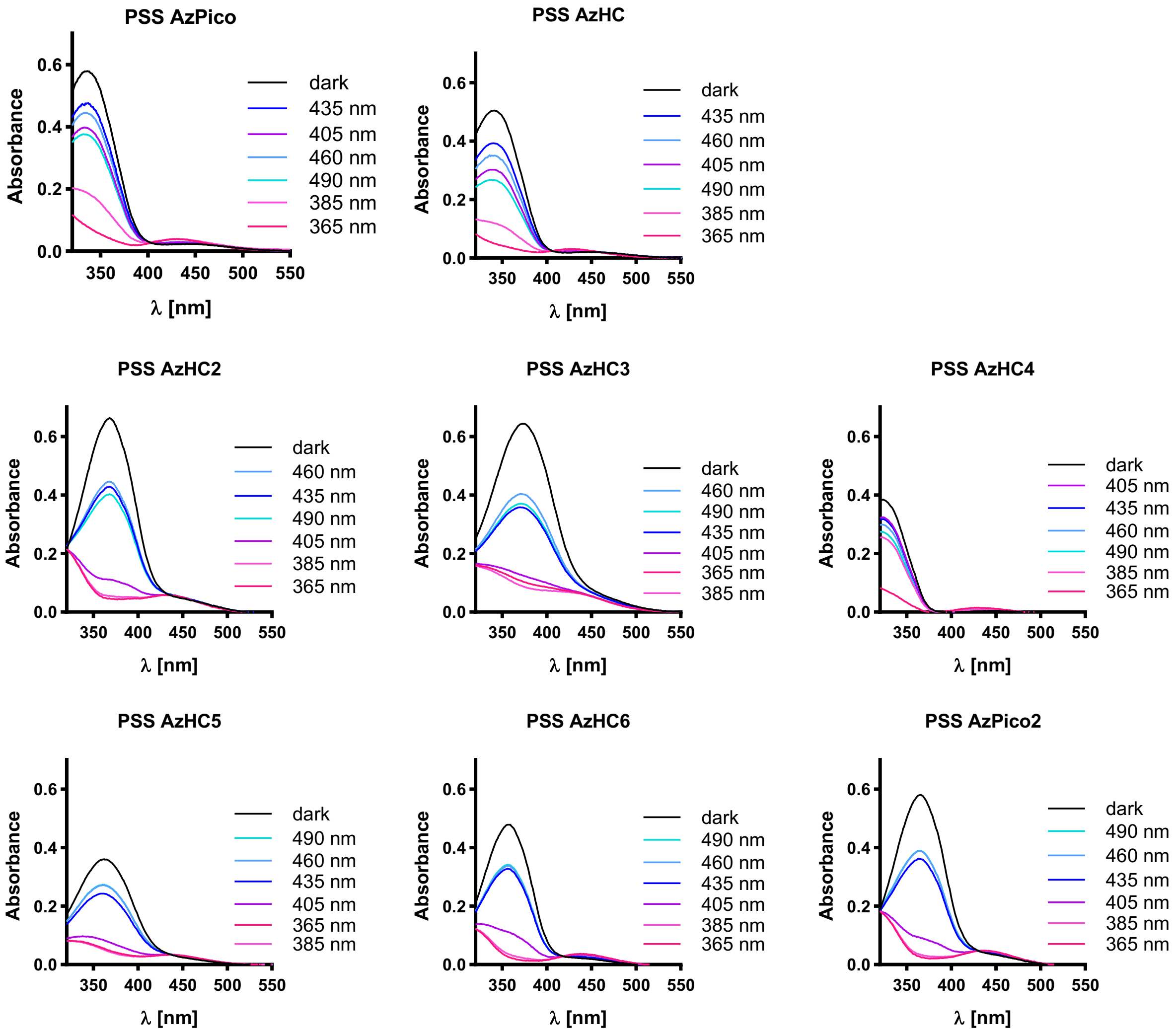

**Figure S9**: PSS spectra of **AzPico1-2** & **AzHC1-6** at different wavelengths. Measured at 20 µM in DMSO/$H_2O$ 9:1 (volumetric ratio). Illumination with CoolLED pE-4000. (**AzHC1** == **AzHC**, **AzPico1** == **AzPico**).

**HPLC for PSS *E/Z* ratios.** Analytical HPLC was performed on an Agilent 1200 SL with (a) a binary pump able to deliver $H_2O$:MeCN eluent mixtures containing 0.1% formic acid at a 1 mL/min flow rate [for PSS measurements of final compounds, only MeCN was used], (b) YMC Carotenoid 5µM, 4.6 x 1500 mm, maintained at 40 °C, (c) an Agilent 1200 series diode array detector. Samples (100 µM, MeCN) in HPLC vials were irradiated for at least 6 min from top with the monochromator light source (slit opening set to maximum of 15 nm) until PSS was reached, then injected (*E* and *Z* separate well); PSS ratio was then determined by integrating the signal at an isosbestic region found by UV/VIS in pure MeCN, and calculating the ratio from the integrals (**Table S3**).

**Table S3**. PSS %*E* & %*Z* for **AzPico** & **AzHC** at different wavelengths (monochromator, narrow-bandwidth), determined with HPLC integration (296 nm).

| | **AzPico** | | **AzHC** | |
|---|---|---|---|---|
| λ [nm] | % *E* | % *Z* | % *E* | % *Z* |
| 330 | 16 | 84 | 21 | 79 |
| 340 | 8 | 92 | 6 | 94 |
| 350 | 5 | 95 | 4 | 96 |
| 360 | 5 | 95 | 4 | 96 |
| 370 | 11 | 89 | 7 | 93 |
| 380 | 23 | 77 | 23 | 78 |
| 390 | 38 | 62 | 52 | 48 |
| 400 | 79 | 21 | 74 | 26 |
| 410 | 84 | 16 | 81 | 19 |
| 420 | 84 | 16 | 80 | 20 |
| 430 | 82 | 18 | 79 | 21 |
| 440 | 80 | 20 | 76 | 24 |
| 450 | 77 | 23 | 72 | 28 |
| 460 | 74 | 27 | 68 | 32 |
| 470 | 70 | 30 | 63 | 37 |
| 480 | 67 | 33 | 59 | 41 |
| 490 | 64 | 36 | 55 | 45 |
| 500 | 61 | 39 | 52 | 48 |

**Action Spectra, Cell-Free.** Samples were measured by UV-Vis with CooLED pE-4000 illumination (most LEDs are relatively narrow-band) during an initial 1 min dark (*E*) followed by alternating illumination cycles of 1 min at 365 nm (50 mW/mm$^2$) then 2 min of each indicated wavelength (10 mW/mm$^2$), to gain an impression of the relative rates per-photon of approaching PSS at the indicated wavelengths (**Figure S10**).

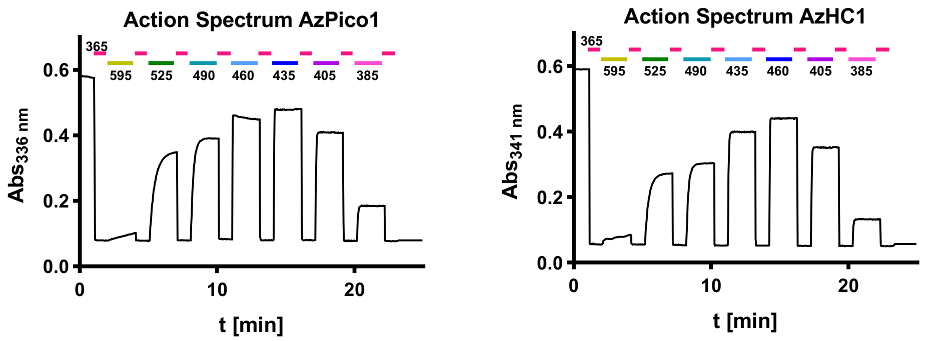

**Figure S10**. Action Spectra for **AzPico** & **AzHC** (LED illuminations, moderate-bandwidth).

**Spontaneous *Z*→*E* Relaxation, Cell-Free.** Samples were monitored by UV-Vis to confirm that spontaneous *Z*→*E* relaxation is orders of magnitude slower than assay timescales, i.e. active bidirectional photoswitching is needed to modulate the activity of **AzPico** and **AzHC** (**Figure S11**).

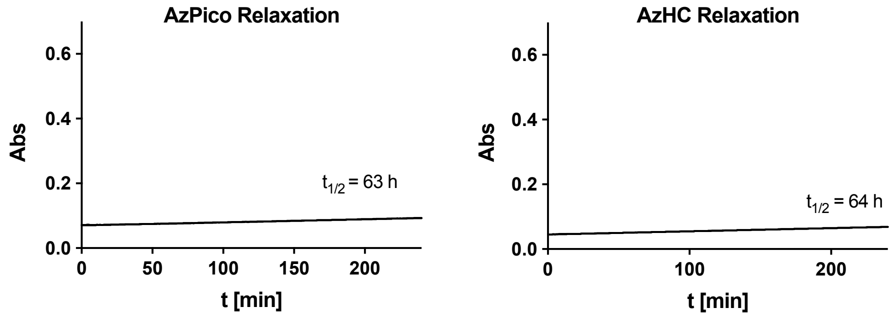

**Figure S11**. Very slow thermal *Z*→*E* relaxation of **AzPico** and **AzHC** at 37°C at 20 µM in DMSO:$H_2O$ 9:1. Data was force-fitted as exponential decay to obtain *lower limits* to the relaxation half times.

### 3 Cultured Cell Lines (primarily for Figure 1 and Figure 2)

#### 3.1 FLIPR $Ca^{2+}$ influx assay in cell suspensions.

We initially screened for photoswitchability of activity in cells, using a fluorometric imaging plate reader (**FLIPR**) calcium flux assay, with human embryonic kidney cell line HEK293 stably transfected to express mouse TRPC4β- or TRPC5-CFP fusion protein ($HEK_{mTRPC4\beta\text{-}CFP}$ / $HEK_{mTRPC5\text{-}CFP}$). The FLIPR setup provides high-content data, but it is limited to photocontrol LED wavelengths of 365 nm (best-*Z*) and 447 nm (suboptimal-*E*), while imaging at 470 nm excitation (Fluo-4 AM as $Ca^{2+}$ indicator; this imaging light counteracts both PSSs). Therefore, we expected the FLIPR results to under-estimate the true photocontrol power accessible to the reagents.

Cells were grown to 70%-90% confluency in a 75-cm$^2$ culture flask, trypsinised, washed and resuspended in BSA-containing (0.1%) HBS, and supplemented with 3 µM Fluo-4/AM. As described earlier, stably TRPC-transfected cells were induced with tetracycline 24 h prior to the experiments.[40,41] After incubation at 37°C for 30 min [Fluo-4 loading], cells were again washed, resuspended, and dispensed (40 µl/well) into black pigmented, clear-bottom 384-well plates (Greiner µ-clear). To image the Fluo-4 fluorescence during the application and photoswitching of test compounds, these plates were first mounted to a Tecan Fluent 480 (Tecan, Männedorf, Switzerland) liquid handling device, equipped with a 384-tip multichannel arm and a gripper arm. A custom-made plate-imaging device was used for photoswitching during fluorescence imaging, which features LED-based excitation modules (365 nm and 447 nm used here for photoswitching, and 470 nm used here for Fluo-4 readout) projected onto the bottom of the microwell plate, with a cooled scientific complementary metal oxide sensor (sCMOS) camera (Zyla 5.5, Andor, Belfast, UK) equipped with a Nokton 42.5 mm f/0.95 (Voigtlaender, Fürth, Germany) for detection. The LEDs were controlled with an Arduino device and constant current drivers (1 A for 365 and 470 nm LEDs; 3 A for 447 nm LED); detection was under the control of Micromanager 2.0 software. Compounds were serially diluted and applied to the Fluo-4-loaded cell suspensions inside this custom plate-imaging device, then fluorescence intensities (under 470 nm excitation) were continuously recorded during programmed illuminations e.g. with alternating 365 nm and 447 nm phases.

**AzPico**, **AzPico2**, **AzHC**, and **AzHC2** all showed photoswitchable activity on TRPC5 and were tested also on TRPC4 (**Figure S12**); the other tested compounds (**AzHC3-6**) did not show activity on TRPC5 and were not tested on TRPC4.

Concentration-response curves for the *Z*-isomers (channel activation) were then constructed from the maximal signals during 365 nm phases, after a suitable number of "priming" cycles to allow the system to reach stable performance (**Figure S13**).

All experiments were conducted as a minimum of 3 independent experiments with each experiment averaging data in technical duplicates.

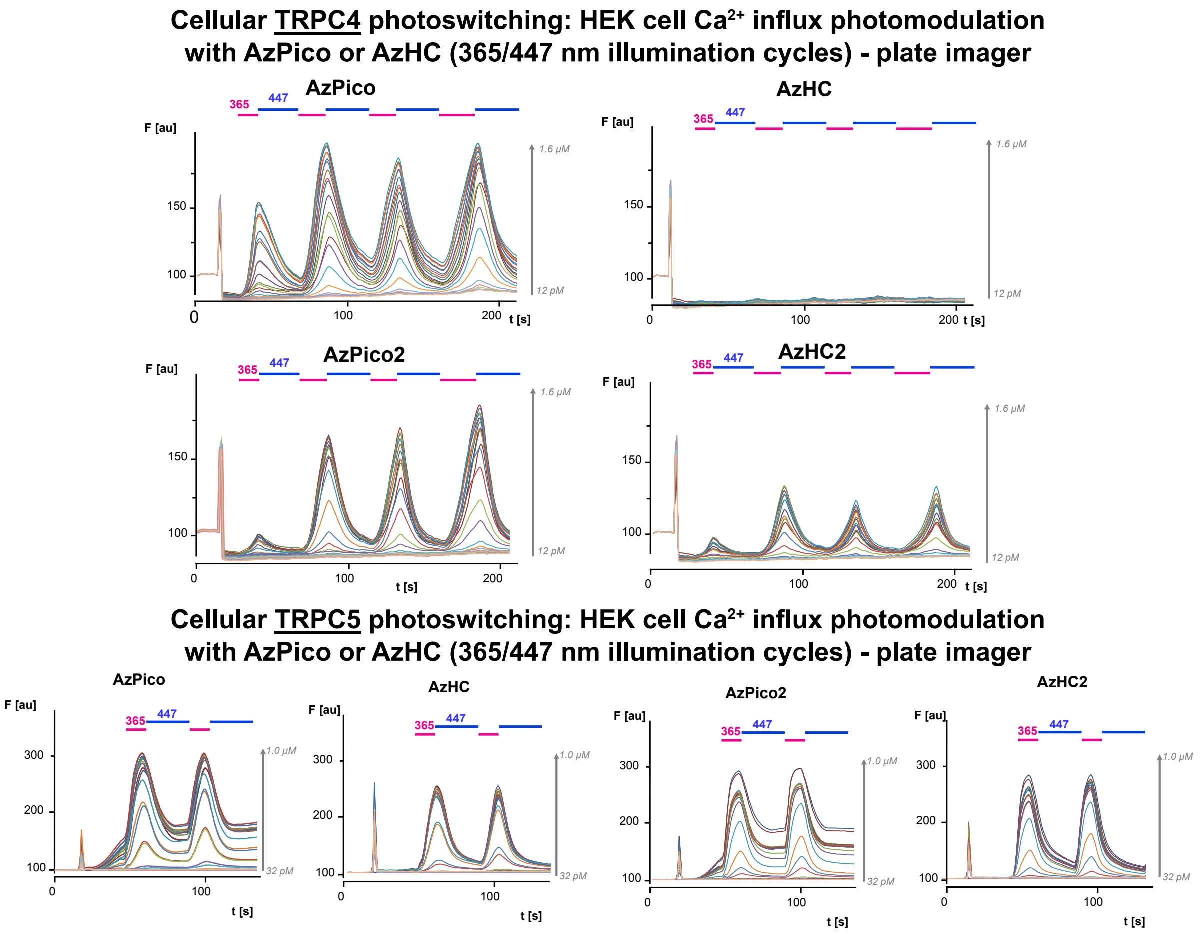

**Figure S12**. Activity of **AzPico**, **AzPico2**, **AzHC**, **AzHC2** on TRPC5 and TRPC4 (FLIPR assay).

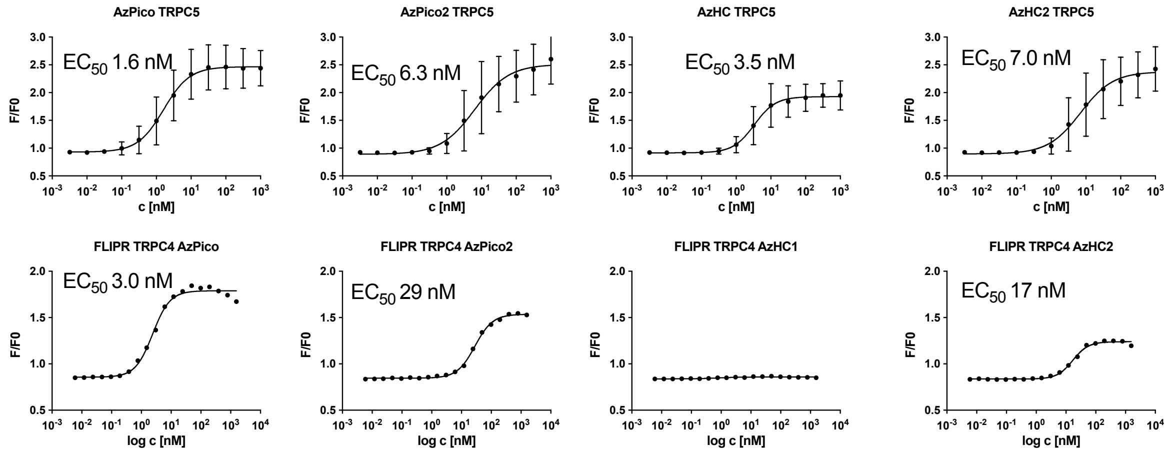

**Figure S13**. Dose response curves of *Z*-**AzPico**, *Z*-**AzPico2**, *Z*-**AzHC**, and *Z*-**AzHC2** on TRPC5 and TRPC4 (FLIPR assay).

#### 3.2 Ephys Characterisation of Photocontrolled Induced Ionic Currents. @MS

***Expanded from main text:*** We recorded full wavelength-dependent action spectra of **AzPico** and **AzHC** under fast photoswitching in ephys in TRPC4 and/or TRPC5-expressing cells (**Figure 2**, **Figure S14**). The advantage of the ephys setup is that freely chosen monochromated wavelengths can be used for photoswitching, while in parallel there is no optical imaging light to counteract these optimal wavelengths: and this allows for optimising the photoswitchability. They now performed to the full potential of ideal efficacy switches. For example, **AzPico** reversibly photomodulated TRPC4 currents over >60 consecutive cycles with a fully constant activation profile, without fatigue (**Figure 2a,b**). Current/voltage (I/V) curves show a strong activation of ion flux at best-*Z* (360 nm) PSS; yet, only at nonphysiological voltages (<-80 mV or >+60 mV) could any small differences between basal activity and good-*E* (440 nm) PSS be detected (**Figure 2c**). The repeatability of on/off photocycling allowed us to extract action spectra for both (1) photoactivation (**Figure 2d,e**) and (2) photodeactivation (**Figure 2f,g**) *in situ* in live cells. To obtain the activation action spectrum we used an irradiation protocol with varying wavelengths from 330-505 nm for 500 ms followed by 200 ms of 440 nm to set the channel activity back to a constant baseline value. For the deactivation action spectrum we first activated with 300 ms of 360 nm and then applied varying wavelengths from 330-505 nm for 200 ms followed by a baselining step with 360 nm and 440 nm. Optimal on-responses were elicited in the range of 350-365 nm, whereas a broader wavelength range of 400-480 nm can be applied for effective off-photoswitching.

A special feature of a high-potency efficacy photoswitch with similar-affinity isomers is that, if binding site saturation is ensured, the PSS dependence of activity should give insights into the stoichiometry of target activity modulation that would not be accessible or reliable when simply titrating always-active ligand/s. For example, at 400 nm the PSS *E*:*Z* ratio is ca. 3:1 (**Table S3**), and there is no channel activation detectable (**Figure 2e,g**): giving the expectation that occupying one of the four TRPC4/5 binding sites with the low nanomolar xanthine agonist *Z*-**AzPico** is still insufficient for channel activity, if the other three sites are occupied by the inverse agonist *E*-**AzPico**. As a corollary, the observed complete shutdown of channel currents at 25% *Z*-occupancy is a dramatic reflection of the power of efficacy switching: a potency switch with only 75%-complete *Z*→*E*-switching away from such a potent *Z* isomer could not deliver such a functional ON-OFF switch when its biological response, at the quasi-linear dose-response region of a sigmoid curve, is given by the logarithm of the *Z*-concentration.

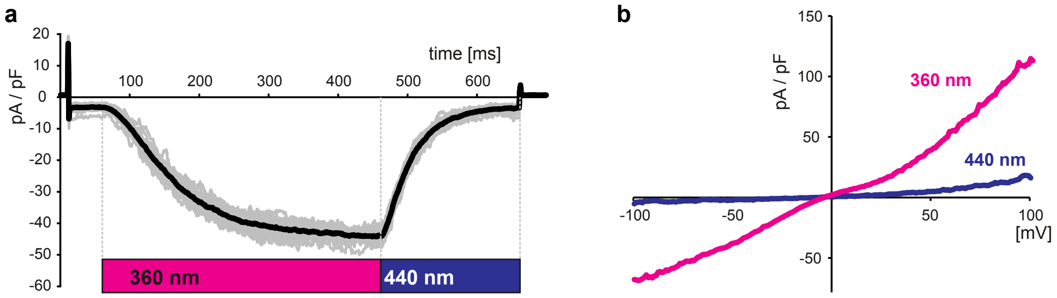

**Figure S14**. **(a)** Electrophysiological whole-cell recordings of TRPC5 currents in voltage clamp ($V_h$=-80 mV) mode. Overlay of ionic currents in a TRPC5-expressing HEK293 cell during >60 consecutive cycles of 360/440 nm illumination in the presence of 10 nM **AzPico**. **(b)** *I*/*V* curves of whole cell currents in voltage-clamped TRPC5-expressing cells exposed to 360 nm and 440 nm light with 10 nM **AzPico** in the bath solution.

#### 3.3 Pharmacology of TRPC4 and TRPC5 variants (Figure S8)

##### 3.3.1 Plasmids

Generation of TRPC5-SYFP2 and TRPC4β-SYFP2 has been reported previously.[8] TRPC5-V579I-SYFP2 and TRPC4β-I575V-SYFP2 were generated using the Q5® Site-Directed Mutagenesis Kit

(New England Biolabs) using back-to-back primers to insert the desired mutations. Primers were designed using NEBaseChanger® (https://nebasechanger.neb.com/) and purchased desalted from Integrated DNA Technologies.

**TRPC5-V579I-SYFP2** forward primer: 5' CTTCTGGTCTataTTTGGCCTTTTAAATC 3'; reverse primer 5' AGTGACTGAAGAGTCTCAAAG 3'

**TRPC4β-I575V-SYFP2** forward primer: 5' GTTTTGGTCAgtaTTTGGGCTCATC 3'; reverse primer: 5' AGGGACTGCAGTGTCTCA 3'.

All constructs were sequenced to verify identity (Azenta).

##### 3.3.2 Chemicals

**AM237** was prepared according to previously reported procedures.[8] (−)-englerin A (**EA**) was obtained from PhytoLab (Vestenbergsgreuth, Germany). **AM237**, **EA** and **AzHC** were made up as 10 mM stocks in 100% DMSO, aliquots of which were stored at −20 °C (**AM237** and **AzHC**) or −80 °C (EA). Further dilutions of compounds were made in DMSO and these were dissolved 1:1000 in a compound buffer (SBS + 0.01% pluronic acid) before being added to cells. Fura-2 AM (Invitrogen UK) was dissolved at 1 mM in DMSO and stored at –20 °C.

##### 3.3.3 Intracellular calcium recordings

Intracellular calcium ($[Ca^{2+}]_i$) was measured using the ratiometric $Ca^{2+}$ dye Fura-2. Experiments with TRPC5-SYFP2, TRPC5-V579I-SYFP2, TRPC4β-SYFP2, and TRPC4β-I575V-SYFP2 were carried out in HEK 293 cells (ATCC, Teddington, UK) transiently transfected with the relevant plasmid.

HEK293 cells were maintained in Dulbecco's Modified Eagle Medium, high glucose, GlutaMAX, + pyruvate (ThermoFisher Scientific), supplemented with foetal bovine serum (FBS (Merck); 10%), penicillin–streptomycin (ThermoFisher Scientific) (100 units per ml; 100 $\mu g ml^{-1}$). Cells were kept in a humidified incubator at 37 °C at 5% $CO_2$. Passage of the cells was carried out twice weekly.

Cells were transfected with the relevant plasmid using jetPRIME transfection reagent (VWR). Assays were performed 48 h after transfection. 24 h before experiments, transfected cells were plated onto black, clear bottom poly-D-lysine coated 96-well plates (ThermoFisher Scientific) at a density of 60,000 cells/well. To load cells with the Fura-2 dye, media was removed, and cells were incubated with standard bath solution (SBS; composition, in mM, NaCl 130, KCl 5, glucose 8, HEPES 10, $MgCl_2$ 1.2, and $CaCl_2$ 1.5) containing 2 µM Fura-2 acetoxymethyl ester (Fura-2 AM (Invitrogen UK)) and 0.01% pluronic acid (Merck) for 1 h at 37 °C. Dye-containing media was removed and the cells washed twice briefly with SBS. SBS was then changed to recording buffer (SBS with 0.1% BSA (Sigma Life Science) and 0.5% DMSO for **Figure S8ab**; SBS with 0.01% pluronic acid and 0.1% DMSO for **Figure S8c-g**) immediately prior to experimentation. Inclusion of BSA and a higher concentration of DMSO in **Figure S8ab** was to improve the solubility of **AzHC**. However, these additions did not affect the responses: recordings without BSA and with 0.1% DMSO gave near-identical data (not shown).

For **Figure S8a-d**, loaded compound buffer on a clear round-bottom 96-well plate (Sarstedt, Germany) was illuminated at 380 nm by the *DISCO* system[42] for a minimum of 20 min. Ambient light was excluded from this plate between DISCO illumination and $Ca^{2+}$ measurement. The level of activation seen to be produced by **EA** and **AM237** indicated these compounds were not degraded by illumination at 380 nm.

$[Ca^{2+}]_i$ was measured in the FlexStation3 (Molecular Devices, Wokingham, UK), by alternating excitation at 340 nm and 380 nm, with an emission at 510 nm. Measurements were performed at rt for 300 s at 5 s intervals. **EA**, **AzHC**, and **AM237** were dissolved at 2× final concentration in compound buffer (see above for composition) and added to cells after recording for 60 s.

### 4 Structural biology (Figure 3)

#### 4.1 Structural biology of TRPC5:AzHC

##### 4.1.1 Plasmid

Structural studies of TRPC5:AzHC were conducted using the previously described construct MBP-hTRPC5$_{\Delta766–975}$,[31] which contains human TRPC5 in C-terminally truncated form (Δ766–975) with an N-terminal maltose-binding protein tag followed by a PreScission protease cleavage site. Bacmids and baculoviruses were produced according to the Bac-to-Bac protocol (Invitrogen). The BacMam vector was a kind gift from Professor Eric Gouaux (Vollum Institute).[43]

##### 4.1.2 Protein expression and purification

P2 virus was added to 2.0 million per ml of Freestyle™ 293-F Cells (ThemoFisher Scientific) in Gibco FreeStyle 293 Expression Medium (Invitrogen) at a final volume of 7.5% at 37 °C and 5% $CO_2$. After 8-12 h, 5 mM sodium butyrate (Sigma Aldrich) was added, and the temperature was lowered to 30 °C. After a further 40 h, cells were harvested by centrifugation and then frozen. The protein purification protocol was adapted from Duan et al.[44] Unless stated otherwise, all detergents (and amphipol PMAL-C8) were supplied by Generon and Anatrace. For a typical purification, a 200 ml cell pellet was thawed and resuspended with 20 ml of 1% DDM, 0.1% CHS, 150 mM NaCl (Sigma Aldrich), 30 mM HEPES (Sigma Aldrich) pH 7.4, 1 mM DTT (Fisher Scientific Ltd) and protease inhibitor cocktail (Sigma Aldrich), and incubated by rotating at 4 °C for 1 h. The insoluble material was removed by centrifugation at 10,000×*g* for 1 h at 4 °C. The soluble fraction was incubated with 500 µl bed volume of pre-washed amylose resin (New England Biolabs) for 12-16 h, rotating at 4 °C. The resin was washed with 30 ml of 0.1% DDM and 0.01% CHS, 150 mM NaCl, 30 mM HEPES pH 7.4, 1 mM DTT. The resin was resuspended in 8 ml of 0.2% PMAL-C8, 150 mM NaCl, 30 mM HEPES pH 7.4, 1 mM DTT and rotated for 6 h at 4 °C. The detergent was removed by addition of Biobeads (Bio-rad) at a ratio of 10 mg/ml, followed by incubation for 16-20 h. The resin was washed with 20 ml of buffer without detergent (150 mM NaCl, 30 mM HEPES pH 7.4, 1 mM DTT). To avoid denaturation of **AzHC** by DTT, we further washed the resin with 4 ml of the same buffer *without* DTT before elution. The sample was eluted in 2 ml of the same buffer (without DTT) plus 50 mM maltose (Sigma Aldrich). The eluate was subjected to centrifugation at 20,000×*g* for 15 min at 4 °C to remove any precipitated material. The supernatant was concentrated step-wise to the required concentration with 100 kDa cut off Vivaspin 500 concentrators (Sigma Aldrich). In a typical purification protocol, ~200 µg of purified TRPC5 was produced from 200 ml of cell suspension.

##### 4.1.3 Sample preparation and data collection

To access the required isomer of **AzHC**, the compound samples were illuminated with blue light (440 nm, 3W LED) for **(*E*)-AzHC** or UV-A light (365 nm, 3W LED) for **(*Z*)-AzHC**. Following this illumination, to minimise the risk of photoswitching, the compounds were manipulated only under red light (650 nm LED) or under the light source used to access the required isomer. For cryoEM studies, 1 mg of purified TRPC5 in PMAL-C8 at 1.0 mg/ml was incubated with 100 µM of the relevant **AzHC** isomer (taken from a 10 mM DMSO stock, never exceeding 1% DMSO final concentration in protein samples). A 3.5 µl aliquot of the sample was applied to an UltrAuFoil Holey Au R1.2/1.3, 300 mesh grid (Quantifoil), which had been glow-discharged twice for 45 s using a Pelco easyGlow glow discharge unit. An FEI Vitribot, retrofitted with the light source used to access the relevant **AzHC** isomer, was used to blot the grids for 6 s (blot force 1) at 100% humidity and 4 °C before plunging into liquid ethane. The grids were loaded into an FEI Titan Krios transmission electron microscope (Astbury Biostructure Laboratory, University of Leeds) operating at 300 kV, fitted with a Falcon 4i direct electron detector. Automated data collection was carried out using EPU software, with fringe-free imaging in counting mode, using a defocus range between −0.7 to −3 µm in 0.3 µm increments. We collected a total of 3,000 EER movies for the **(*E*)-AzHC** sample and

3,504 EER movies for the **(*Z*)-AzHC** sample. In both cases, the movies were collected with a pixel size of 0.74 Å and a total dose of 35.72 e−/Å$^2$. The fractions were combined into 27 frames resulting an exposure dose of 1 e−/Å$^2$ per frame.

###### 4.1.4 Image processing

An overview of the image processing protocol is shown in **Figure S11** and **Figure S12**. All processing was completed in Cryosparc-v4.1 and v4.4[45] unless stated otherwise. The initial drift and beam-induced motions were corrected for using *Patch Motion Correction* while CTF estimation was performed using *Patch CTF Estimation*, both with default settings. For particle picking we used a TRPC5 map previously obtained in-house, down-filtered to 20 Å to generate 50 templates using *Create templates* tool in Cryosparc. The obtained templates were used for template picking, resulting - after filtering the picks based on the NCC score (>0.26) - in 565k particles for the **(*E*)-AzHC** sample, and 702k particles for the **(*Z*)-AzHC** sample. After several rounds of 2D classification, particle stacks of 134k and 105k for **(*E*)-AzHC** and **(*Z*)-AzHC**, respectively, were produced. Further, *Ab-initio* was used to generate five initial models, from which at least one had the expected shape. The five models were used to sort particles further using *Heterogeneous refinement*, resulting in particle stacks of 87k for **(*E*)-AzHC** and 60k **(*Z*)-AzHC**. A further round of heterogeneous refinement with the fully extracted particle set, using as input the best model and three bad models to pull out bad particles was then performed. Using this approach, we observed that our final stack of good particles increased by roughly 20% while also improving particle orientation by either increasing the number of exotic orientation or by increasing the particle number in low-populated 2D classes. After the last heterogeneous refinement, we ended up with 161k particles for **(*E*)-AzHC** and 125k particles for **(*Z*)-AzHC**. A final round of 2D classification was used to remove bad particles or classes that did not show high resolution features. The final particle stack comprised 89k and 69k particles for **(*E*)-AzHC** and **(*Z*)-AzHC**, respectively. These particle stacks were taken forward and refined using non-uniform refinement with 5 extra passes, and having defocus refinement, CTF refinement (all option true) and EWS correction active. In both cases the structures were refined without imposing symmetry to a global resolution of 2.60 Å for **TRPC5:(*E*)-AzHC** and 2.90 Å for **TRPC5:(*Z*)-AzHC**, as estimated based on the gold standard FSC = 0.143 criterion. We used ResolveCryoEM[46] features of Phenix[47] to improve the interpretability of the map.

###### 4.1.5 Model building

The models of **TRPC5:(*E*)-AzHC** and **TRPC5:(*Z*)-AzHC** were built using ModelAngelo[48]. The models obtained were inspected and manually completed in Coot[49]. Several rounds of real-space refinement were performed in Phenix before fitting the corresponding ligand. For ligand fitting, we used LigandFit[50] in Phenix with small molecule constraints generated by eLBOW[51] function within Phenix, using AM1 geometry optimisation starting from a SMILE string. After automated fitting of the ligand, we manually checked the structures in Coot and performed addition real-space refinement in Phenix. Protein-ligand interactions were visualised with PoseEdit[52] (https://proteins.plus) and ChimeraX[53,54]. All structural images were produced using ChimeraX, Coot or one of the softwares used for protein-ligand visualisation.

###### 4.1.6 Additional TRPC5:AzHC Data and Figures

**Supporting Note 4** contains the figures and discussion that are key to the paper; this section contains figures and data on workflows (**Figures S15-S16**), statistics (**Table S4**), and data quality (**Figures S17-18**).

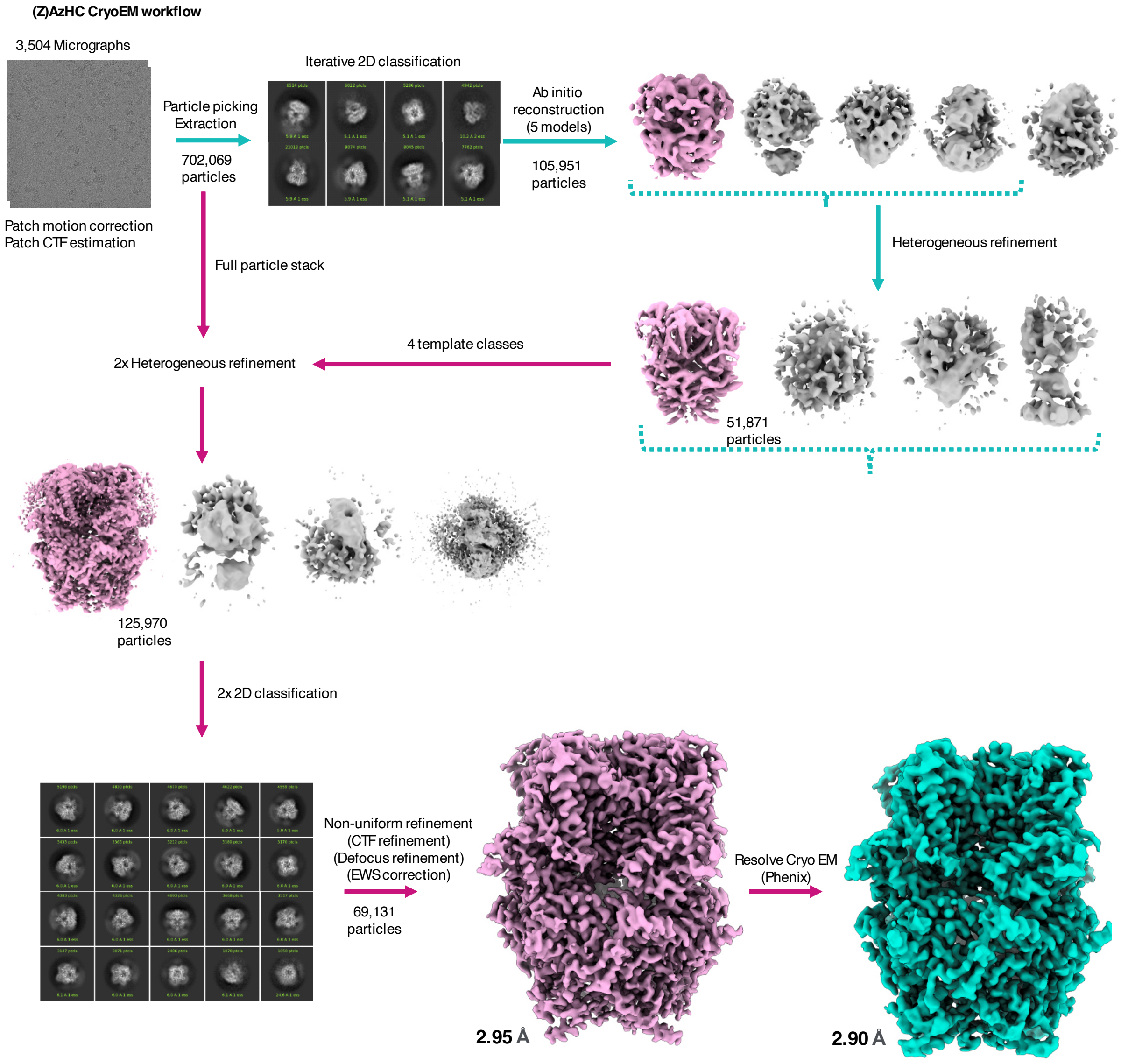

**Figure S15**. CryoEM workflow for TRPC5 in complex with (*Z*)-**AzHC**.

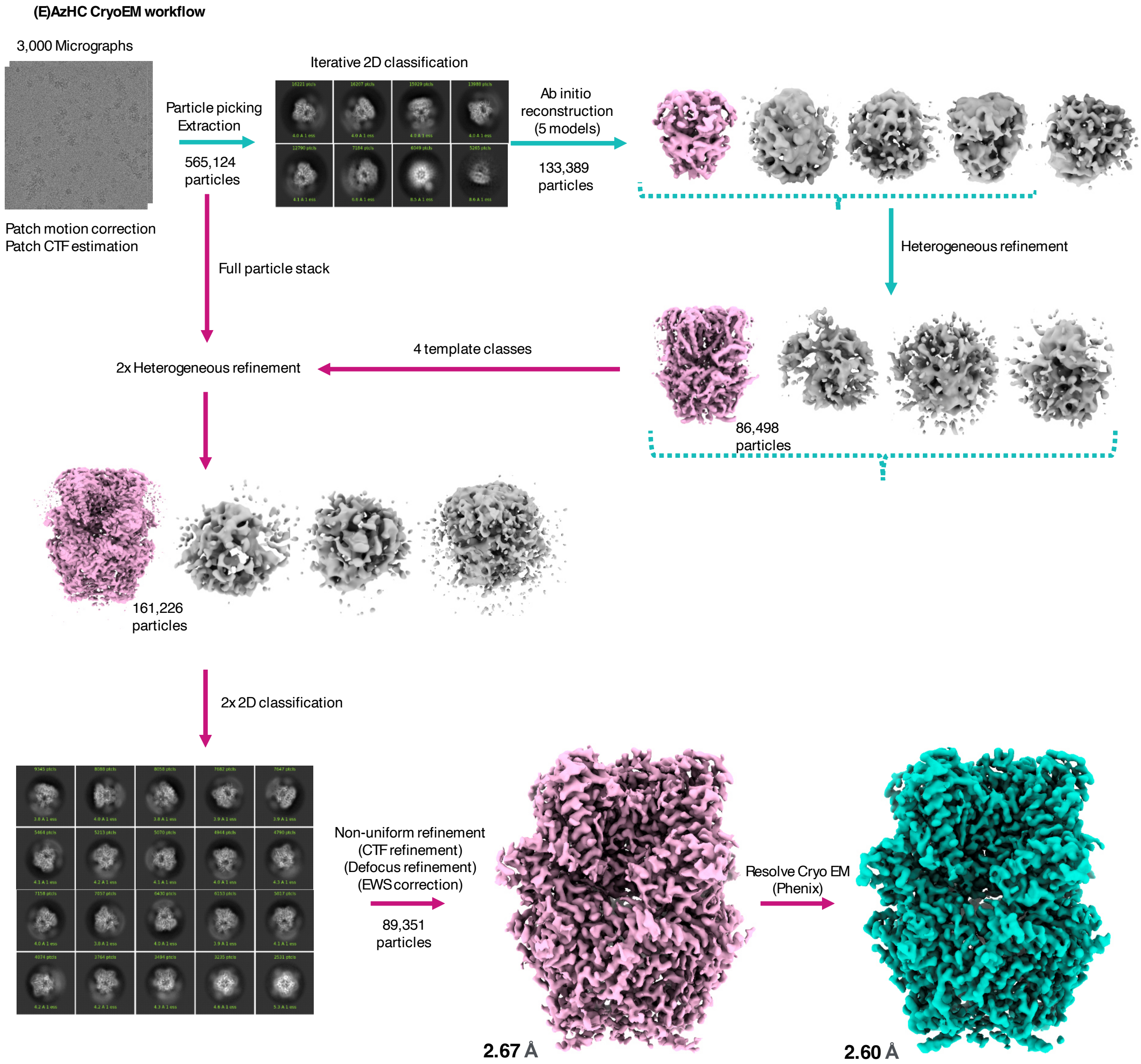

**Figure S16**. CryoEM workflow for TRPC5 in complex with (*E*)-**AzHC.**

**Table S4.** Cryo-EM data collection, refinement, and validation statistics.

| | **(*E*)-AzHC** | **(*Z*)-AzHC** |
|---|---|---|
| **Data collection and processing** | | |
| Magnification | 165k | 165k |
| Voltage (kV) | 300 | 300 |
| Electron exposure ($e^{-}/Å^{2}$) | 35.72 | 35.72 |
| Defocus range (µm) | -0.7 to -3.00 | -0.7 to -3.00 |
| Pixel size (Å) | 0.74 | |
| Symmetry imposed | C1 | |
| Final particle images (no.) | 89,351 | 69,131 |
| Map resolution range (Å) | 2.6 | 2.9 |
| FSC threshold | 0.143 | 0.143 |
| **Refinement** | | |
| Initial model | ModelAngelo | ModelAngelo |
| Map sharpening *B* factor ($Å^{2}$) | 66.4 | 72.1 |
| *Model composition* | | |
| Non-hydrogen atoms | 22307 | 22211 |
| Protein residues | 2705 | 2705 |
| Ligands | 4 | 4 |

| | | |
|---|---|---|
| *Bonds (RMSD)* | | |
| Length (Å) (# > 4σ) | 0.003 (0) | 0.003 (0) |
| Angles (°) (# > 4σ) | 0.368 (0) | 0.595 (8) |
| **Validation** | | |
| **MolProbity score** | **2.35** | **1.33** |
| EMRinger score* | 3.25 | 2.66 |
| Clash score | 18.52 | 6.10 |
| *Ramachandran plot (%)* | | |
| Outliers | 0.00 | 0.00 |
| Allowed | 2.14 | 1.54 |
| Favored | 97.86 | 98.46 |
| *Ramachandran Z-score* | | |
| whole | 1.60 (0.16) | 1.06 (0.16) |
| helix | 1.58 (0.12) | 1.04 (0.12) |
| sheet | --- (---) | --- (---) |
| loop | -0.06 (0.21) | 0.17 (0.22) |
| Rotamer outliers (%) | 5.33 | 0.62 |
| Cβ outliers (%) | NA | NA |
| *Peptide plane (%)* | | |
| Cis proline/general | 0.0/0.0 | 0.0/0.0 |
| Twisted proline/general | 0.0/0.0 | 0.0/0.0 |
| CaBLAM outliers (%) | 0.11 | 0.19 |
| *ADP (B-factors)* | | |
| Iso/Aniso (#) | 22211/0 | 22211/0 |
| *B* factors (Å$^2$) | (min/max/mean) | (min/max/mean) |
| Protein | 6.68/176.88/76.94 | 34.19/220.63/112.90 |
| Ligand | 13.55/105.99/46.90 | 44.59/149.06/86.47 |
| *Occupancy* | | |
| Mean | 1.00 | 1.00 |
| occ = 1 (%) | 100.00 | 100.00 |
| 0 < occ < 1 (%) | 0.00 | 0.00 |
| occ > 1 (%) | 0.00 | 0.00 |
| **Resolve CryoEM** | | |
| *Box* | | |
| Lengths (Å) | 119.88, 120.62, 141.34 | 111.74, 111.00, 137.64 |
| Angles (°) | 90.00, 90.00, 90.00 | 90.00, 90.00, 90.00 |
| Supplied Resolution (Å) | 2.6 | 2.9 |
| Resolution Estimates (Å) | Masked/*Unmasked* | Masked/*Unmasked* |
| d FSC (half maps; 0.143) | ---/--- | ---/--- |
| d 99 (full/half1/half2) | (2.8/---/---) / *(2.8/---/---)* | (3.3/---/---) / (3.3/---/---) |
| d model | 2.8/2.8 | 3.2/ 3.2 |
| d FSC model (0/0.143/0.5) | (2.3/2.4/2.6) / *(2.4/2.4/2.6)* | (2.3/2.3/2.9) / (2.3/2.3/2.9) |
| Map min/max/mean | -8.10/12.26/0.00 | -5.71/10.04/0.00 |
| **Model vs. Data** | | |
| CC (mask) | 0.89 | 0.88 |
| CC (box) | 0.64 | 0.64 |
| CC (peaks) | 0.68 | 0.65 |
| CC (volume) | 0.87 | 0.87 |
| Mean CC for ligands | 0.88 | 0.83 |

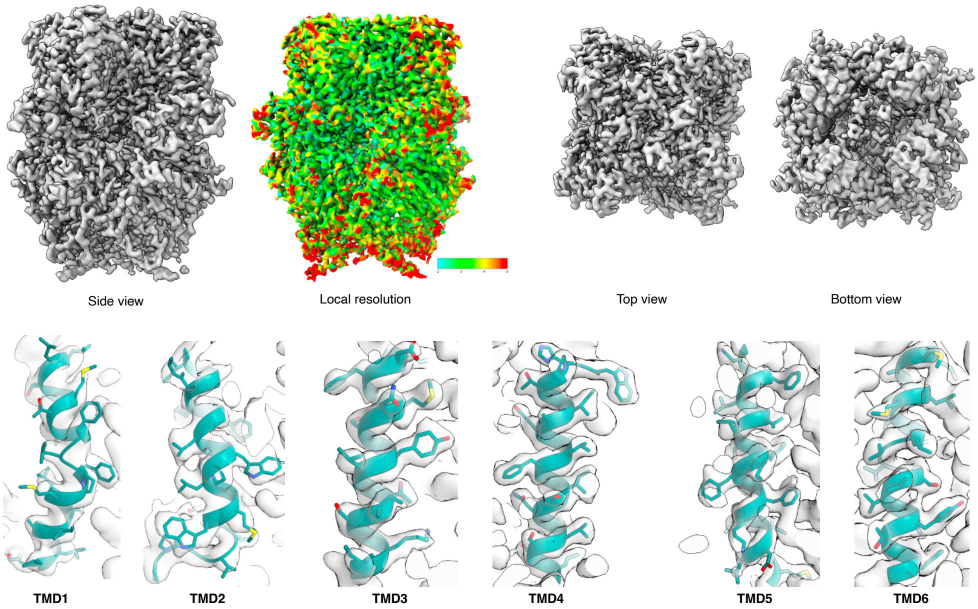

**Figure S17**. **CryoEM data quality 1.** Top: CryoEM map for TRPC5 in complex with **(*E*)-AzHC** (different views and local resolution shown). Bottom: Fit of the 6 trans-membrane domains of TRPC5 (teal) in the cryoEM map (grey mesh).

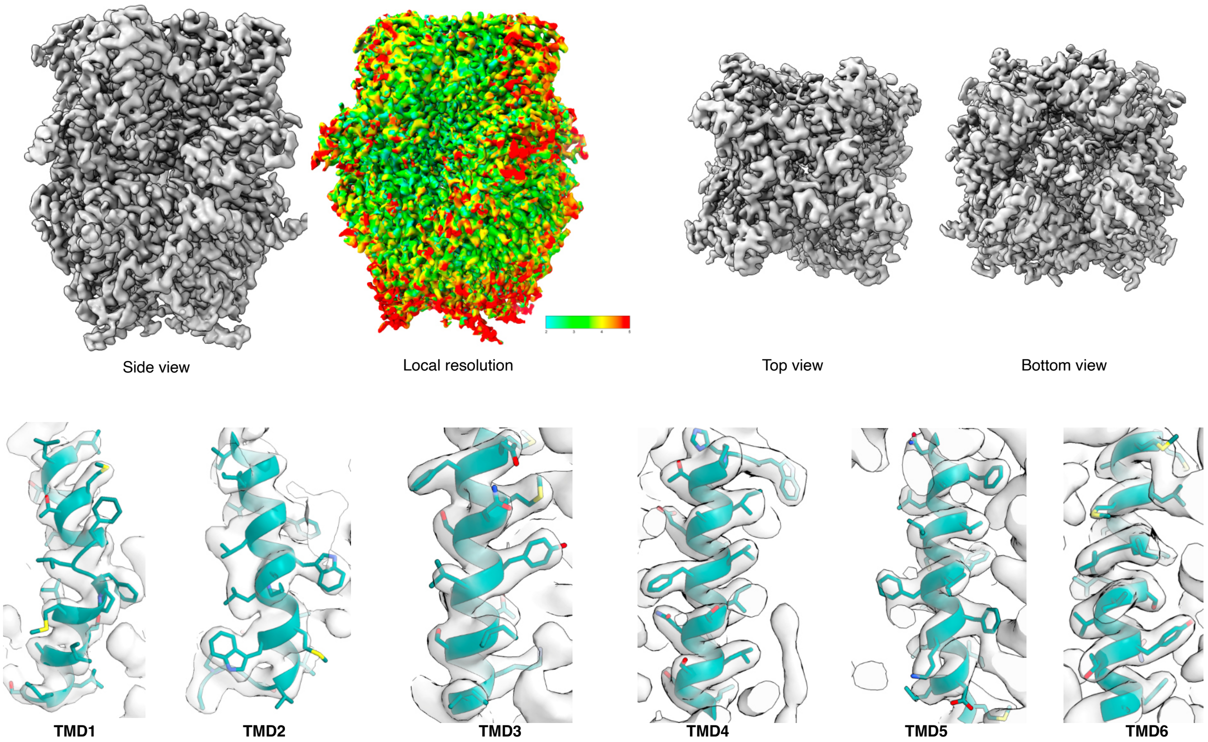

**Figure S18**. **CryoEM data quality 2.** Top: CryoEM map for TRPC5 in complex with **(*Z*)-AzHC** (different views and local resolution shown). Bottom: Fit of the 6 trans-membrane domains of TRPC5 (teal) in the cryoEM map (grey mesh).

#### 4.2 Structural biology of TRPC4:AzPico

##### 4.2.1 Cryo-EM grid preparation and screening

TRPC4$_{DR}$ from zebrafish was expressed and purified for cryo-EM as described previously.[33] For cryo-EM **AzPico** dissolved in DMSO was added to TRPC4 (final concentration of 100 µM, and 1% DMSO). The detergent was then exchanged with amphipols and incubated overnight before plunging using a Vitrobot cryo-plunger (FEI Thermo Fisher).

For the dark state (***E*-AzPico**), sample preparation was performed under red light illumination and residual light was minimized to prevent isomerization. In case of the bright state (***Z*-AzPico**), the molecule was activated for ~ 1 min using LEDs with a wavelength of 365 nm.

2.5 µl of TRPC4$_{DR}$ at a concentration of 0.45 mg ml−1 were applied onto freshly glow-discharged holey carbon grids (C-Flat (1.2/1.3) 400 mesh) blotted using 3.0 s blotting time, 0 blotting force with 100% humidity at 4°C and vitrified in liquid ethane cooled by liquid nitrogen.

##### 4.2.2 Cryo-EM data acquisition and image processing

Data sets were collected using EPU software on Titan Krios microscopes (FEI Thermo Fisher) operated at 300 kV and equipped with an X-FEG. All the datasets were collected using the aberration-free image shift (AFIS) feature of EPU to speed up the data-collection process. Equally dosed frames were collected on K3 (Gatan) direct electron detectors in super-resolution mode in combination with a GIF quantum-energy filter set to a filter width of 20 eV. The dataset was collected with a pixel size of 0.455 in super resolution mode. Typically, 60 frames were collected with a total dose of ~60 $e^{-}Å^{-2}$. Data collection was monitored live using TranSPHIRE,[55] allowing for direct adjustments of data acquisition settings when necessary, i.e. defocus range or astigmatism. Preprocessing included drift correction with MotionCor2,[56] creating aligned full-dose and dose-weighted micrographs. The super-resolution images were binned twice after motion correction. CTF estimation was also performed within TranSPHIRE using CTFFIND 4.1.13[57] on non-dose-weighted aligned micrographs. Unaligned frame averages were manually inspected and removed based on ice and image quality, resulting in a removal of 5–20% of the data sets. Following processing steps were performed using motion-corrected dose-weighted sums in the SPHIRE software package unless otherwise indicated.[58]

Single particles were picked automatically with crYOLO using the general model.[59] The particles were then windowed to a final box size of 300 × 300 pixels. Reference-free 2-D classification and cleaning of the data set was performed with the iterative stable alignment and clustering approach ISAC[60] in SPHIRE. A subset of particles producing 2-D class averages and reconstructions with high-resolution features were then selected for further structure refinement in Relion 3.0.[61] 3D classification was performed with C1 symmetry to classify the subpopulation. The classes having high-resolution features bound with ligands were selected and further polished and CTF-refined. Finally, the polished particles were exported to CryoSPARC[45] to improve the resolution with non-uniform refinement.

The previously reported model of TRPC4[33] was initially docked into the density and fitted into the map as rigid body using UCSF Chimera.[53] The model was further adjusted to fit in the density using Coot[62] with an iterative process of real space refinement in Phenix[63] and model adjustment in Coot until convergence as evaluated by model-to-map fit with valid geometrical parameters. For the ligand molecules, cif files were generated using eLBOW tool in Phenix and used as geometrical restraints in Coot and Phenix during modelling and refinement respectively.

##### 4.2.3 Additional TRPC4:AzPico Data and Figures

**Supporting Note 4** contains the figures and discussion that are key to the paper; this section contains figures and data on workflows (**Figure S19**), refinement and statistics (**Tables S5-S7**).

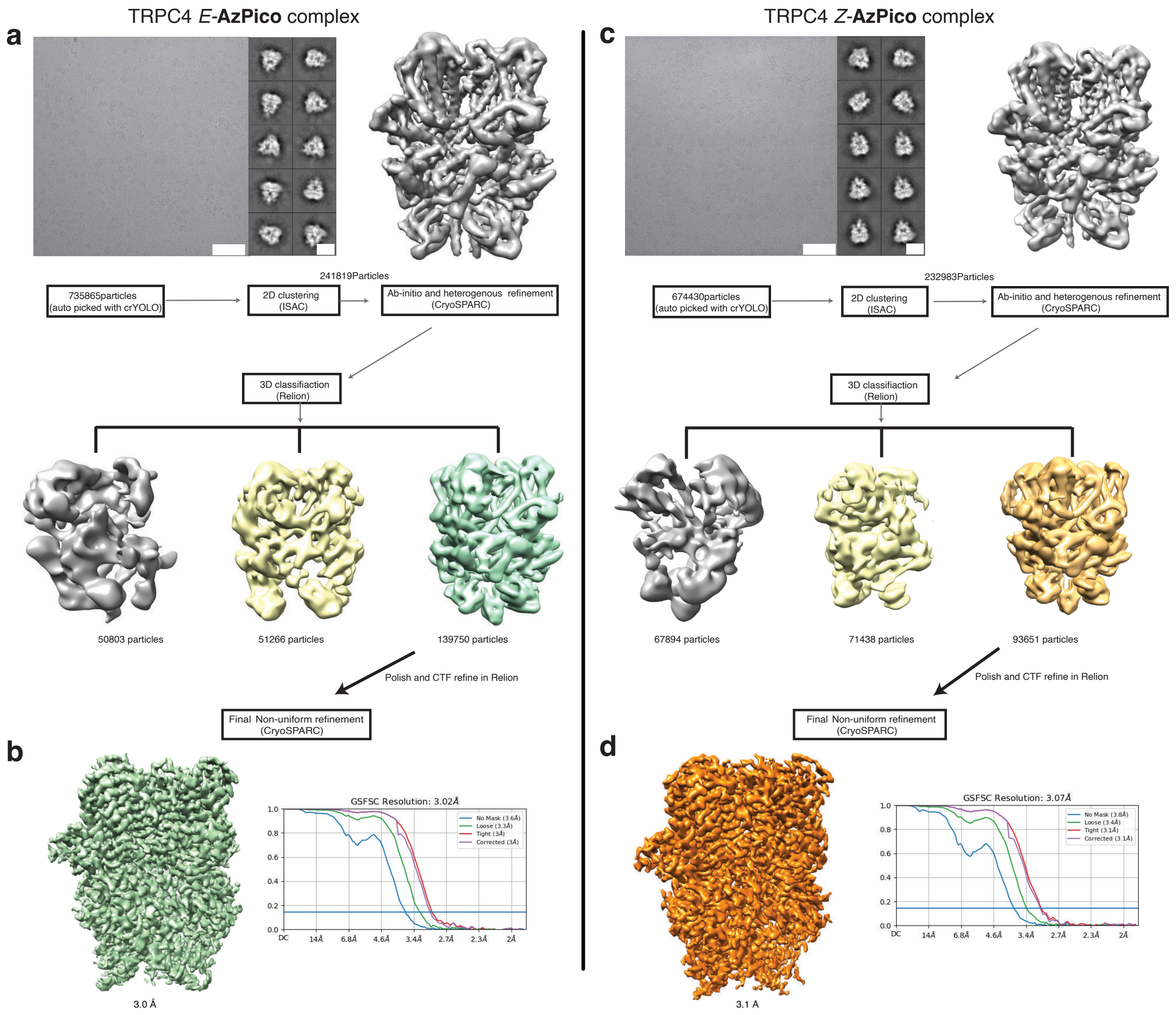

**Figure S19. Cryo-EM data processing of TRPC4 in complex with AzPico (*E* and *Z* isomers). (a)** Representative digital micrograph (scale bar, 50 nm) and selected 2D class averages (scale bar, 10 nm) of TRPC4 in complex with *E*-**AzPico**. The initial density obtained after *ab initio* and heterogeneous refinement is shown next to the 2D class averages. **(b)** Density obtained after 3D classification and a final round of non-uniform refinement. The Fourier shell correlation curves (FSC) corresponding to two independently refined particle subsets are shown. The horizontal light blue line indicates the FSC=0.143 criterion used for resolution estimation. **(c,d)** Same as in **a**,**b**, respectively, for the TRPC4 in complex with *Z*-**AzPico**. Scale bars: 50 nm and 10 nm, respectively.

**Table S5**. Data Collection details

| Sample | TRPC4-*E*-**AzPico** | TRPC4-*Z*-**AzPico** |
|---|---|---|
| Voltage [kV] | 300 | 300 |
| Defocus range [μM] | 0.48- 4.04 | 0.49- 3.07 |
| Camera | K3 Super resolution | K3 Super resolution |
| Pixel size [Å] | 0.455 /0.91[a] | 0.455 /0.91[a] |
| Exposure time [s] | 3.0 | 3.5 |
| Total electron dose [$e^{-}/Å^2$] | 52.47 | 56.34 |
| Frames per movie | 60 | 60 |
| Number of images | 2,598 | 3,311 |

**Table S6**. Refinement and model validation statistics

| Sample | TRPC4- *E*-**AzPico** | TRPC4- *Z*-**AzPico** |
|---|---|---|
| Number of particles used in refinement | 139750 | 93651 |
| Final resolution [Å] | 3.0 | 3.1 |
| Map sharpening factor [$Å^2$] | 111.7 | 101.9 |
| Electron dose particles final refinement [$e^-/Å^2$] | Polished particles | Polished particles |

**Table S7**. Model geometry and Refinement statistics

| | | |
|---|---|---|
| | Atomic model composition | |
| Non-hydrogen atoms | | |
| | Refinement (Phenix) | |
| RMSD bond | 0.002 | 0.002 |
| RMSD angle | 0.506 | 0.589 |
| Model to map fit, CC mask | 0.79 | 0.76 |
| Ramachandran plot (%) | Validation | |
| outliers | 0.12 | 0.12 |
| allowed | 4.14 | 4.62 |
| favoured | 95.74 | 95.26 |
| Rotamer outliers (%) | 2.95 | 4.25 |
| Molprobity score | 1.93 | 1.99 |
| EMRinger score | 1.86 | 2.46 |

### 5 Cultured neurons and chromaffin cells (Figure 4)

##### 5.1.1 Animals

All mice were kept according European Animal Welfare regulations and ethical guidelines from the local governing body (approval number of the Institutional Animal Care and Use Committee: **Az. 2.4.1.3/Bruns**). TRPC1/C4/C5 triple knockout (**145tko**), TRPC5 single knockout (**5ko**), and TRPC5 IC eR26 τGFP mice (i.e. TRPC5 knock-in, **5ki**) were generated as described previously;[64–66] C57BL/6N mice from Charles River, housed under the same conditions, were used as the wildtype (**wt**) controls.

##### 5.1.2 Primary Cell culture

<u>Primary hippocampal autaptic neurons</u> (i.e. neurons cultured under conditions where single neurons grow in isolation and make synapses only back onto themselves) were prepared as described[65]. Briefly, neurons were prepared from age matched **145tko**,[64] **5ko**, **wt** and **5ki**[65] P0-P1 mice (all C57Bl/6N strain) and cultured on pre-seeded astrocytic microislands. Neurons were grown for 10-17 days in NBA medium containing 2% B-27, 1% Glutamax and 1% penicillin/streptomycin at 5% $CO_2$ and 37°C. Only max. ca. 50% of **wt** cells express Trpc5;[65] while their results set expectations for the ability of **AzPico** to *directly* photostimulate wt hippocampal neurons, the **5ki** cells can be considered an appropriate control for the **5ko** and **145tko** cells in terms of determining the relevance of Trpc5 to the direct photocontrol of endogenous neuronal activity. Fluorescence imaging was used to confirm the knock-in efficiency in **5ki** cells before recording (**Figure S20**).

<u>Adrenal chromaffin cells</u> were prepared as described previously.[67] Briefly, chromaffin glands from adult wt and TRPC tko mice (10-12 weeks old) were digested in an enzyme solution containing 20-25 U/ml papain (Worthington). After inactivation, cells were washed with pre-warmed DMEM growth medium, supplemented with 0.4% PenStrep, and 1% ITSX (Thermo Fisher Scientific). After trituration, cells were plated onto coverslips and incubated for two days at 37 °C / 11 % $CO_2$. Electrophysiological recordings were carried out at DIC2.

##### Patch clamp electrophysiology

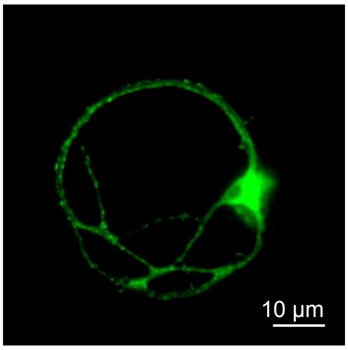

| Dunn's multiple comparisons test | Significant? | Adjusted P Value |
|---|---|---|
| TRPC5 τGFP vs. wt | No | >0.9999 |
| TRPC5 τGFP vs. TRPC tko | Yes | 0.0024 |
| TRPC5 τGFP vs. TRPC5 ko | No | 0.1092 |
| wt vs. TRPC tko | Yes | 0.0437 |
| wt vs. TRPC5 ko | No | 0.709 |
| TRPC tko vs. TRPC5 ko | No | >0.9999 |

**Figure S20** (related to **Figure 4**). **left:** fluorescence image of autaptic **5ki** neuron (GFP imaging), used to confirm the presence of TRPC5. **right**: expanded statistics for the data shown in **Figure 4ab** (one-way ANOVA post-hoc Kruskal Wallis and Dunn's multiple comparison test). Note that values listed in **Figure 4d** were mean ± SEM with Student's two-tailed t-test used instead for statistics.

For photoswitching **AzPico** (30 nM), cells were transiently illuminated on the stage of an inverted microscope (Axiovert 200, Zeiss, Germany) with light from a Polychrome 4 monochromator (Till Photonics, Germany) at the indicated wavelengths, with default slit width 10 nm.

<u>Ephys of autaptic hippocampal neurons</u>: Recording pipettes had a resistance of 4-5 MΩ. Inward currents were measured in the voltage-clamp mode at -70 mV in extracellular solution containing (in mM): 120 NaCl, 2.6 KCl, 1-2 $CaCl_2$, 2 $MgCl_2$, 20 HEPES, 30 glucose, pH 7.3, 290-300 mOsm. The intracellular patch pipette solution contained (in mM): 140 K-gluconate, 11 NaCl, 2 Mg-ATP, 0.2 $Na_2$-GTP, 1.1 EGTA, 11 HEPES, 11 glucose, 280 mOsm (pH 7.3 with NaOH). Recordings were

performed with an EPC10 amplifier (HEKA Electronic, Germany) controlled by Pulse 8.5 program (HEKA Electronic, Germany). Only cells with an access resistance of 5-15 MΩ, 60-80 % resistance compensation and a leak current of <300 pA were analysed. Ephys measurements were recorded at the digitalization rate of 20 kHz and analysed using customized routine in IgorPro (Wavemetrics, USA). Data were analysed using Igor Pro 5.0. The maximal current amplitudes were determined at the phase's plateau devoid of any spontaneous activity (thus the “460 nm” values would typically reflect a value from between 3-5 seconds into the followup 460 nm phase, while the “UV” values would typically reflect a value from the five seconds before switching back to 460 nm). Each cell's photoswitch-based current differential **$\Delta IC_\lambda$** [pA] was determined as the absolute value of its 365 nm plateau current minus its followup 460 nm plateau current. **Figure 4ab** lists mean ± SEM values; one-way ANOVA post-hoc Kruskal Wallis used for statistics (**Figure S20**).

Ephys of cultured chromaffin cells: Current recordings were performed at room temperature in voltage clamp mode using the whole cell patch clamp technique (holding potential – 70 mV). The extracellular Ringer's solution used for all electrophysiological recordings contained (in mM): 130 NaCl, 4 KCl, 2 $CaCl_2$, 1 $MgCl_2$, 30 glucose, 10 HEPES-NaOH (pH 7.3, 310 mOsm). Thick-walled borosilicate pipettes (pipette resistance 4-5 MΩ) were filled with intracellular solution containing (in mM): 75 K-glutamate, 10 HEPES, 10 NaCl, 121 glucose, 0.1 EGTA (300 mOsm, pH = 7.3 with KOH). 360 nm light was used for activation, flanked by illuminations at 470 nm which silenced the channel activity. Cell membrane capacitance (CM) was determined before and after UV activation (360 nm). Current signals were filtered at 2.9 kHz and digitized gap-free at 20 kHz prior to analysis. Data were acquired with the Pulse software (HEKA, Lambrecht, Germany) and capacitance measurements were performed by the Lindau–Neher technique (sine wave stimulus: 1000 Hz, 35 mV peak-to-peak amplitude, DC-holding potential -70 mV). To avoid misinterpretation, current values in sine wave stimulus phases used for CM determination are not displayed in **Figure 4c**.

### 6 Mouse experiments – brain tissue slices (Figure 5)

**Mice.** Adult female mice (7 - 20 weeks old) were kept under standard light/dark cycle (12:12; lights on 0600; lights off 1800) with food (Ssniff feed containing 9% fat, 24% protein, and 67% carbohydrate) and water ad libitum. Mice were maintained in IVC housing containing enrichment (nesting, bedding and other material). We used the following mouse strains: B6.Cg-7630403G23RikTg(Th-cre)1Tmd/J (RRID:IMSR_JAX:008601, referred to as Th-Cre mice)[68], B6;129S-Gt(ROSA)26Sortm95.1(CAG-GCaMP6f)Hze/J (RRID:IMSR_JAX:024105, referred to as R26-GCaMP6f or Ai95D mice)[69], B6.Cg-Gt(ROSA)26Sortm14(CAG-tdTomato)Hze/J (JR # 007914, referred to as R26-tdTomato mice)[70] and Trpc5tm1.1Lbi (RRID:IMSR_JAX:024535; MMRRC Stock No: 37349-JAX, referred to as Trpc5-E5–/– mice.[36,71] We crossed Th-Cre mice with either R26-tdTomato or R26-GCaMP6f reporter mice resulting in a strain in which all Th+ cells are identifiable either through their red fluorescence (referred to as Th-tdTomato) or through their green fluorescence (referred to as Th-GCaMP6f mice). Th-tdTomato mice were heterozygous for Cre and tdTomato. The Th-GCaMP6f mice were also crossed with Trpc5-E5–/– mice, resulting in a strain in which all Th+ cells, identifiable through their green fluorescence, are deficient for Trpc5-E5 (referred to as Th-GCaMP6f-ΔTrpc5). Th-GCaMP6f and Th-GCaMP6f-ΔTrpc5 mice were heterozygous for Cre and GCaMP6f.

Animal care and experimental procedures were performed in accordance with the guidelines established by the German Animal Welfare Act, European Communities Council Directive 2010/63/EU, the institutional ethical and animal welfare guidelines of the Saarland University (approval number of the Institutional Animal Care and Use Committee: **CIPMM-2.2.4.1.1, 2.4.1.1.-Leinders-Zufall**). The number of animals used is a minimum necessary to provide adequate data to test the hypotheses of this project. We minimized the number of animals required by the animal welfare committees wherever possible.

**Solutions and Chemicals for hypothalamic brain slices.** Oxygenated extracellular bath solution (95% $O_2$/5% $CO_2$) was prepared in ultrapure water (>18.2 MΩ-cm resistivity at 25 °C, low ppt in divalent cations) and contained (in mM): 120 NaCl (Gruessing, Germany), 25 $NaHCO_3$ (Merck, Darmstadt, Germany), 5 KCl (Gruessing, Germany), 5 *N,N*-bis(2-hydroxyethyl)-2-aminoethanesulfonic acid (BES), 1 $MgSO_4$, 1 $CaCl_2$ (Gruessing), 10 glucose (Merck); osmolarity: ~300 mOsm/kg and pH: 7.3. The **AzPico** and **AzHC** stock solutions (10 mM or 1 mM) were prepared in anhydrous DMSO. To dissolve **AzPico** and **AzHC** the solution was heated to 60 °C followed by a short period of sonication. The stock solutions of **AzPico** and **AzHC** were further diluted in the extracellular bath solution to make the final working solution. DMSO concentrations were ≤ 0.05 % (vol/vol) except for 1 µM **AzHC**. Here the DMSO concentration was ≤ 0.2 % (vol/vol). Unless stated otherwise, all chemicals were purchased from Sigma (Munich, Germany). Chemicals were of analytical or higher grade.

**Preparation of Hypothalamic Brain Tissue Slices.** All experiments were performed on coronal brain slices (Bregma -1.6 and -2.2 mm) freshly prepared from female mice adapting previously described methods.[36,72] Mice were anesthetized with 20% isoflurane (vol/vol) in propylene glycol using the open-drop method followed by decapitation. Brains were removed quickly, submerged in ice-cold extracellular bath solution, and sliced (275 µm thick) using a vibrating-blade microtome (Leica, Germany). Slices were kept at 31.5 °C for 15 min and then brought back at RT for 30 min before starting an experiment. We minimized the amount of mice wherever possible as requested by the animal welfare committee, but used at least 3 mice per genotype in independent experiments.

**$Ca^{2+}$ Imaging and Combined Laser Scanning-controlled Photoswitching.** We used an upright scanning confocal microscope (Zeiss LSM 880 Indimo) equipped with a standard Argon laser for GCAMP6f excitation at a wavelength of 488 nm and a UV laser (Coherent) emitting 355 nm for photoswitching **AzHC** and **AzPico.**[73–75] Emitted fluorescence was collected between 500 and 560 nm. All scanning head settings, e.g., frame size (512 x 256), pinhole size (16.1 µM section), pixel size (1.04 µM), and pixel dwell (2.05 µs), were kept constant during each experiment. Images were acquired at 1.7 Hz and analysed using a combination of Zen (Zeiss), ImageJ (NIH), Igor (Wavemetrics) and OriginLab (OriginLab Corporation) software.

For laser scanning-controlled photoswitching of **AzHC** or **AzPico**, the UV laser light (355 nm) coupled to the confocal microscope was focused onto the image plane through a 20 x 1.0 NA Plan-Apochromat water immersion objective (Zeiss). The depth of focus was 16 µm which ensured, together with the region of interest (ROI) diameter, illumination of individual cells. Before photoswitching, UV laser light was optimally focused using 18 µm thick brain tissue sections loaded with Hoechst 33342 (1:10000; ThermoFisher) and the semi-automated correction tool of the Zen software (Zeiss). **AzHC** or **AzPico** was added to the bath chamber containing the brain slice and subsequently incubated for about 10 min in the dark. GCaMP6f fluorescence was then measured in the presence of the photoswitchable compound using the 488 nm Argon laser (2% 25 mW). Photoswitching was achieved by directing UV laser light (10 mW) on preselected ROIs using the Zen software (Zeiss), followed by exposure to the Argon laser light to resume monitoring of GCaMP6f fluorescence and switching the **AzHC** or **AzPico** back into the inactive state. The UV exposure time of the Th+ neurons in tissue slices ranged between 23 - 133 ms for **AzHC** and 7 - 156 ms for AzPico, which could be contingent on the depth of the neurons within the brain slice. The overall mean exposure time was either 68 ms with 500 nM **AzHC** or 58 ms with 500 nM **AzPico**. To protect the ultrasensitive GaAsP photomultiplier tubes from the 355 nm light, high-speed shutters closed and opened at the beginning and end of the UV exposure. Hence, GCaMP6f fluorescence could not be collected during this time. Similar photoswitching experiments were performed in absence of photoswitchable compounds to ensure that UV laser light (10 mW) and 488 nm light (2 % 25 mW) do not directly alter cell health or produce artefacts[37,76].

Data were analyzed using a combination of Zen (Zeiss), Igor (Wavemetrics) and OriginLab (OriginLab Corporation) software. The change in GCaMP6f fluorescence was expressed as relative fluorescence changes, i.e. $\Delta F/F_0$ ($F_0$ was the average of the fluorescence values of 30 frames before stimulation). Fluorescence (F) data were normalized with the $Peak_{max}$ obtained during control measurements and plotted as the function of time (t). The $Ca^{2+}$ dynamics were quantified by calculating the area under the curve (AUC). A control AUC was taken at the beginning of each recording for 3 min ($AUC_{488}$). The AUCs after photoswitching of **AzPico** and **AzHC** were calculated from the last 3 min of the recording ($AUC_{355}$). The total duration of the recordings was 9 min. AUC data were plotted either as box or as violin plots to show the distribution and density probability of the data. The violin plots were created using Igor Pro (Wavemetrics) with a bandwidth following Scott's rules and a Gaussian kernel. The response index (RI) of **AzPico** and **AzHC** were deduced by normalizing the estimated change in AUC after photoswitching with the mean of control AUCs, and is a measure used to calculate whether a compound is stimulatory (RI >1), ineffective (RI = 0), or inhibitory (RI <0). The mean and total burst duration along with the mean frequency of $Ca^{2+}$ waves were calculated at the level of 20% of the signal amplitude of the normalized $Ca^{2+}$ waveform measured for 3 minutes under control conditions, and later compared with $Ca^{2+}$ signals measured in the last 3 minutes of recordings after the UV illumination. The mean amplitude was calculated by taking account of all the $F_{max}$ of $Ca^{2+}$ waveforms measured within 3 minutes under control conditions and after UV illumination. The change in the area under the curve ($\Delta$AUC) is the $AUC_{355}$ minus $AUC_{488}$. Through the Igor Pro software package, user-defined functions in combination with an iterative Levenberg-Marquardt nonlinear, least-squares fitting routine were applied to the data. Dose-response curves were fitted by the equation:

$$f(x) = Emin + (Emax - Emin)/\{1 + [\,EC50/x\,]^n\}$$

where x is the drug concentration, Emin the baseline response, Emax the maximal response at saturating concentrations, and EC50 the drug concentration that produces 50% of the maximal response, with slope n being the Hill coefficient of the sigmoid curve.

**Quantification and Statistical Analysis**. Statistical analyses were performed using Origin Pro 2017G (OriginLab Corporation, Northampton, MA, USA). Assumptions of normality and homogeneity of variance were tested before conducting the following statistical approaches. A paired two-tailed Student's t-test was used to measure the significance of the differences between two distributions of the same Th+ neuron. Multiple groups were compared using the Kruskal-Wallis ANOVA in combination with the Dunn's test. The probability of error level (alpha) was chosen to be 0.05. Box plots display the interquartile ranges, median (line) and mean (black rhombus) values with whiskers indicating SD values. Additional data and representations relevant to **Figure 5** are given in **Figure S21** and detailed in its caption:

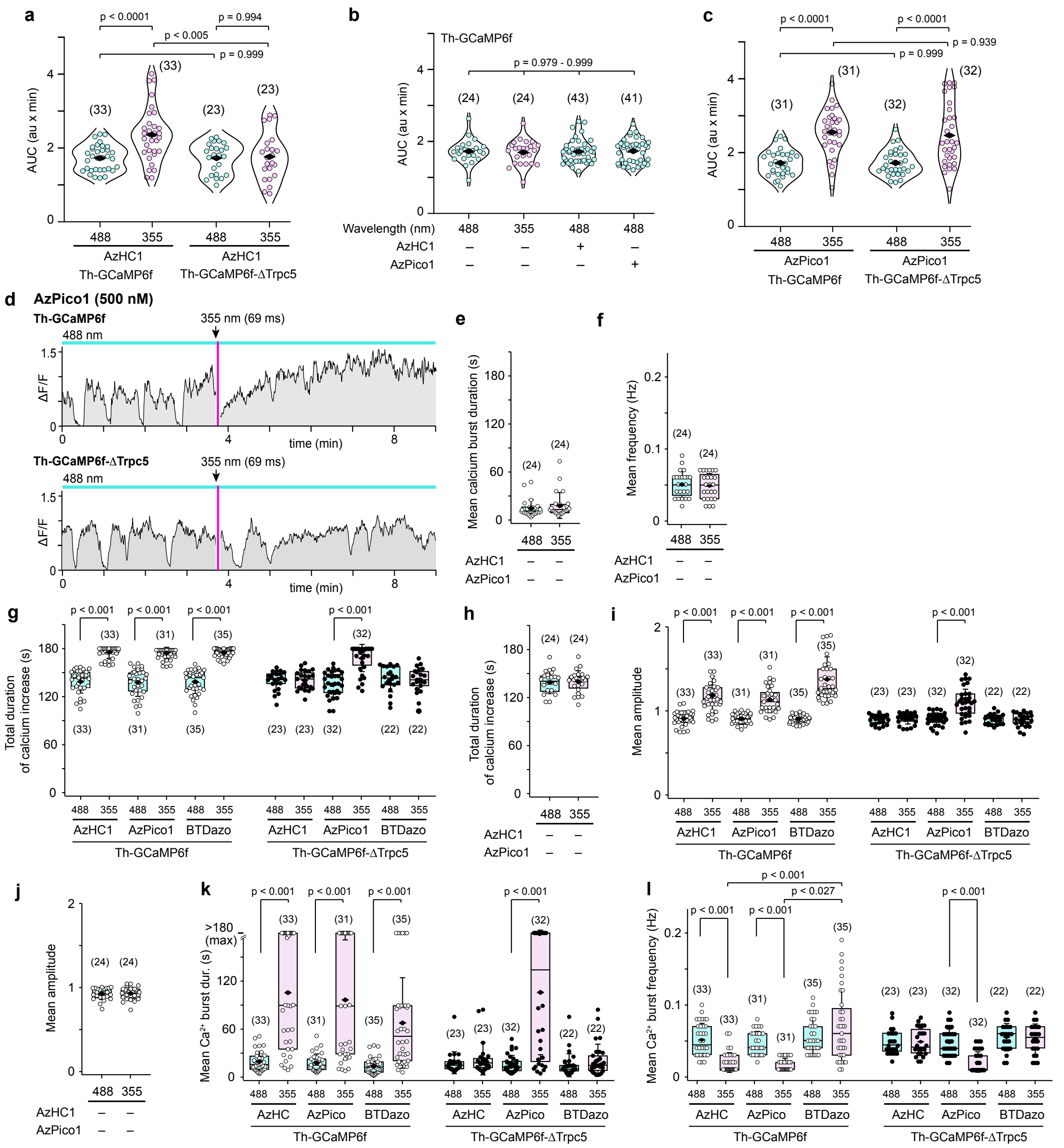

**Figure S21** (related to **Figure 5**; **note: for expanded legend to Figure 5, see Figure S27**). (**a**) Area under the curve (AUC) from $Ca^{2+}$ signals before (488 nm) and after 355 nm UV illumination of Th+ neurons treated with **AzHC** (500 nM) in arcuate hypothalamic brain slices of Th-GCaMP6f and Th-GCaMP6f-ΔTrpc5 mice. (**b**) AUC from $Ca^{2+}$ signals before (488 nm) and after 355 nm UV illumination of Th+ neurons in absence of **AzHC** and **AzPico** as well as at 488 nm illumination in presence of either **AzHC** (500 nM) or **AzPico** (500 nM). (**c**) AUC from $Ca^{2+}$ signals before (488 nm) and after 355 nm UV illumination of Th+ neurons treated with **AzPico** (500 nM) in arcuate hypothalamic brain slices of Th-GCaMP6f and Th-GCaMP6f-ΔTrpc5 mice. (**d**) Original traces of spontaneous $Ca^{2+}$ responses in Th+ neurons of a Th-GCaMP6f and a Th-GCaMP6f-ΔTrpc5 mouse treated with **AzPico** (500 nM) stimulated by *E*→*Z* isomerization of **AzPico** using a 355 nm UV laser pulse for 69 ms. (**e, f**) The mean $Ca^{2+}$ burst duration (**e**) and mean frequency (**f**) of Th+ neurons before (488 nm) and after 355 nm UV laser without photoswitchable modulators. (**g-j**) The total duration of $Ca^{2+}$ increase (**g,h**) and the mean amplitude of the $Ca^{2+}$ fluorescence signal (**i,j**) in Th+ neurons in Th-GCaMP6f and Th-GCaMP6f-ΔTrpc5 mice before (488 nm) and after 355 nm UV laser using (**g,i**) **AzHC** (500 nM), **AzPico** (500 nM) and **BTDazo** (10 µM) or (**h,j**) without photoswitchable modulators. (**k,l**) Replotting main **Figure 5** panels **g,h** with linear rather than logarithmic vertical scaling. ***Additional detail related to both Figure 5 and Figure S21***: Th+ neurons treated with 500 nM *E*-**AzHC** are unaffected by 488 nm imaging (**Fig S19b**), but a ≥23 ms exposure (mean= 68 ± 29 ms, range: 22.3 - 133.2 ms) to 355 nm light (*E*→*Z*) induced a sustained (low frequency) high-$Ca^{2+}$ signal lasting up to ≥3 min (14 of 33 cells; **Figure 5bd**). In wildtype slices, **AzPico** also induced long-lasting high $Ca^{2+}$ signals after 355 nm pulsing (13 out of 31 cells), with overall $Ca^{2+}$ signal properties that are not distinguishable from *Z*-**AzHC** (**Figure 5gh**). Figure 5 and its supplementary data are deposited in Figshare (doi:10.6084/m9.figshare.26232254).

### 7 Spontaneous motility and isometric contractility (Figure 6)

After sacrificing mice in accordance to the European (Council Directive 2010/63/EU) and German guidelines for the welfare of experimental animals, the abdomen was opened, and intestinal loops were mobilised. Ileal segments were obtained starting 10 mm from the ileocoecal junction. Ileal segments with a length of 3 mm for myography experiments or with a length of 8-12 mm for observation of spontaneous movement were cut out and flushed with carbogen-equilibrated KBRS. The adventitial layer was carefully peeled off, using a pair of forceps. The remaining intestinal segments, comprising mucosal, submucosal and muscular layers were placed in carbogen-saturated KBRS and immediately used for the experiments. Throughout these experiments, "wild-type" mice have the same genetic background as the knockout mice.

**Spontaneous motility**: for macroscopic observation, ileal segments were placed in KBRS-filled 24-well plates, and treated with 300 nM atropine (which has long been known to override the inherent oscillatory bias of the gut nerve system to paralyse digestion) or its solvent. The multiwell plate was mounted onto the FLIPR device used in cell culture studies (**Figures 1-2**) with a transmitted light source (dim white light from a fluorescent bulb) applied from the top through a diffusing sheet of white paper. A time-lapse movie was recorded, and 365 nm or 447 nm LED were electronically switched to induce photoswitching. The recorded image stack was cut into substacks, covering a single well, each. The temporal variance of the shadow cast by the segments was calculated for consecutive 1 s bins and taken as a measure to obtain the temporal signature of the motility. Gross movements of the observed ileal segments most likely indicate activity of the longitudinal muscular layer (**Figure 6bc**, **Figure S22b**, **Movie S1**).

**Myographic isometric contractility:** The contractile forces exerted by the circular layer of intestinal segments were quantitatively assessed by measurement in a 4-channel calibrated multi myograph device (DMT 620M, Danish Myo Technology, Hinnerup, Denmark), with digital data acquisition controlled by Labchart software (Version v8; ADINSTRUMENTS). Heated (37°C) bath chambers were filled with carbogen-equilibrated KBRS, and typically 3-mm ileal segments were manoeuvred over custom-made u-shaped hooks that prevented the mucosal layer from prolapsing on the edges. A pre-tension of typically 1.5-2 mN/mm was applied, and ileal segments were allowed to equilibrate to the conditions for at least 10 min in the presence of typically 1-30 nM **AzPico** or its solvent (0.5% DMSO). After renewing the pre-tension, the recording of isometric contractions was started, while applying alternating illuminations typically at 365 nm and 447 nm, or else at 385 nm and 470 nm, (typically 10 s exposure each) over typically 9-11 cycles, with no additional illumination afterwards. Data were sampled at 1 kHz, and displayed after normalisation to the length of the segments (N/m) (**Figure 6d-h**, **Figure S22c-g**).

#### 7.1 Tissue Switching Reveals the Power of the Ideal Efficacy Switch Paradigm

##### 7.1.1 First Saturate with Ligand, Then Dial the Wavelength

Even in the usually tricky situation that a targeted biological effect only occurs when a protein target is *partially but not fully stimulated* (here: physiology-like contractility as in **Figure 6e**, with the overlay of the normal oscillatory frequncy), ideal efficacy switches are *still* perfectly capable of addressing them robustly. What is possible in the rare case that only fixed operating wavelengths may be available [e.g. in **Figure 6ef**, if only 365 nm and 447 nm would be available], would be to use a sub-saturating concentration of the switch (i.e. titrating it in, just like all affinity switch approaches): this could compensate for the fixed wavelength/s being "too complete" in their switching and thus overstimulating the target to drive unwanted activity. As an example, in **Figure S22c**, the 365 nm photogenerated *Z*-rich PSS is "too efficient" at target activation when (presumably saturating) 30 nM **AzPico** is used, resulting in a rictus-like freezing of the muscle with suppression of the desired oscillatory frequency. However, we highlight that titrating an ideal efficacy switch's concentration is no more logical than "driving a Ferrari in reverse, just to reduce its maximum speed". The more reliable approach would be to install e.g. 385 nm or 395 nm LEDs, that drive higher *E/Z* ratios under UV light (c.f. **Figure 1c**), and thus be able to exploit the reproducibility of a saturating concentration of switch without over-stimulating the channel. Indeed, when multiplying the **AzPico** concentration by 10 (to 300 nM), simply altering the LED source to 385 nm allowed to avoid the over-stimulated rictus and instead capture the repeatable, physiological-like oscillatory overlay (**Figure S22f**) even under this saturating concentration.

##### 7.1.2 Ideal Efficacy Switches That Also Have High Affinity

The "ideal" efficacy switch by our definition (includes: high affinity) may benefit from the intensely practical benefit that compound wash-in can be *complete, and long-lasting*, yet tissue-level photocontrol may still be *entirely reproducible even hours later*: as long as saturation is achieved. We demonstrated this by exploiting the high affinity and presumably also hydrophobicity of **AzPico**, first exposing tissues, then exchanging for fresh medium (no **AzPico** content) for 2 hours: and the tissues remained perfectly photoreversibly operable (**Figure S22g**). The exact performance we show there may be assisted by the facts that (a) likely, far more **AzPico** is sequestered in tissues than the concentration of TRPC4/[5] channels (e.g. hydrophobicity-driven partitioning into lipid environments in cells and tissues that act as reservoirs); but potentially also (b) not every TRPC4 channel must be activated in order to obtain the desired physiological-like readout, therefore some more loss over time may be tolerated than for targets where 100% saturation must be ensured. Nonetheless, we note that the persistence time is excellently long; and that even if the degree of activation under 385 nm had reduced, one could still imagine to ***dial*** in a slightly shorter wavelength with higher *Z*-content PSS, even down to 365 nm, in order to reach maximum possible activation.

#### 7.2 Minor Remarks on Additional Experiments (Figure S22) and Prior Art

When returning to basal tension under blue light, we note that there is some temporary suppression of spontaneous oscillations under blue light and even when it is switched off; this is visible both in gross motility and in myography. Most prosaically, one could imagine a temporary desensitisation and ionic balance reset. (If only the motility data were considered, one might suggest that the block lifts after the broadband fluorescent bulb spectrum used for imaging shifts the population *E:Z* equilibrium ratio slightly towards a lower *E* proportion that is less inhibitory to intestinal motions; or, e.g., that a dynamic and TRPC4-dependent effect such as in-pocket switching is in action: but, the same effect is evident *after* 447 nm switchoff in the myography (where bulb illumination is not applied) and it seems an unnecessarily complicated hypothesis. We also note that unless the binding affinities of the *E/Z* isomers are truly identical across all bound stoichiometries, such **AzPico-**derived effects ought also to depend on the dosage applied: not matching to the robust, context-independent, photoequilibrium-driven effects that we pursue, and identify, in this paper.

Related to the issues discussed in section 7.1 above, we note that under 365 nm, when lowering or raising the **AzPico** concentration vs the typically 3 nM+365 nm that works photoreversibly as desired, divergent effects on the tonic and oscillatory forces were seen (e.g. **Figure S22c** for 30 nM+365 nm). This confirms that just 3 nM **AzPico** in the bath solution did not saturate the channel in deep tissue. However, under 447 nm, even at 30 nM which drove the rictus-like state under 365 nm, the tonic tension always dropped to baseline and oscillatory contractions always recovered within 10-30 s (**Figure S22c**, cf **Figure S22d**). This extremely fast switch-off of muscle tension under 447 nm is as notable as the robust and repeatable activation of tension under UV.

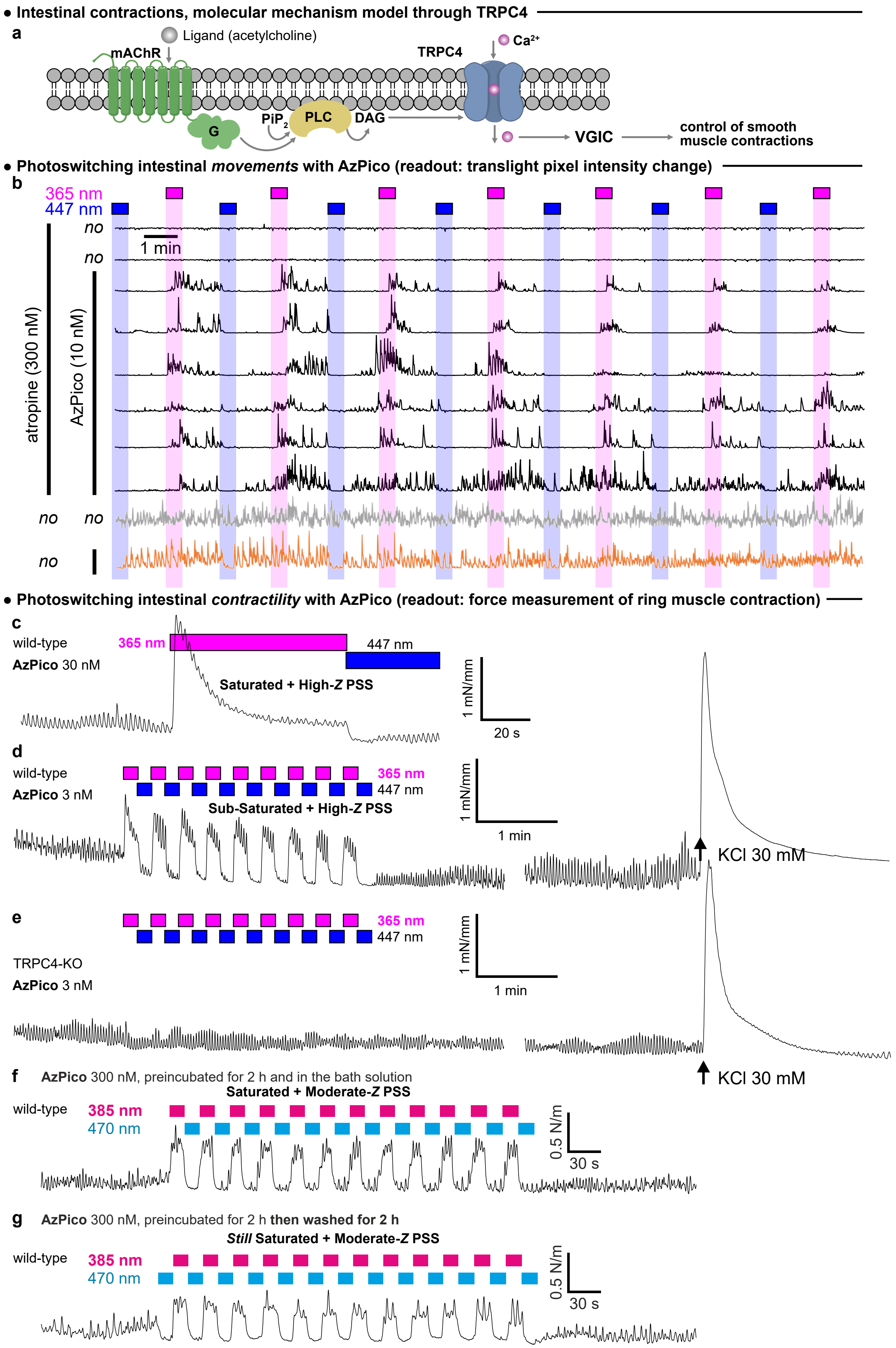

**Figure S22** (related to **Figure 6**). (**a**) **Molecular mechanism** advanced for TRPC4-dependent intestinal contractility (overlaps in content with **Figure 6a**, reproduced for reference). Physiological intestinal contractions are macroscopically coordinated, oscillatory motions. Intestinal segments have a "tonic" resting contractile force (baseline); they also propagate an inherent, slow oscillatory electrical pacemaker potential, that is amplified by classical signaling routes (e.g. via acetylcholine ACh > mAChR > phospholipase C > diacylglycerols > TRPC4 cation influx > membrane depolarisation) until it surpasses a threshold that triggers the oscillatory contractility of peristalsis. (**b**; expands on traces shown in **Figure 6c**) Additional **macroscopic motility** imaging traces (vertical axis: average change of pixel intensity for transmitted light image at time *t* compared to the previous frame, arbitrary units on a common scale (for details see **Figure S28**). The photoreproducibility and photoreversibility of the macroscopic motions triggered despite atropine blockade by **AzPico** *E*→*Z* switching (UV light) are striking. There is no photoresponse in the absence of **AzPico** (top traces and second-from-bottom trace). In the absence of atropine blockade, tonic motility is significant (second trace from bottom); and **AzPico** *E*→*Z* photoswitching has no apparent additional effect, however, especially initial *Z*→*E* phases seem to induce a temporary block on intestinal motility during the time of active 447 nm light application (bottom trace) however, we believe this is better understood as the typical time window for *E*-suppression of spontaneous oscillations (see section 7.2). **(c-e**; expands on data shown in **Figure 6d-f) Myography traces** of ring muscle contraction. (**c**) At high **AzPico** concentration (presumed saturating), under 365 nm the tonic tension increase is larger, and the amplitude of the oscillations is somewhat decreased. (**d**) At low **AzPico** concentrations (3 nM), under 365 nm the *amplitude* of the contractile oscillations increases but their tonic tension does not change; the KCl spike establishes a reference for muscle contractility, and tension drop over time. (**e**) Segments from TRPC4-deficient mice never responded to *E/Z*-**AzPico**. (**f**) Similar experiments as in panels **c-d**, but with vast excess of **AzPico**, can deliver fully photoreversible physiological-type contractility when avoiding the *Z*-maximised PSS of 365 nm (**c**) in favour of a *Z*-moderated PSS at 385 nm: a consequence of the ideal efficacy switch paradigm. (**g**) The high affinity and presumably also hydrophobicity of **AzPico** meant that tissues could be exposed, then the medium replaced (no **AzPico** content), but tissues remain perfectly photoreversibly operable for long times afterwards (here, 2 h).

Bon and Zholos[77] published the conclusion of TRPC4-dependent intestinal contractility on the basis of indirect experiments with the classical drug Pico145, with onset of effect taking minutes, and being irreversible after application (no un-blocking of motility or cycles of effect). While they could use **EA** and carbachol (target: mAChR) to record currents in isolated myocytes, for whole tissue experiments only carbachol was used to activate the intestine segments, with optional **Pico145** to block them: which, essentially, pharmacologically reproduces the effects of TRPC4 genetic knockout[78]. TRPC4 was however never *directly* activated in tissues before, and its effective and reproducible demonstration in this paper may thus have novelty as well as practical value.

#### 7.3 Why is AzPico/AzHC-like efficacy photoswitching useful *in vivo* in practice?

We conclude by briefly recalling the main end-user advantages of **AzPico** and **AzHC**, noting also that these ought to also apply to *any other high-potency ideal efficacy switches, for other targets,* created according to the paradigm in **Figure 1**. Their paradigm allows easy spatiotemporal targeting, through reversible, rapid, non-invasive modulation using light; they can fully reproducibly apply the same *levels of applied bioactivity* (c.f. baseline, physiologically stimulated, or overstimulated, as shown in **Figure 6e-g**) regardless of local or inter-assay concentration variations (that are otherwise significant and arise from a mixture of ADME-PK, technical handling, inter-animal differences, different distances of target cells from a reservoir or blood vessel, or assay timescales, etc: see section **7.1.1**) as long as appropriate fixed wavelength/s are dialled in (i.e. it is convenient to apply); unlike slow wash-in/wash-out rates, their photoswitch mechanism acts much faster than tissue desensitisation, thus allowing multiple cycles of on-/off- switching in tissues (e.g. in each segment of the intestine), for statistics and reproducibility; and photoswitching also acts as an effective internal control e.g. in neighbouring cells in the same tissue slice, or by considering UV-vs-blue activity.

### 8 Chemistry

#### 8.1 Materials and Methods - Chemistry

**Reagents and Conditions.** Unless stated otherwise, (1) all reactions and characterisations were performed with unpurified, undried, non-degassed solvents and reagents, used as obtained, under closed air atmosphere without special precautions; (2) “hexane” used for chromatography was distilled from commercial crude isohexane fraction by rotary evaporation; (3) “column” and “chromatography” refer to manual flash column chromatography on Merck silica gel Si-60 (40−63 µm); (4) “MPLC” refers to flash column chromatography purification on a Biotage Selekt system, using prepacked silica cartridges purchased from Biotage; (5) procedures and yields are unoptimized; (6) yields refer to isolated chromatographically and spectroscopically pure materials; (7) all eluent and solvent mixtures are given as volume ratios unless otherwise specified. (8) Thin-layer chromatography (TLC) was run on 0.25 mm Merck silica gel plates (60, F-254). UV light (254 nm) was used as a visualising agent.

**Nuclear magnetic resonance (NMR) spectroscopy.** Standard NMR characterisation was by $^{1}H$- and $^{13}C$-NMR spectra on a Bruker Ascend 400 (400 MHz & 101 MHz for $^{1}H$ and $^{13}C$ respectively) and on a Bruker Ascend 500 (500 MHz & 126 MHz for $^{1}H$ and $^{13}C$ respectively). Chemical shifts (δ) are reported in ppm calibrated to residual non-perdeuterated solvent as an internal reference. Peak descriptions singlet (s), doublet (d), triplet (t), quartet (q), multiplet (m) and broad (br) are used. NMR spectra are given in **Part 10**.

**High resolution mass spectrometry (HRMS).** HRMS was carried out by the Zentrale Analytik of the LMU Munich using ESI ionisation on a Thermo Finnigan LTQ FT Ultra Fourier Transform Ion Cyclotron Resonance Spectrometer.

**High Performance liquid chromatography (HPLC) coupled with mass spectrometry (MS).** Analytical HPLC-MS was performed on an Agilent 1100 SL with (a) a binary pump to deliver H2O:MeCN eluent mixtures containing 0.1% formic acid at a 0.4 mL/min flow rate, (b) YMC-Triart C18 column (3.0 µm; 50 mm × 3 mm) maintained at 40 °C (c) an Agilent 1100 series diode array detector, (d) an Agilent LC/MSD iQ mass spectrometer. Typical run conditions were a linear gradient of H2O:MeCN from 90:10 to 0:100 (first 5 min), then 0:100 for 2 min for flushing; then the column was (re)equilibrated with 90:10 eluent mixture for 2 min.

#### 8.2 Chemical Synthesis Overview

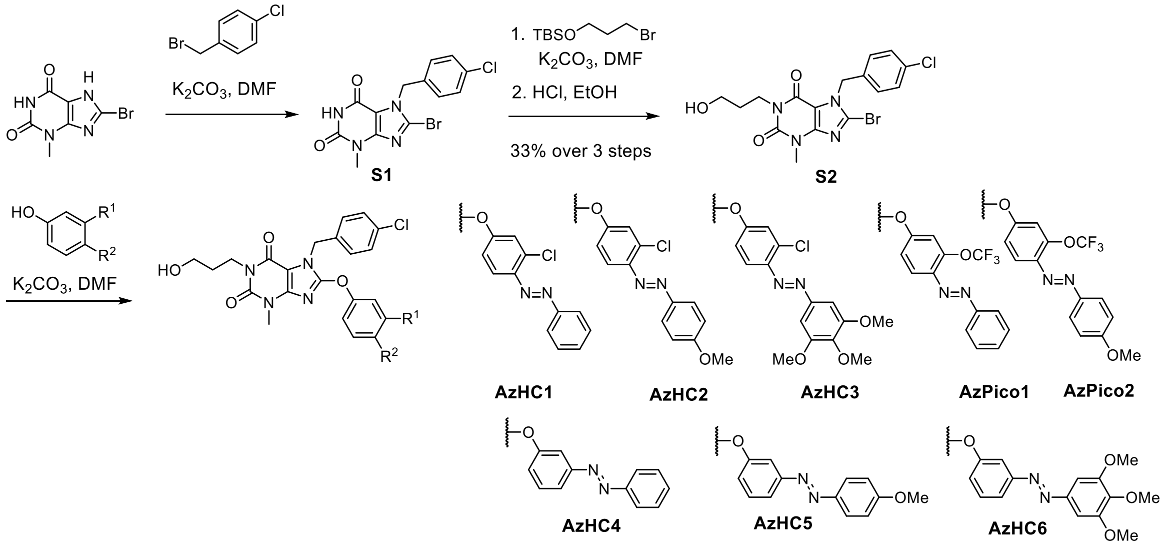

**Scheme S1**: Synthetic overview of **AzPico**s & **AzHC**s

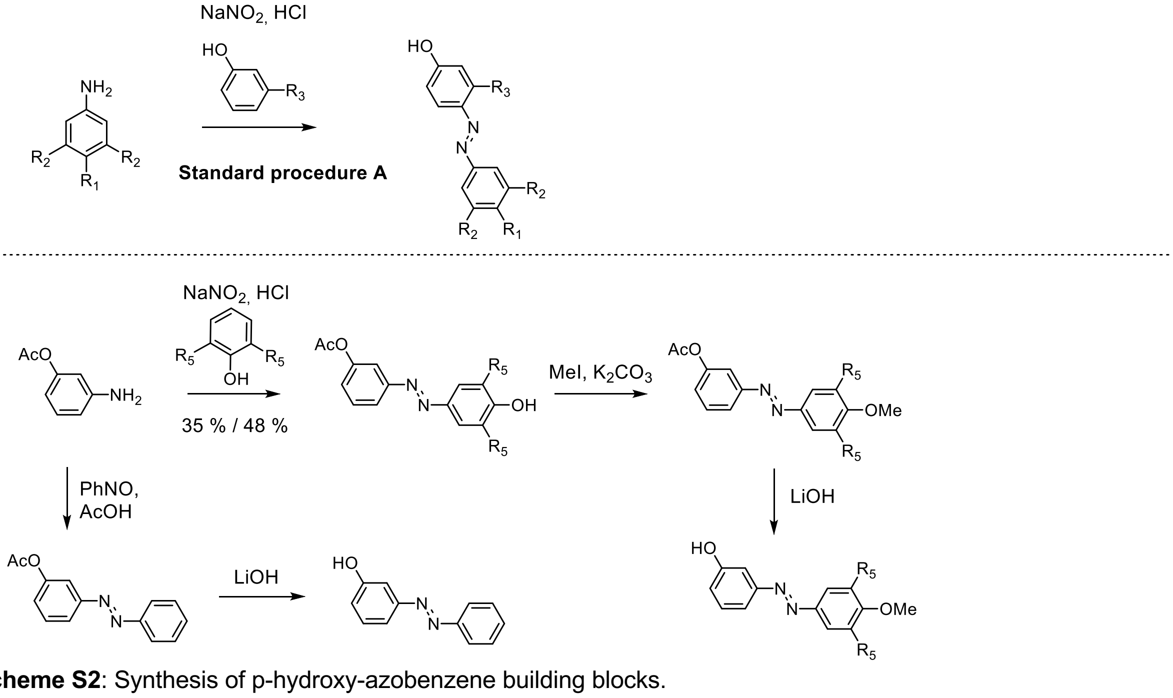

**Scheme S2**: Synthesis of p-hydroxy-azobenzene building blocks.

#### 8.3 Standard Synthetic Procedures

##### General Procedure A

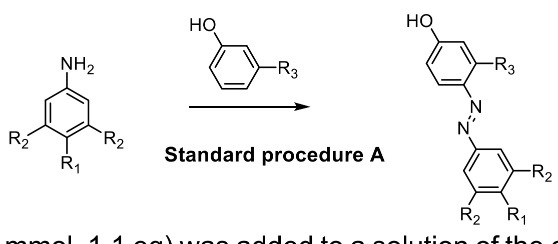

$NaNO_2$ (1.10 ml, 2 M, 2.20 mmol, 1.1 eq) was added to a solution of the corresponding aniline (2.00 mmol, 1.0 eq) in Hydrochloric acid (4 ml, 2M) and MeOH (4 ml) at 0 °C. The reaction mixture was allowed to stir for 15 min at 0 °C and was subsequently added to a solution of the phenol (2.0 mmol,

1.0 eq) in MeOH (6 ml) and Buffer (6 ml of 0.5 M $K_2HPO_4$ and 2 ml of 1 M KOH). The pH was adjusted to 9 - 11 and the reaction was stirred for 45 min at 0°C. The reaction was quenched by addition of st. $NH_4Cl$, and the mixture was extracted with EA (3 x 100 ml). The combined organic layers were dried over anhydrous $Na_2SO_4$, filtered, and concentrated. The crude product was purified by flash chromatography. If not stated otherwise the reaction was carried out in 2 mmol scale.

#### 8.4 Synthesis of Building Blocks

**8-bromo-7-(4-chlorobenzyl)-3-methyl-3,7-dihydro-1H-purine-2,6-dione (S1)**

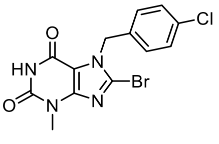

*Compound is lit. known but no NMR spectra were provided.*[1,9]

The compound was prepared according to a literature procedure:[9] A flask was charged with 8-bromo-3-methylxanthine (1.23 g, 5 mmol, 1.0 eq), 4-Chlorobenzyl bromide (1.03 g, 5 mmol, 1.0 eq), potassium carbonate (1.04 g, 7.5 mmol, 1.5 eq) and DMF (50 ml) and the mixture was stirred at 45°C for 90 min. The mixture was cooled down, partitioned between brine (200 ml) and EtOAc (200 ml), the organic layer was washed 2x with brine (100 ml), dried over $Na_2SO_4$, filtered and concentrated. The crude product was purified by flash chromatography (Gradient: Hexane/EtOAc, 8:2 to 1:1) to afford **S1** (1.37 g, 3.71 mmol, 74 %) as colourless solid.

**$^1$H NMR** (400 MHz, DMSO-*d6*) δ [ppm]: 7.42 (d, *J* = 8.5 Hz, 2H), 7.28 (d, *J* = 8.4 Hz, 2H), 5.49 (s, 2H), 3.29 (s, 3H). **$^{13}$C NMR** (101 MHz, DMSO-*d6*) δ [ppm]: 157.03, 152.90, 149.24, 135.10, 132.48, 129.18, 128.69, 126.10, 109.43, 48.32, 28.53.

**8-bromo-7-(4-chlorobenzyl)-1-(3-hydroxypropyl)-3-methyl-3,7-dihydro-1H-purine-2,6-dione (S2)**

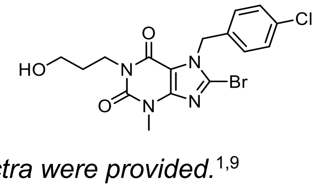

*Compound is lit. known but no NMR spectra were provided.*[1,9]

A flask was charged with **S1** (350 mg, 0.95 mmol, 1.0 eq), (3-Bromopropoxy)(tert-butyl)dimethylsilane (218 µl, 0.95 mmol, 1.0 eq), potassium carbonate (196 mg, 1.42 mmol, 1.5 eq) and DMF (5 ml) and the mixture was stirred at 90°C for 5 h. The mixture was cooled down, 6M HCl (3.16 ml, 18.9 mmol, 20 eq) was added carefully and the mixture was stirred for further 90 min at 90°C. The mixture was cooled down again, neutralized with st. $NaHCO_3$ and partitioned between $H_2O$ (50 ml) and EtOAc (50 ml), the organic layer was washed 2x with brine (500 ml), dried over $Na_2SO_4$, filtered and concentrated. The crude product was purified by flash chromatography (Gradient: Hexane/EtOAc, 1:1 to 100 % EtOAc) to afford **S2** (234 mg, 0.56 mmol, 59 %) as colourless solid.

**$^1$H NMR** (400 MHz, DMSO-*d6*) δ [ppm]: 7.43 (d, *J* = 8.5 Hz, 2H), 7.28 (d, *J* = 8.5 Hz, 2H), 5.51 (s, 2H), 4.45 (t, *J* = 5.2 Hz, 1H), 3.98 – 3.85 (m, 2H), 3.50 – 3.38 (m, 5H), 1.74 – 1.60 (m, 2H). **$^{13}$C NMR** (101 MHz, DMSO-*d6*) δ [ppm]: 153.61, 150.44, 147.97, 134.63, 132.64, 129.06, 128.78, 128.29, 108.33, 58.81, 48.74, 38.59, 30.88, 29.55.

**3-chloro-4-(phenyldiazenyl)phenol (S3)**

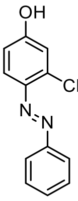

Prepared according to **General Procedure A** using aniline (184 µl 2 mmol) and 3-chlorophenol (211 µl, 2 mmol) Purification: hexane/EtOAc, 5:95 to 65:35; **S3** (382 mg, 1.64 mmol, 82 %) was obtained as red solid.

**Rf** = 0.36 hexane/EtOAc, 8:2. **$^1$H NMR** (400 MHz, $CDCl_3$) δ [ppm]: 7.96 – 7.90 (m, 2H), 7.74 (d, *J* = 8.9 Hz, 1H), 7.56 – 7.43 (m, 3H), 7.04 (d, *J* = 2.7 Hz, 1H), 6.80 (dd, *J* = 8.9, 2.7 Hz, 1H), 5.40 (s, 1H). **$^{13}$C NMR** (101 MHz, $CDCl_3$) δ[ppm]: 158.52, 152.93, 143.21, 137.52, 131.14, 129.26, 123.22, 118.99, 117.17, 114.92.

**3-chloro-4-((4-methoxyphenyl)diazenyl)phenol (S4)**

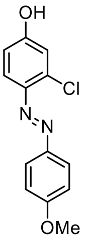

Prepared according to **General Procedure A** using p-anisidine (246 mg 2 mmol) and 3-chlorophenol (232 µl, 2.2 mmol) Purification: hexane/EtOAc, 5:95 to 65:35. **S4** (218 mg, 0.83 mmol, 42 %) was obtained as brown solid.

**Rf** = 0.40 hexane/EtOAc, 8:2. **$^1$H NMR** (400 MHz, $CDCl_3$) δ [ppm]: 7.92 (d, *J* = 9.0 Hz, 2H), 7.69 (d, *J* = 8.8 Hz, 1H), 7.06 – 6.94 (m, 3H), 6.78 (dd, *J* = 8.9, 2.7 Hz, 1H), 3.89 (s, 3H). **$^{13}$C NMR** (101 MHz, $CDCl_3$) δ [ppm]: 162.15, 158.11, 147.39, 143.18, 136.74, 125.08, 118.85, 117.09, 114.90, 114.39, 55.74.

**3-chloro-4-((3,4,5-trimethoxyphenyl)diazenyl)phenol (S5)**

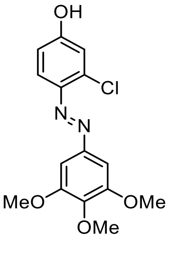

Prepared according to **General Procedure A** using 3,4,5-trimethoxyaniline (183 mg, 1 mmol) and 3-chlorophenol (116 µl, 1.1 mmol) Purification: hexane/EtOAc, 5:95 to 65:35. **S5** (260 mg, 0.81 mmol, 81 %) was obtained as orange solid.

**Rf** = 0.45 hexane/EtOAc, 8:2. **$^1$H NMR** (400 MHz, DMSO-*d6*) δ 7.67 (d, *J* = 9.0 Hz, 1H), 7.21 (s, 2H), 7.04 (d, *J* = 2.5 Hz, 1H), 6.86 (dd, *J* = 9.0, 2.6 Hz, 1H), 3.87 (s, 6H), 3.75 (s, 3H).

**4-(phenyldiazenyl)-3-(trifluoromethoxy)phenol (S6)**

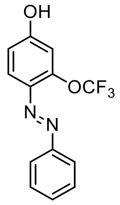

Prepared according to **General Procedure A** using aniline (92 µl, 1 mmol) and 3-(Trifluoromethoxy)phenol (117 µl, 0.9 mmol) Purification: hexane/EtOAc, 10:90 to 60:40. **S6** (183 mg, 0.65 mmol, 65 %) was obtained as orange solid.

**Rf** = 0.38 hexane/EtOAc, 8:2. **$^1$H NMR** (500 MHz, $CDCl_3$) δ [ppm]: 7.95 – 7.89 (m, 2H), 7.81 (d, *J* = 8.9 Hz, 1H), 7.54 – 7.44 (m, 3H), 6.93 – 6.91 (m, 1H), 6.86 (dd, *J* = 8.9, 2.6 Hz, 1H). **$^{13}$C NMR** (126 MHz, $CDCl_3$) δ[ppm]: 159.12, 152.78, 148.50, 139.34, 131.28, 129.28, 123.17, 120.72 (d, *J* = 258.3 Hz)., 118.93, 114.88, 109.96. **$^{19}$F NMR** (471 MHz, $CDCl_3$) δ [ppm]: -57.40.

*Note on $^{13}$C spectrum: $CF_3$ quartet was detected as doublet.*

**4-((4-methoxyphenyl)diazenyl)-3-(trifluoromethoxy)phenol (S7)**

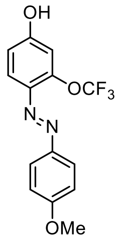

Prepared according to **General Procedure A** using p-anisidine (246 mg, 2 mmol) and 3-(Trifluoromethoxy)phenol (286 µl, 2.2 mmol) Purification: hexane/EtOAc, 10:90 to 60:40. **S6** (225 mg, 0.72 mmol, 36 %) was obtained as brown solid.

**Rf** = 0.42 hexane/EtOAc, 8:2. **$^1$H NMR** (400 MHz, $CDCl_3$) δ [ppm]: 7.95 – 7.88 (m, 2H), 7.76 (d, *J* = 8.9 Hz, 1H), 7.04 – 6.98 (m, 2H), 6.89 (dd, *J* = 2.7, 1.3 Hz, 1H), 6.83 (dd, *J* = 8.9, 2.6 Hz, 1H), 3.89 (s, 3H). **$^{13}$C NMR** (101 MHz, $CDCl_3$) δ[ppm]: 162.24, 158.50, 147.93, 147.40, 139.45, 125.05, 120.73 (d, *J* = 257.9 Hz), 118.71, 114.85, 114.39, 109.91, 55.74.

*Note on $^{13}$C spectrum: $CF_3$ quartet was detected as doublet.*

**3-aminophenyl acetate (S8)**

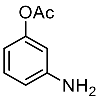

Prepared according to a literature procedure.[79]

Acetic anhydride (0.47 mL, 5.0 mmol, 1.0 eq) and $Et_3N$ (0.75 ml, 5.5 mmol, 1.1 eq) were added to a solution of 3-Aminophenol (546 mg, 5.0 mmol, 1.0 eq) in $CH_2Cl_2$ (50 ml) and stirred for 20 h at room temperature. The mixture was partitioned between EtOAc (200 ml) and st $Na_2CO_3$ (200 ml) solution, the organic layer washed with st $NH_4Cl$ (200 ml) and brine (200 ml) then dried over $Na_2SO_4$, filtered and concentrated to obtain **S8** (695 mg, 4.6 mmol, 92%) as brown oil.

**$^1$H NMR** (400 MHz, $CDCl_3$) δ[ppm]: 7.13 (dd, *J* = 8.0 Hz, 1H), 6.55 (ddd, *J* = 8.1, 2.2, 0.9 Hz, 1H), 6.48 (ddd, *J* = 8.0, 2.2, 0.9 Hz, 1H), 6.43 (dd, *J* = 2.2 Hz, 1H), 3.89 (br, 2H), 2.27 (s, 3H).

Spectroscopic data match literature data.[79]

**3-((4-hydroxyphenyl)diazenyl)phenyl acetate (S9)**

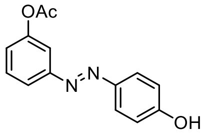

Prepared according to **General Procedure A** using aniline **S8** (130 mg, 0.86 mmol, 1.0 eq) and phenol (81 mg, 0.86 mmol, 1.0 eq). Purification: hexane/EtOAc, 10:90 to 60:40. **S9** (106 mg, 0.41 mmol, 48 %) was obtained as orange solid.

**$^1$H NMR** (400 MHz, $CDCl_3$) δ 7.87 – 7.80 (m, 2H), 7.78 (ddd, *J* = 7.9, 1.8, 1.0 Hz, 1H), 7.60 (t, *J* = 2.1 Hz, 1H), 7.51 (t, *J* = 8.0 Hz, 1H), 7.17 (ddd, *J* = 8.0, 2.4, 1.0 Hz, 1H), 6.90 (d, *J* = 8.8 Hz, 2H), 5.56 (br, 1H), 2.35 (s, 3H). **HPLC-MS** rt = 6.88 min; ESI (m/z): $[M+H]^+$ = 256.9.

**3-((4-hydroxy-3,5-dimethoxyphenyl)diazenyl)phenyl acetate (S10)**

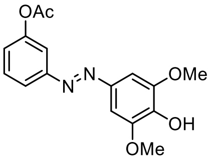

Prepared according to **General Procedure A** using aniline **S8** (130 mg, 0.86 mmol, 1.0 eq) and 2,6-dimethoxyphenol (133 mg, 0.86 mmol, 1.0 eq). Purification: hexane/EtOAc, 10:90 to 60:40. **S9** (96 mg, 0.30 mmol, 35 %) was obtained as red oil.

**$^1$H NMR** (400 MHz, $CDCl_3$) δ 7.79 (ddd, *J* = 8.0, 1.9, 1.0 Hz, 1H), 7.64 (t, *J* = 2.1 Hz, 1H), 7.51 (t, *J* = 8.0 Hz, 1H), 7.29 (s, 2H), 7.17 (ddd, *J* = 8.1, 2.3, 1.0 Hz, 1H), 3.98 (s, 6H), 2.33 (s, 3H). **HPLC-MS** rt = 6.96 min; ESI (m/z): $[M+H]^+$ = 317.0.

**3-((4-methoxyphenyl)diazenyl)phenyl acetate (S11)**

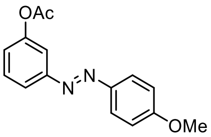

A vial was charged with **S9** (80 mg, 0.31 mmol, 1.0 eq), $K_2CO_3$ (65 mg, 0.47 mmol, 1.5 eq), acetone (3 ml) and iodomethane (20 µl, 0.33 mmol, 1.05 eq), sealed and stirred for 16h at room temperature. The mixture was partitioned between water (25 ml) and EtOAc (25 ml), the organic layer was washed with brine (2x 25 ml), dried over $Na_2SO_4$, filtered and concentrated. The crude product was purified by flash chromatography (Gradient: Hexane 100% to hexane/EtOAc, 3:7) to afford **S11** (47 mg, 0.17 mmol, 56 %) as red solid.

**HPLC-MS** rt = 7.71 min; ESI (m/z): $[M+H]^+$ = 271.0.

**3-((3,4,5-trimethoxyphenyl)diazenyl)phenyl acetate (S12)**

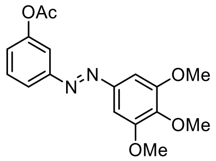

A vial was charged with **S10** (80 mg, 0.25 mmol, 1.0 eq), $K_2CO_3$ (52 mg, 0.38 mmol, 1.5 eq), acetone (3 ml) and iodomethane (17 µl, 0.27 mmol, 1.05 eq), sealed and stirred for 16h at room temperature. TLC monitoring revealed incomplete conversion and further $K_2CO_3$ (52 mg, 0.38 mmol, 1.5 eq), and iodomethane (17 µl, 0.27 mmol, 1.05 eq) were added to the reaction, and the mixture was stirred for further 8h. The mixture was partitioned between water (25 ml) and EtOAc (25 ml), the organic layer was washed with brine (2x 25 ml), dried over $Na_2SO_4$, filtered and concentrated. The crude product was purified by flash chromatography (Gradient: hexane/EtOAc, 1:0 to 3:7) to afford **S12** (63 mg, 0.19 mmol, 75 %) as red solid.

**HPLC-MS** rt = 7.58 min; ESI (m/z): $[M+H]^+$ = 331.0.

**3-(phenyldiazenyl)phenol (S13)**

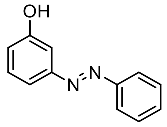

The procedure was conducted according to a literature procedure.[80]

A mixture of 3-aminophenol (109 mg, 1.0 mmol, 1.0 eq), nitrosobenzene (107 mg, 1.0 mmol, 1.0 eq) and acetic acid (5 ml) was stirred at room temperature. After 16 h the mixture was neutralized with st. $NaHCO_3$, extracted with EtOAc (50 ml) and the organic layer was washed with brine (2x 50 ml), dried over $Na_2SO_4$, filtered and concentrated. The crude product was purified by flash chromatography (Gradient: Hexane 100% to hexane/EtOAc, 4:6) to afford **S13** (22 mg, 0.11 mmol, 11 %) as brown oil.

**$^1$H NMR** (400 MHz, CDCl3) δ [ppm]: 7.97 – 7.78 (m, 2H), 7.59 – 7.44 (m, 4H), 7.42 – 7.29 (m, 2H), 7.10 – 6.84 (m, 1H), 5.15 (s, 1H). **HPLC-MS** rt = 7.10 min; ESI (m/z): $[M+H]^+$ = 198.9.

Spectral data match literature data.[80]

**3-((4-methoxyphenyl)diazenyl)phenol (S14)**

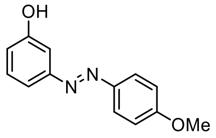

KOH (146 mg, 2.6 mmol, 15 eq) was added to a solution of **S11** (47 mg, 0.17 mmol, 1.0 eq) in MeOH (2 ml) and water (0.2 ml). The reaction mixture was stirred for 14 h, quenched by addition of st $NHCl_4$ (20 ml), extracted with EtOAc (20 ml), the organic layer was washed with brine (2x 20 ml), dried over $Na_2SO_4$, filtered and concentrated. The crude product was used immediately without further purification.

**HPLC-MS** rt = 7.10 min; ESI (m/z): $[M+H]^+$ = 229.0.

**3-((3,4,5-trimethoxyphenyl)diazenyl)phenol (S15)**

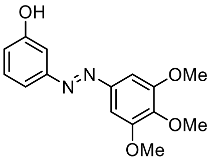

KOH (161 mg, 2.9 mmol, 15 eq) was added to a solution of **S12** (63 mg, 0.19 mmol, 1.0 eq) in MeOH (2 ml) and water (0.2 ml). The reaction mixture was stirred for 14 h, quenched by addition of st $NHCl_4$ (20 ml), extracted with EtOAc (20 ml), the organic layer was washed with brine (2x 20 ml), dried over $Na_2SO_4$, filtered and concentrated. The crude product was used immediately without further purification.

**HPLC-MS** rt = 6.88 min; ESI (m/z): $[M+H]^+$ = 289.0.

#### 8.5 Synthesis of AzPicos & AzHCs

##### AzPico1

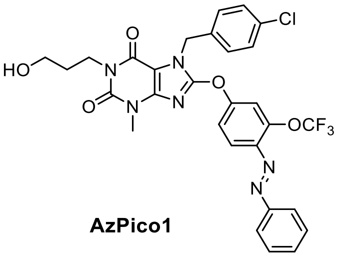

**AzPico1**

A vial was charged with **S2** (51 mg, 0.12 mmol, 1.0 eq), **S6** (34 mg, 0.12 mmol, 1.0 eq), $K_2CO_3$ (33 mg, 0.24 mmol, 2.0 eq) and DMF (1.2 ml), sealed and the reaction was stirred at 80°C. After 5 h the reaction was cooled to room temperature, $H_2O$ (2.4 ml) was added, and after 30 min the precipitate was collected by filtration. The crude product was recrystallized from EtOH to afford **AzPico1** (25 mg, 0.040 mmol 33 %) as yellow solid.

**$^1$H NMR** (500 MHz, DMSO-*d6*) δ [ppm]: 7.96 – 7.82 (m, 4H), 7.73 – 7.56 (m, 4H), 7.49 – 7.40 (m, 4H), 5.47 (s, 2H), 4.46 (t, *J* = 5.2 Hz, 1H), 4.00 – 3.85 (m, 2H), 3.54 – 3.38 (m, 2H), 3.34 (s, 3H), 1.84 – 1.63 (m, 2H). **$^{13}$C NMR** (126 MHz, DMSO-*d6*) δ [ppm]: 155.43, 153.97, 152.05, 151.65, 150.57, 146.03, 145.41, 141.76, 135.18, 132.68, 132.45, 129.72, 129.65, 129.60, 128.77, 122.83, 120.12 (d, *J* = 257.8 Hz), 119.82, 118.71, 115.06, 102.81, 58.82, 46.02, 38.40, 31.01, 29.55. **$^{19}$F NMR** (471 MHz, DMSO-*d6*) δ [ppm]: -56.75. **HRMS** (ESI, m/z): $[M+H]^+$ calcd for $C_{29}H_{25}ClF_3N_6O_5$: 629.1522; found: 629.1517

*Note on $^{13}$C spectrum: $CF_3$ quartet was detected as doublet.*

##### AzHC1

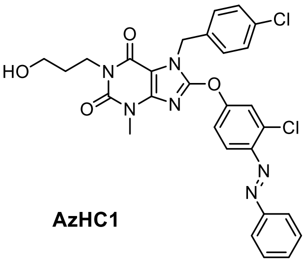

**AzHC1**

A vial was charged with **S2** (50 mg, 0.12 mmol, 1.0 eq), **S3** (27 mg, 0.12 mmol, 1.0 eq), $K_2CO_3$ (32 mg, 0.23 mmol, 2.0 eq) and DMF (1 ml), sealed and the reaction was stirred at 80°C. After 5 h the reaction was cooled to room temperature, $H_2O$ (2 ml) was added, and after 30 min the precipitate was collected by filtration. The crude product was recrystallized from EtOH to afford **AzHC1** (36 mg, 0.062 mmol 53 %) as orange solid.

**$^1$H NMR** (400 MHz, DMSO-*d6*) δ [ppm]: 7.97 – 7.90 (m, 2H), 7.83 (d, *J* = 2.6 Hz, 1H), 7.79 (d, *J* = 9.0 Hz, 1H), 7.66 – 7.59 (m, 3H), 7.52 (dd, *J* = 9.0, 2.6 Hz, 1H), 7.47 – 7.40 (m, 4H), 5.46 (s, 2H), 4.46 (t, *J* = 5.2 Hz, 1H), 4.02 – 3.86 (m, 2H), 3.50 – 3.39 (m, 2H), 3.32 (s, 3H), 1.76 – 1.66 (m, 2H). **$^{13}$C NMR** (126 MHz, DMSO) δ[ppm]: 155.28, 153.96, 152.03, 151.76, 150.56, 145.46, 145.36, 135.17, 135.02, 132.68, 132.29, 129.67, 129.64, 128.78, 122.96, 121.46, 119.42, 118.78, 102.79, 58.82, 46.01, 38.38, 31.01, 29.64. **HRMS** (ESI, m/z): $[M+H]^+$ calcd for $C_{28}H_{25}Cl_2N_6O_4$: 579.1309; found: 579.1302

**AzPico2**

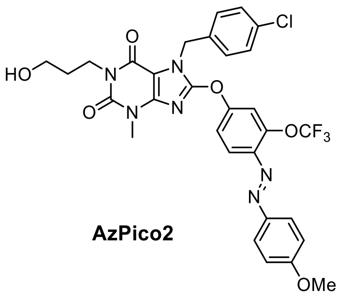

AzPico2

A vial was charged with **S2** (50 mg, 0.12 mmol, 1.0 eq), **S7** (37 mg, 0.12 mmol, 1.0 eq), $K_2CO_3$ (32 mg, 0.23 mmol, 2.0 eq) and DMF (1.2 ml), sealed and the reaction was stirred at 80°C. After 5 h the reaction was cooled to room temperature, $H_2O$ (2.4 ml) was added, and after 30 min the precipitate was collected by filtration. The crude product was recrystallized from EtOH to afford **AzPico2** (32 mg, 0.048 mmol 49 %) as yellow solid.

**$^1$H NMR** (500 MHz, DMSO-*d6*) δ [ppm]: 7.91 (d, *J* = 9.0 Hz, 2H), 7.85 (d, *J* = 9.0 Hz, 1H), 7.82 – 7.79 (m, 1H), 7.58 (dd, *J* = 9.0, 2.6 Hz, 1H), 7.47 – 7.41 (m, 4H), 7.18 (d, *J* = 9.0 Hz, 2H), 5.46 (s, 2H), 4.46 (t, *J* = 5.2 Hz, 1H), 3.97 – 3.92 (m, 2H), 3.49 – 3.41 (m, 2H), 3.32 (s, 3H), 1.76 – 1.65 (m, 2H). **$^{13}$C NMR** (126 MHz, DMSO-*d6*) δ [ppm]: 162.78, 154.73, 153.95, 151.77, 150.56, 146.44, 145.55, 145.41, 141.94, 135.19, 132.67, 129.64, 128.76, 125.02, 120.13 (d, *J* = 258.2 Hz), 119.79, 118.55, 115.01, 114.90, 102.76, 58.82, 55.78, 45.99, 38.38, 31.01, 29.54. **HRMS** (ESI, m/z): $[M+H]^+$ calcd for $C_{30}H_{27}ClF_3N_6O_6$: 659.1627; found: 659.1613.

*Note on $^{13}$C spectrum: $CF_3$ quartet was detected as doublet.*

**AzHC2**

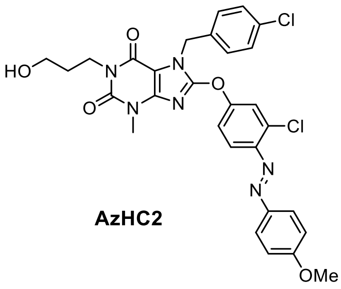

AzHC2

A vial was charged with **S2** (50 mg, 0.12 mmol, 1.0 eq), **S4** (27 mg, 0.12 mmol, 1.0 eq), $K_2CO_3$ (32 mg, 0.23 mmol, 2.0 eq) and DMF (1 ml), sealed and the reaction was stirred at 80°C. After 5 h the reaction was cooled to room temperature, $H_2O$ (2 ml) was added, and after 30 min the precipitate was collected by filtration. The crude product was recrystallized from EtOH to afford **AzHC2** (61 mg, 0.10 mmol 86 %) as yellow solid.

[1]**H NMR** (500 MHz, DMSO-*d6*) δ [ppm]: 7.96 – 7.91 (m, 2H), 7.79 (d, J = 2.6 Hz, 1H), 7.75 (d, J = 8.9 Hz, 1H), 7.49 (dd, J = 9.0, 2.6 Hz, 1H), 7.44 (s, 4H), 7.17 (d, J = 9.0 Hz, 2H), 5.45 (s, 2H), 4.46 (s, 1H), 3.97 – 3.91 (m, 2H), 3.47 – 3.41 (m, 2H), 3.32 (s, 3H),1.74 – 1.66 (m, 2H). [13]**C NMR** (126 MHz, DMSO-*d6*) δ [ppm]: 162.67, 154.65, 153.95, 151.89, 150.56, 146.41, 145.49, 145.48 135.19, 134.31, 132.68, 129.66, 128.78, 125.15, 121.41, 119.40, 118.62, 114.82, 102.75, 58.82, 55.78, 45.99, 38.38, 31.01, 29.63. **HRMS** (ESI, m/z): $[M+H]^+$ calcd for $C_{29}H_{27}Cl_2N_6O_5$: 609.1414; found: 609.1408.

**AzHC3**

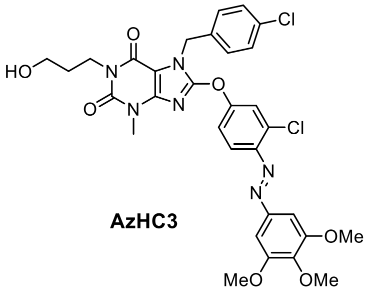

**AzHC3**

A vial was charged with **S2** (43 mg, 0.10 mmol, 1.0 eq), **S5** (32 mg, 0.10 mmol, 1.0 eq), $K_2CO_3$ (28 mg, 0.20 mmol, 2.0 eq) and DMF (1 ml), sealed and the reaction was stirred at 80°C. After 5 h the reaction was cooled to room temperature, $H_2O$ (2 ml) was added, and after 30 min the precipitate was collected by filtration. The crude product was recrystallized from DMSO/$H_2O$ (volumetric ratio 9:1) and the filter cake washed with cold MeOH to afford **AzHC3** (15 mg, 0.022 mmol 22 %) as orange solid.

[1]**H NMR** (400 MHz, DMSO-*d6*) δ [ppm]: 7.82 (d, *J* = 2.6 Hz, 1H), 7.76 (d, *J* = 9.0 Hz, 1H), 7.50 (dd, *J* = 9.0, 2.6 Hz, 1H), 7.45 – 7.42 (m, 4H), 7.30 (s, 2H), 5.45 (s, 2H), 4.46 (t, *J* = 5.2 Hz, 1H), 3.97 – 3.90 (m, 2H), 3.89 (s, 6H), 3.78 (s, 3H), 3.49 – 3.41 (m, 2H), 3.32 (s, 3H), 1.75 – 1.65 (m, 2H). [13]**C NMR** (101 MHz, DMSO-*d6*) δ [ppm]: 155.05, 153.96, 153.39, 151.77, 150.56, 147.92, 145.47, 145.31, 141.09, 135.17, 134.67, 132.68, 129.66, 128.78, 121.42, 119.38, 118.74, 102.78, 100.78, 60.33, 58.81, 56.04, 46.19, 38.43, 31.01, 29.64. **HRMS** (ESI, m/z): $[M+H]^+$ calcd for $C_{31}H_{31}Cl_2N_6O_7$: 669.1626; found: 669.1616.

**AzHC4**

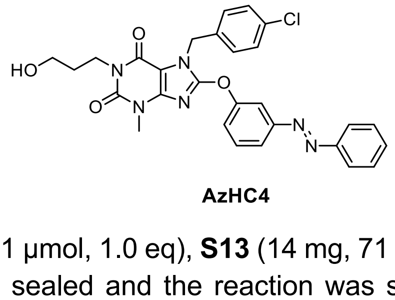

**AzHC4**

A vial was charged with **S2** (30 mg, 71 µmol, 1.0 eq), **S13** (14 mg, 71 µmol, 1.0 eq), $K_2CO_3$ (20 mg, 141 µmol, 2.0 eq) and DMF (1 ml), sealed and the reaction was stirred at 80°C. After 5 h the reaction was cooled to room temperature, $H_2O$ (2 ml) was added, and after 30 min the precipitate was collected by filtration. The crude product was recrystallized from DMSO/$H_2O$ (volumetric ratio 9:1) and the filter cake washed with cold MeOH to afford **AzHC4** (9 mg, 17 µmol, 23 %) as yellow solid.

[1]**H NMR** (400 MHz, DMSO-*d6*) δ [ppm]: 7.94 – 7.89 (m, 2H), 7.88 – 7.83 (m, 1H), 7.79 – 7.77 (m, 1H), 7.70 (t, *J* = 8.0 Hz, 1H), 7.65 – 7.53 (m, 4H), 7.47 – 7.42 (m, 4H), 5.47 (s, 2H), 4.45 (t, *J* = 5.2 Hz, 1H), 3.98 – 3.86 (m, 2H), 3.49 – 3.40 (m, 2H), 3.29 (s, 3H), 1.75 – 1.65 (m, 2H). [13]**C NMR** (101 MHz, DMSO-*d6*) δ [ppm]: 153.94, 153.90, 153.01, 152.66, 151.70, 150.58, 145.57, 135.34, 132.64, 132.06, 130.90,

129.61, 129.56, 128.80, 122.75, 122.63, 121.39, 112.31, 102.60, 58.82, 45.92, 38.34, 31.02, 29.61. **HRMS** (ESI, m/z): $[M+H]^+$ calcd for $C_{28}H_{26}ClN_6O_4$: 545.1699; found: 545.1693.

#### AzHC5

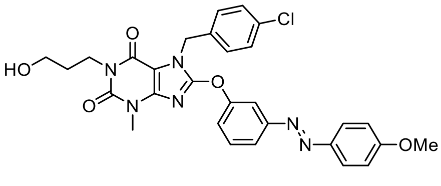

**AzHC5**

A vial was charged with **S2** (73 mg, 0.17 mmol, 1.0 eq), **S14** (39 mg, 0.17 mmol, 1.0 eq), $K_2CO_3$ (39 mg, 0.17 mmol, 2.0 eq) and DMF (1.7 ml), sealed and the reaction was stirred at 80°C. After 5 h the reaction was cooled to room temperature, $H_2O$ (3.4 ml) was added, and after 30 min the precipitate was collected by filtration. The crude product was recrystallized from DMSO/$H_2O$ (volumetric ratio 9:1) and the filter cake washed with cold MeOH to afford **AzHC5** (22 mg, 0.038 mmol, 23 %) as brown solid.

**$^1$H NMR** (400 MHz, DMSO-*d6*) δ [ppm]: 7.91 (d, *J* = 9.0 Hz, 2H), 7.83 – 7.78 (m, 1H), 7.73 – 7.70 (m, 1H), 7.66 (t, *J* = 8.1 Hz, 1H), 7.53 – 7.47 (m, 1H), 7.46 – 7.42 (m, 4H), 7.15 (d, *J* = 9.0 Hz, 2H), 5.46 (s, 2H), 4.45 (t, *J* = 5.2 Hz, 1H), 3.98 – 3.90 (m, 2H), 3.88 (s, 3H), 3.49 – 3.40 (m, 2H), 3.28 (s, 3H), 1.74 – 1.64 (m, 2H). **$^{13}$C NMR** (101 MHz, DMSO-*d6*) δ 162.46, 153.93, 153.89, 153.18, 152.72, 150.57, 145.95, 145.59, 135.35, 132.64, 130.77, 129.61, 128.80, 124.89, 121.87, 121.07, 114.73, 112.04, 102.57, 58.82, 55.72, 45.91, 38.34, 31.02, 29.60. **HRMS** (ESI, m/z): $[M+H]^+$ calcd for $C_{29}H_{28}ClN_6O$: 575.1804; found: 575.1799.

#### AzHC6

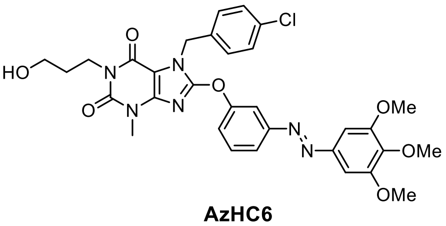

**AzHC6**

A vial was charged with **S2** (81 mg, 0.19 mmol, 1.0 eq), **S15** (55 mg, 0.19 mmol, 1.0 eq), $K_2CO_3$ (53 mg, 0.38 mmol, 2.0 eq) and DMF (2 ml), sealed and the reaction was stirred at 80°C. After 5 h the reaction was cooled to room temperature, $H_2O$ (4 ml) was added, and after 30 min the precipitate was collected by filtration. The crude product was recrystallized from DMSO/$H_2O$ (volumetric ratio 9:1) and the filter cake washed with cold MeOH to afford **AzHC5** (17 mg, 0.027 mmol, 14 %) as orange solid.

**$^1$H NMR** (400 MHz, DMSO-*d6*) δ [ppm]: 7.89 – 7.80 (m, 1H), 7.75 (t, *J* = 2.2 Hz, 1H), 7.69 (t, *J* = 8.1 Hz, 1H), 7.59 – 7.52 (m, 1H), 7.48 – 7.42 (m, 4H), 7.28 (s, 2H), 5.47 (s, 2H), 4.45 (t, *J* = 5.2 Hz, 1H), 4.02 – 3.90 (m, 2H), 3.89 (s, 6H), 3.77 (s, 3H), 3.49 – 3.40 (m, 2H), 3.28 (s, 3H), 1.75 – 1.63 (m, 2H). **$^{13}$C NMR** (101 MHz, DMSO-*d6*) δ [ppm]: 153.93, 153.92, 153.37, 152.94, 152.71, 150.57, 147.49, 145.58, 140.83, 135.34, 132.64, 130.86, 129.61, 128.80, 122.34, 121.25, 112.32, 102.60, 100.57, 60.29, 58.82, 56.05, 45.91, 38.34, 31.02, 29.61. **HRMS** (ESI, m/z): $[M+H]^+$ calcd for $C_{31}H_{32}ClN_6O_7$: 635.2016; found: 635.2007.

## S1

$^{1}$**H NMR** (400 MHz, DMSO-*d6*)

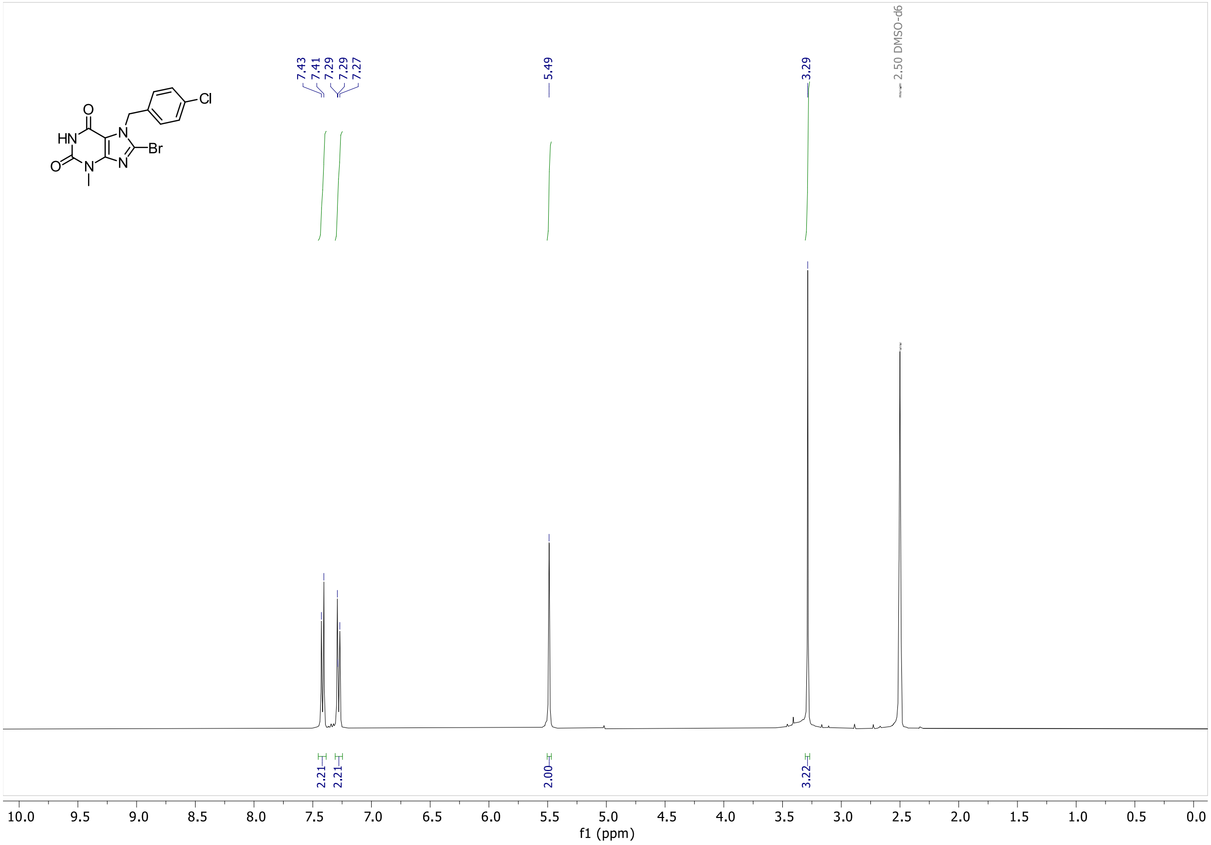

$^{13}$**C NMR** (101 MHz, DMSO-*d6*)

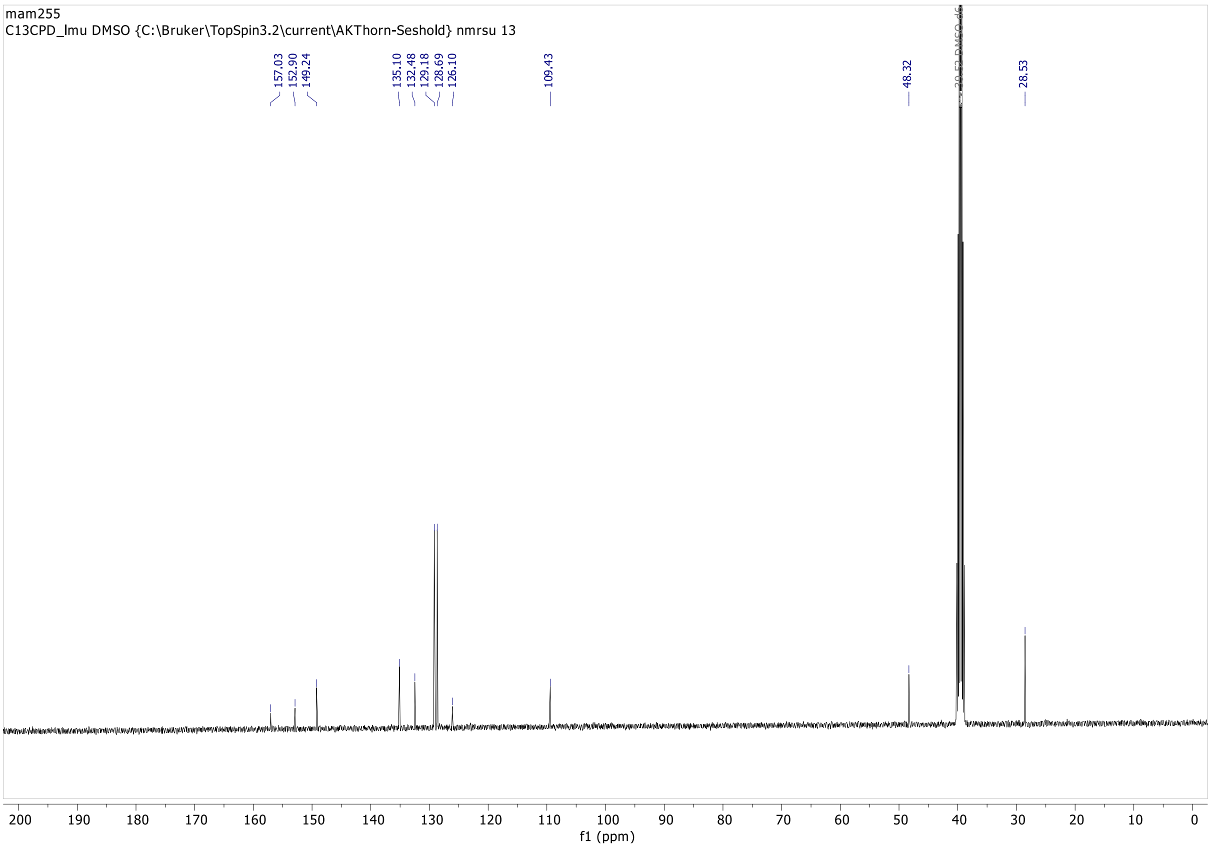

**S2**

**$^{1}$H NMR** (400 MHz, DMSO-*d6*)

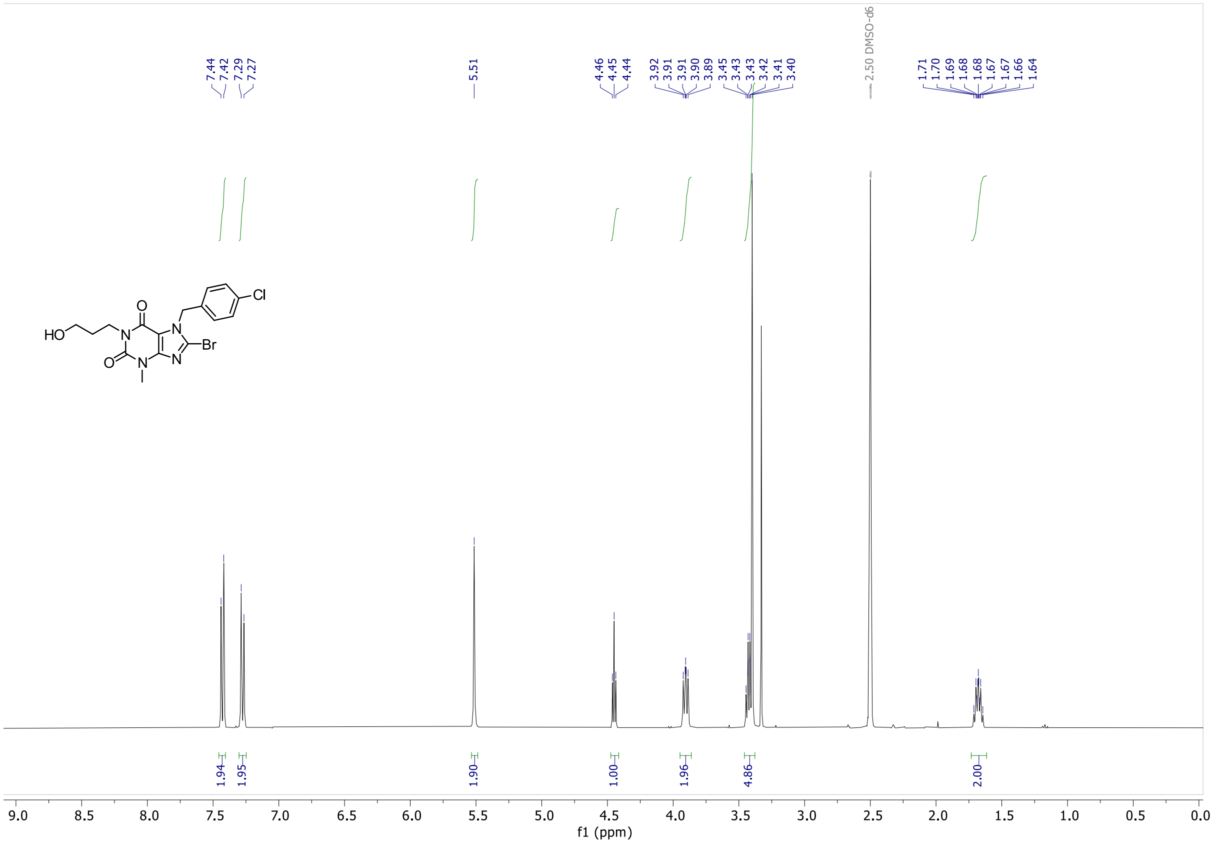

**$^{13}$C NMR** (101 MHz, DMSO-*d6*)

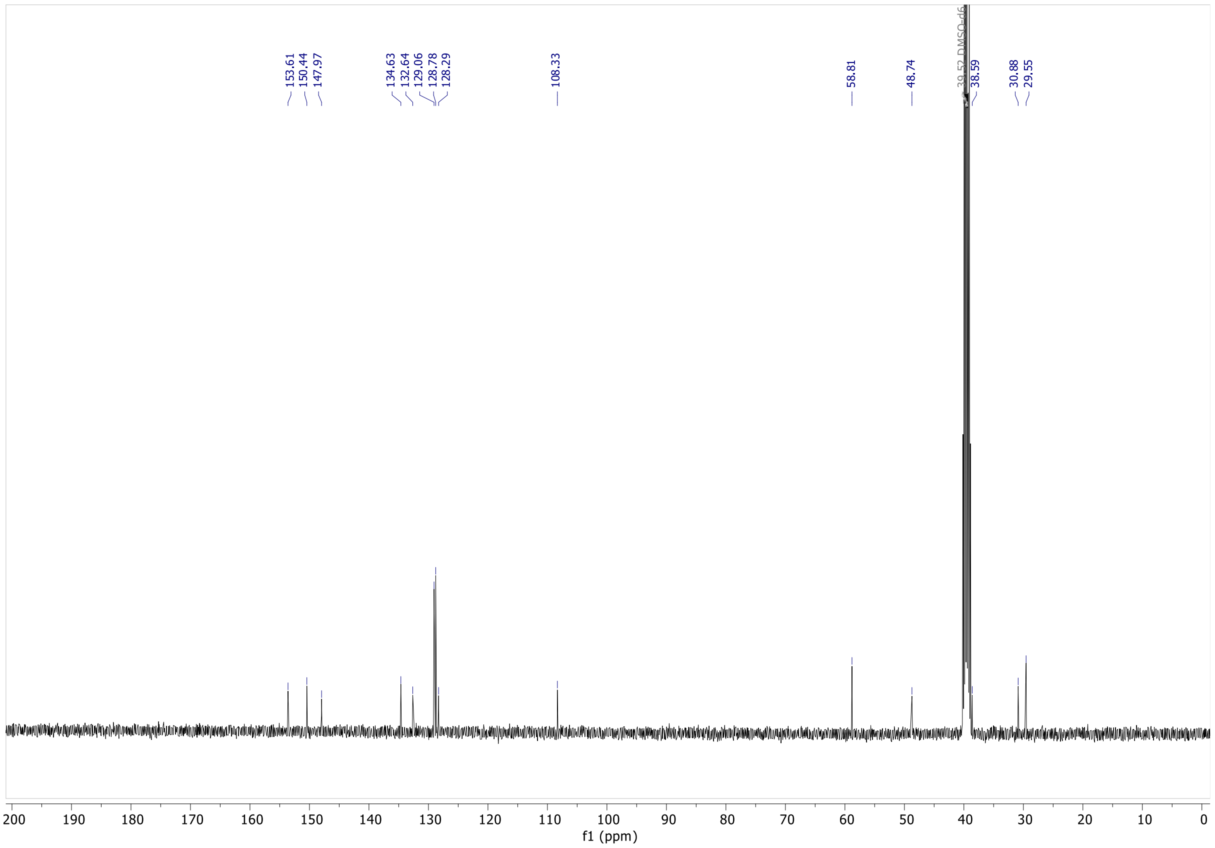

## S3

**¹H NMR** (400 MHz, $CDCl_3$)

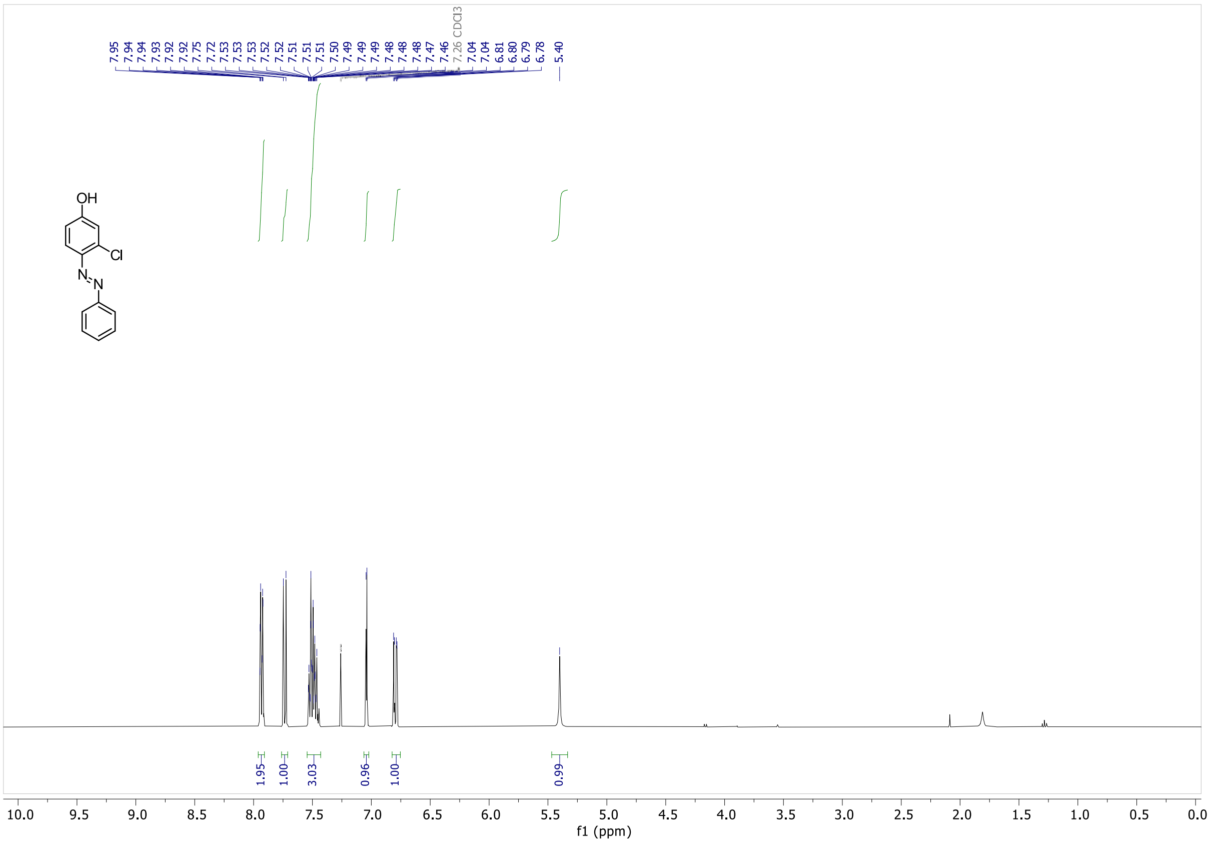

**¹³C NMR** (101 MHz, $CDCl_3$)

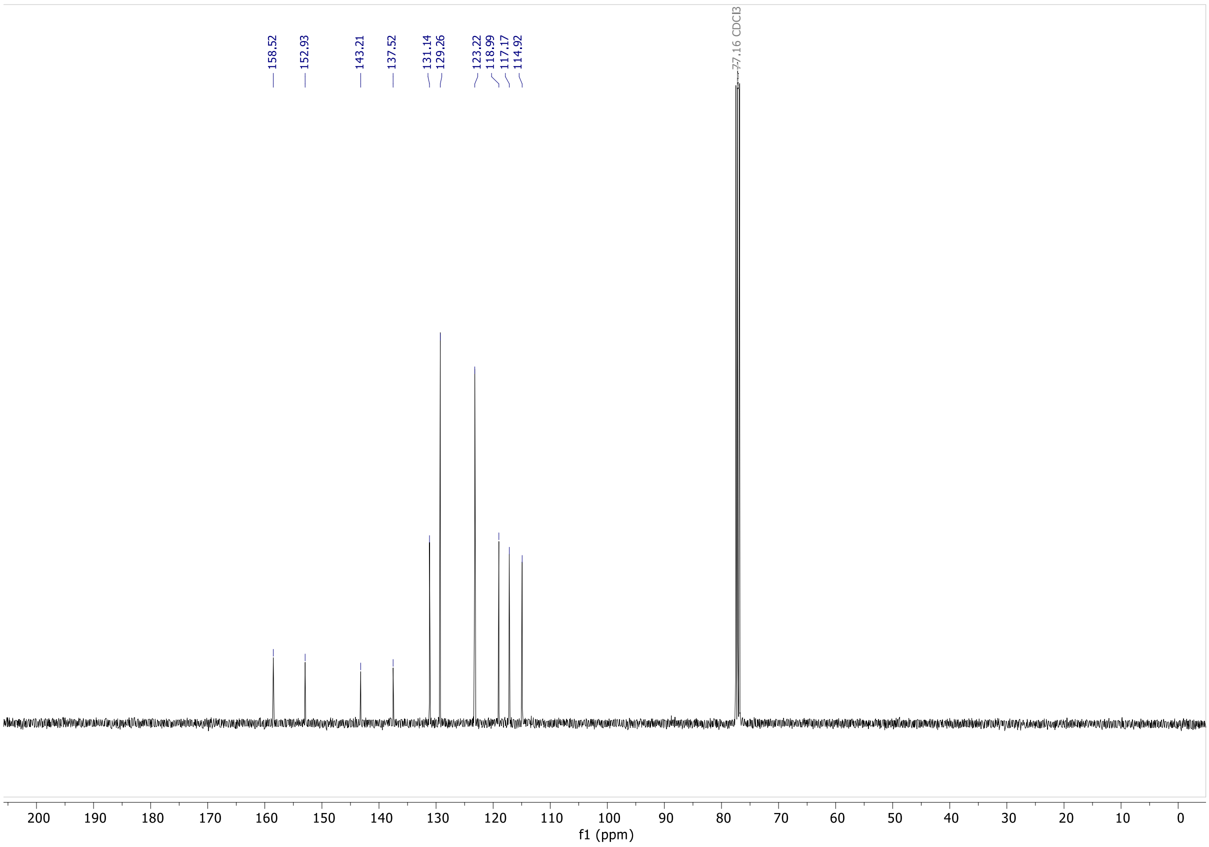

**S4**

**¹H NMR** (400 MHz, $CDCl_3$)

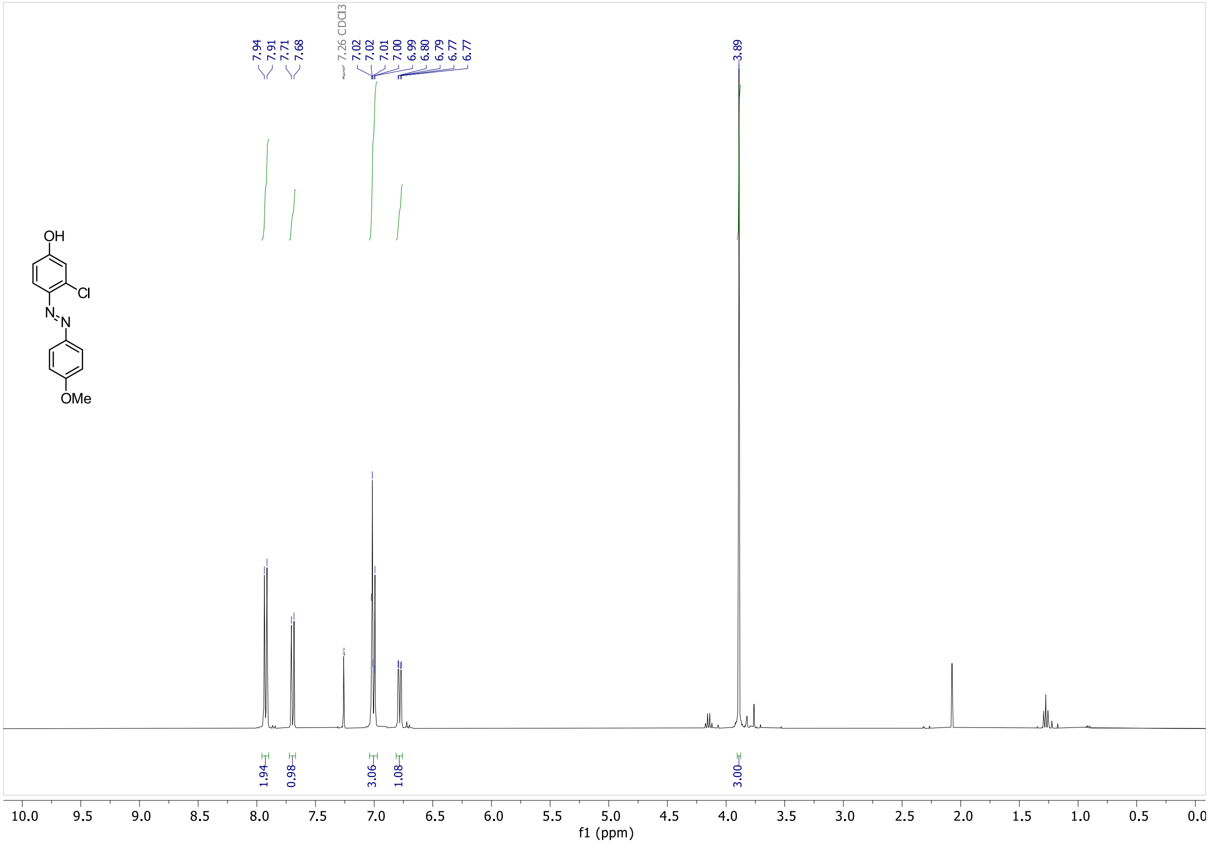

**¹³C NMR** (101 MHz, $CDCl_3$)

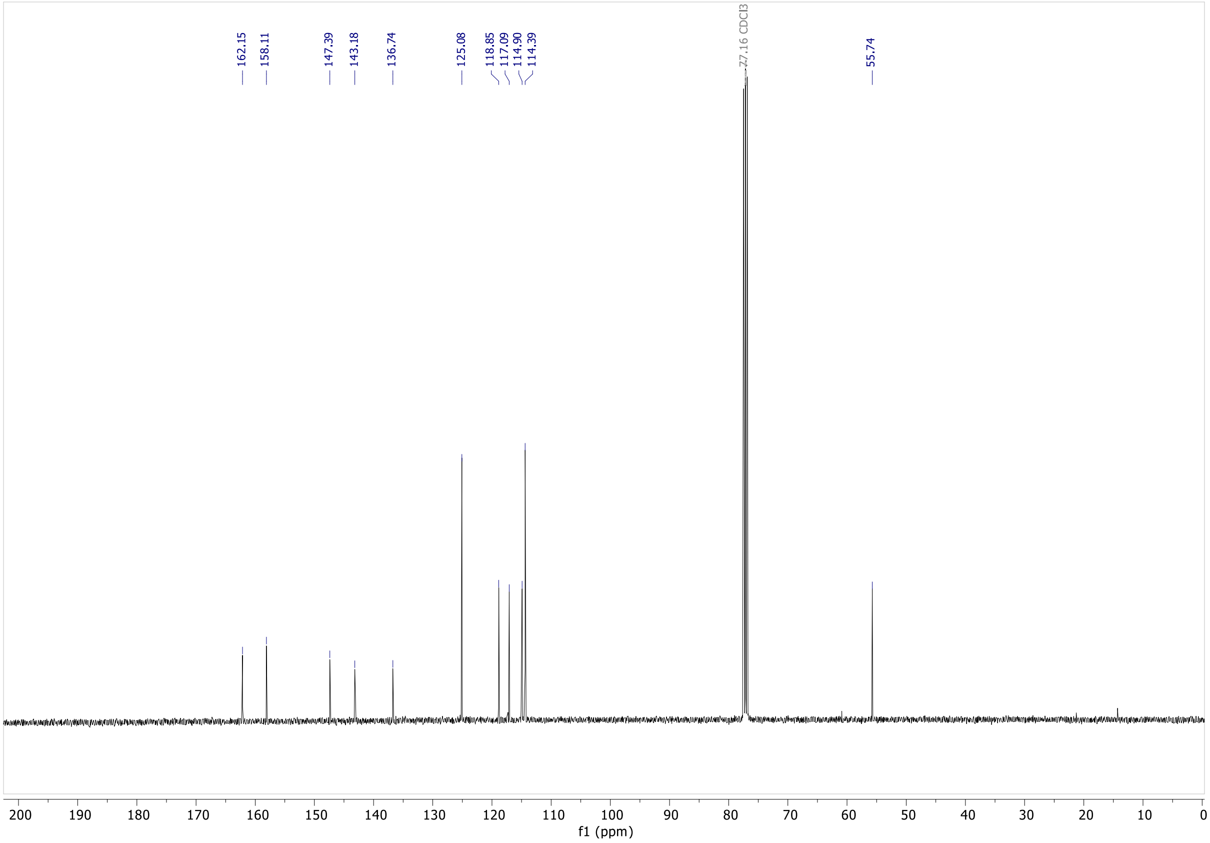

**S5**

$^{1}$**H NMR** (400 MHz, $CDCl_3$)

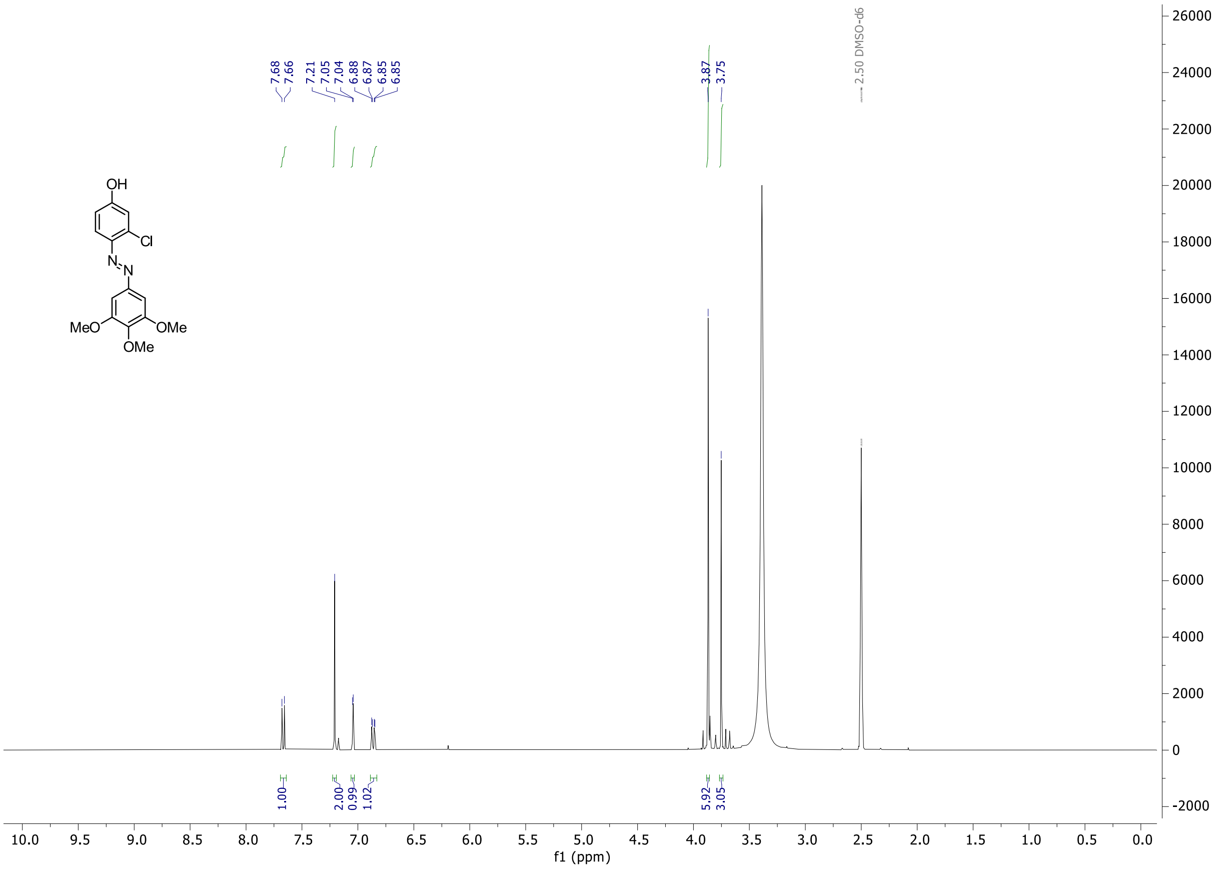

## S6

**¹H NMR** (500 MHz, $CDCl_3$)

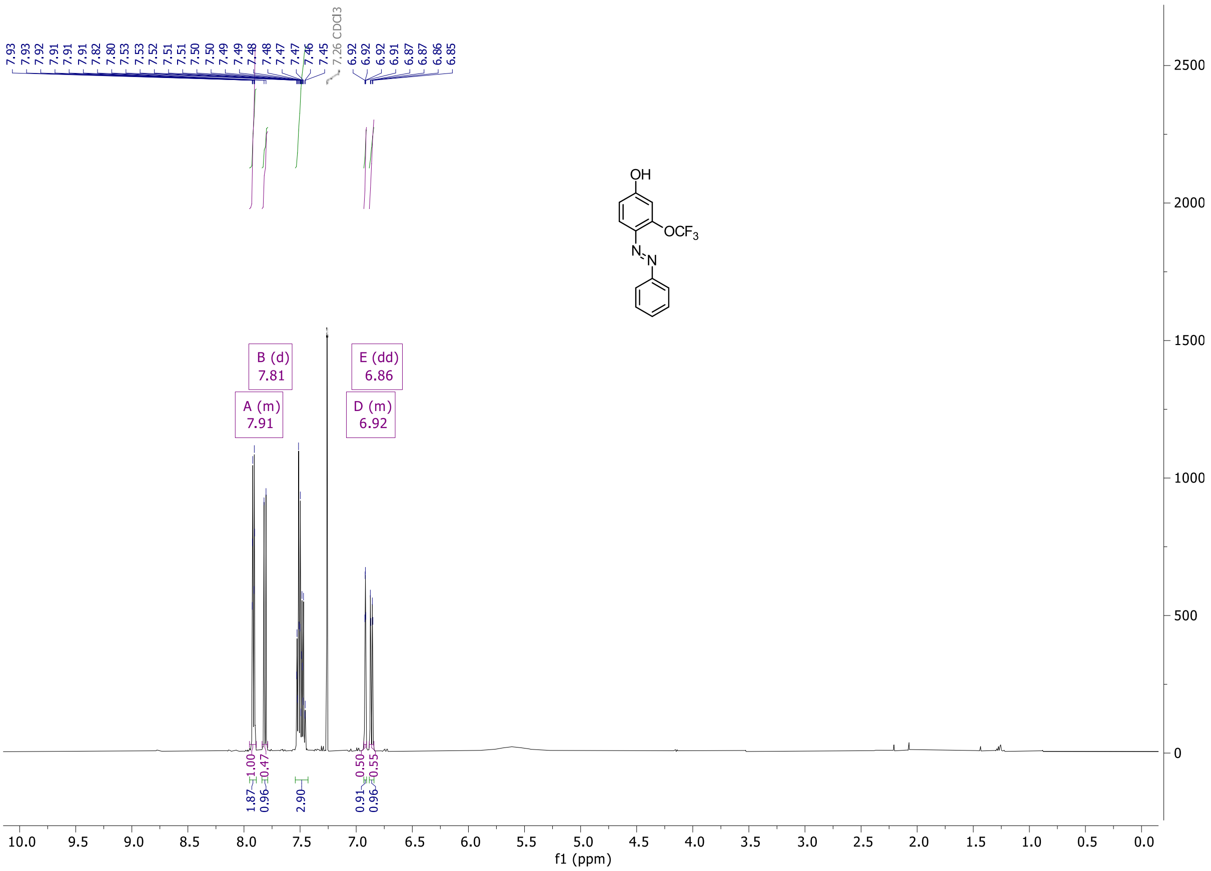

**¹³C NMR** (126 MHz, $CDCl_3$)

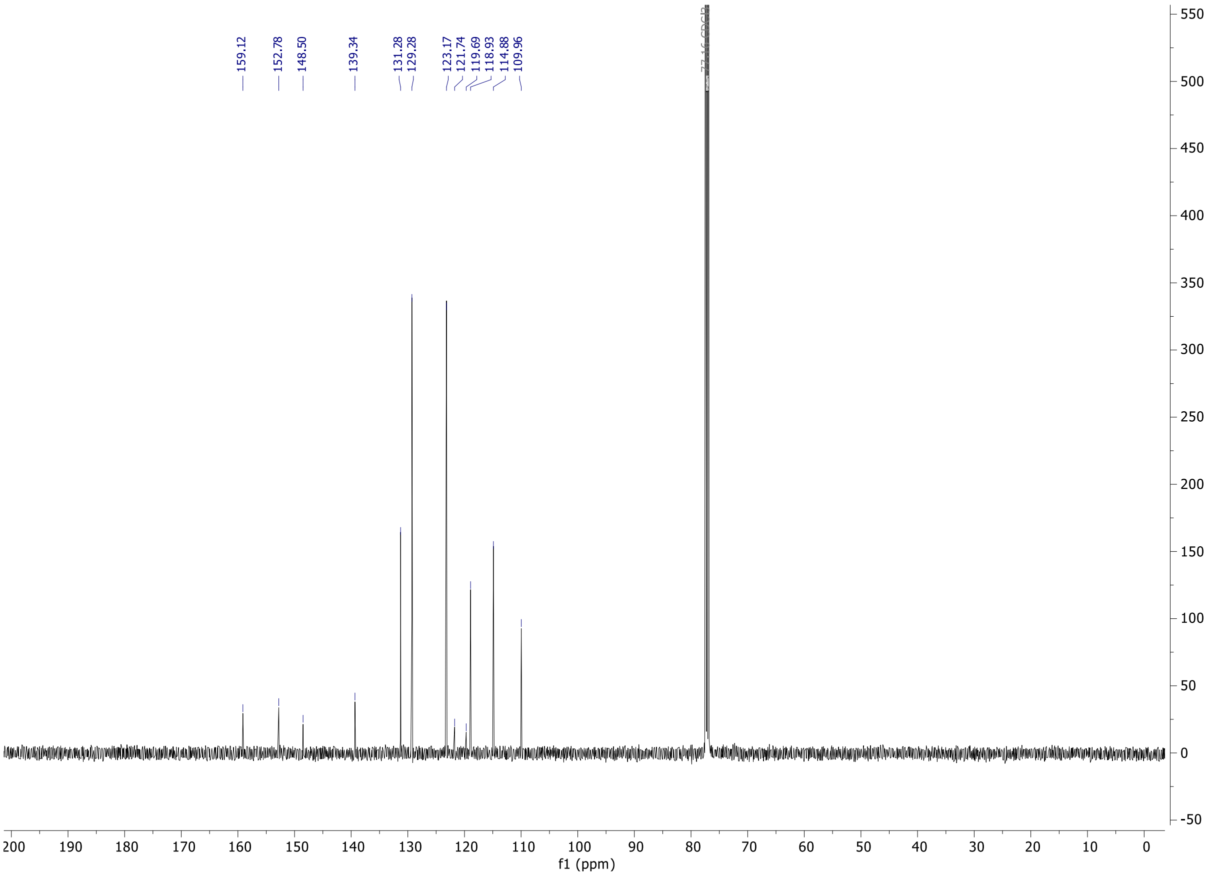

**S7**

**¹H NMR** (400 MHz, $CDCl_3$)

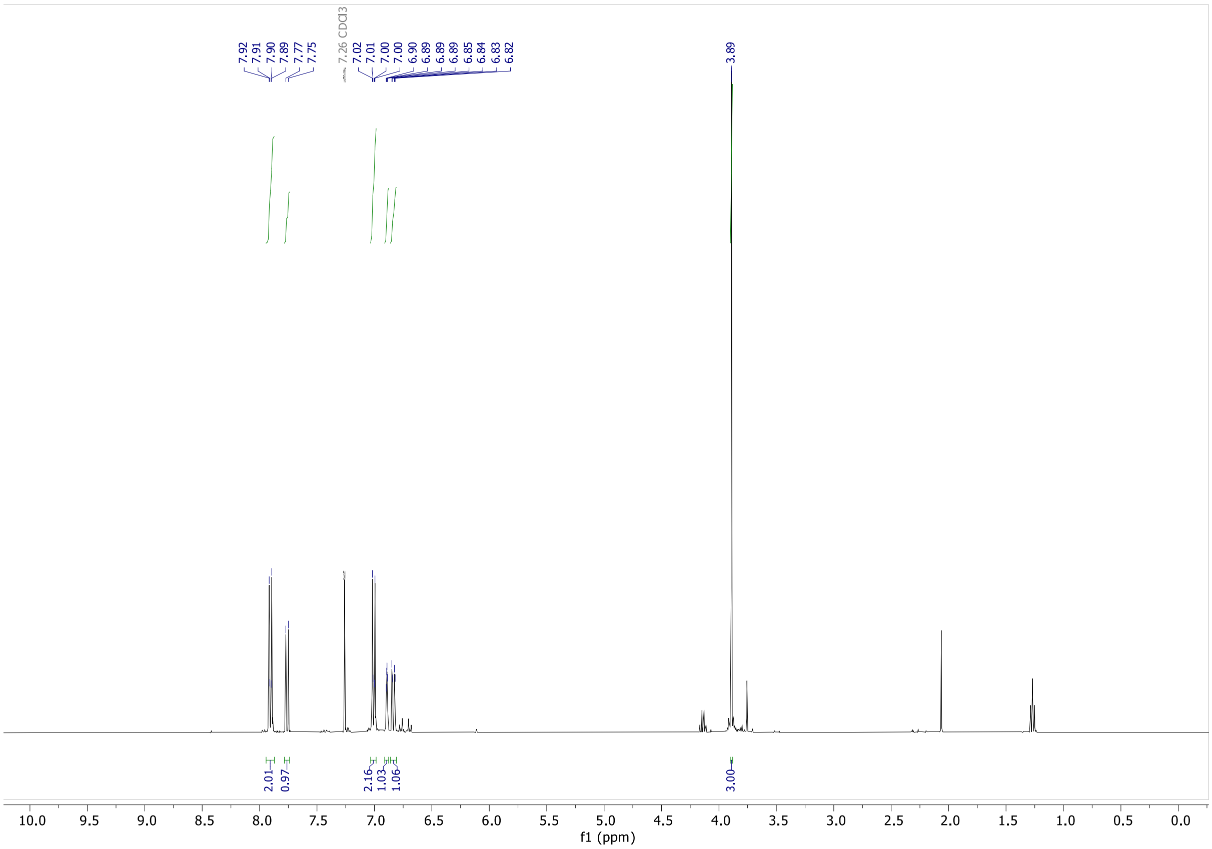

**¹³C NMR** (101 MHz, $CDCl_3$)

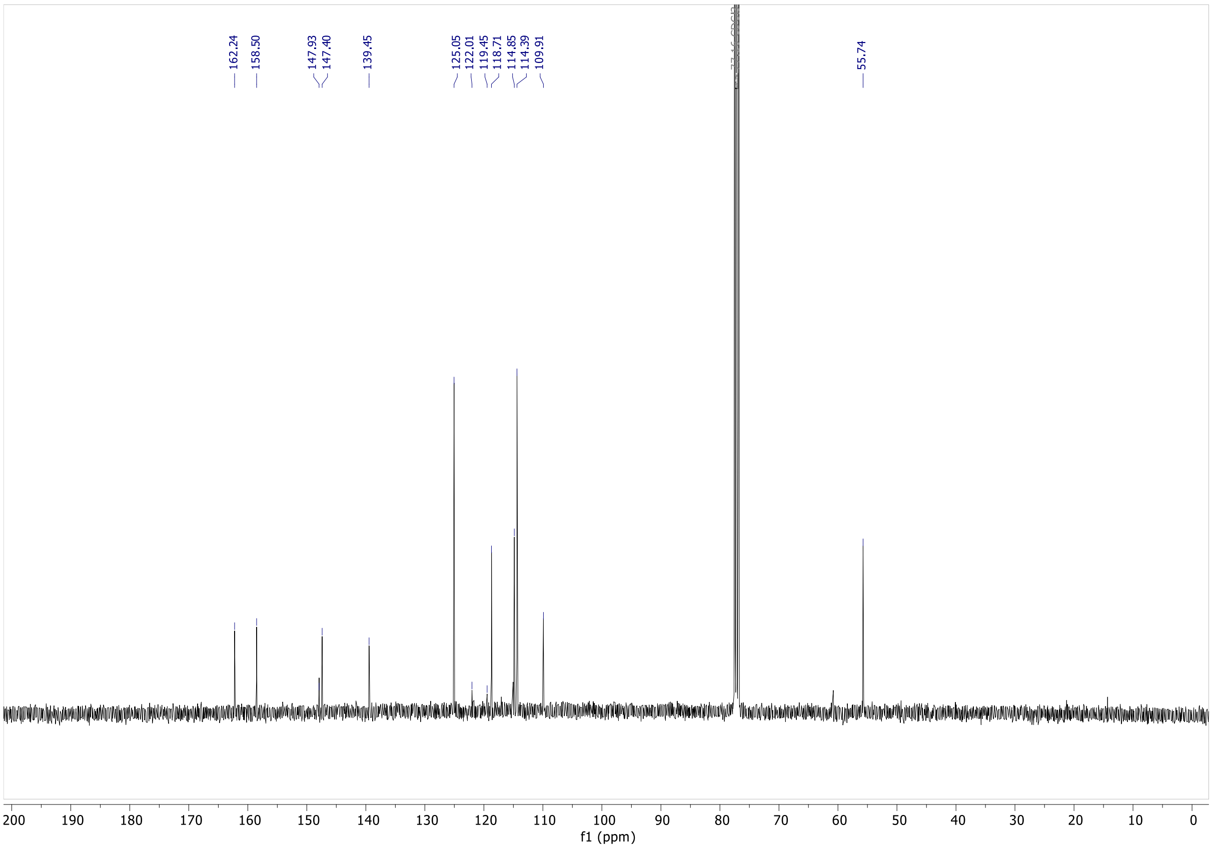

**S8**

**[1]H NMR** (400 MHz, $CDCl_3$)

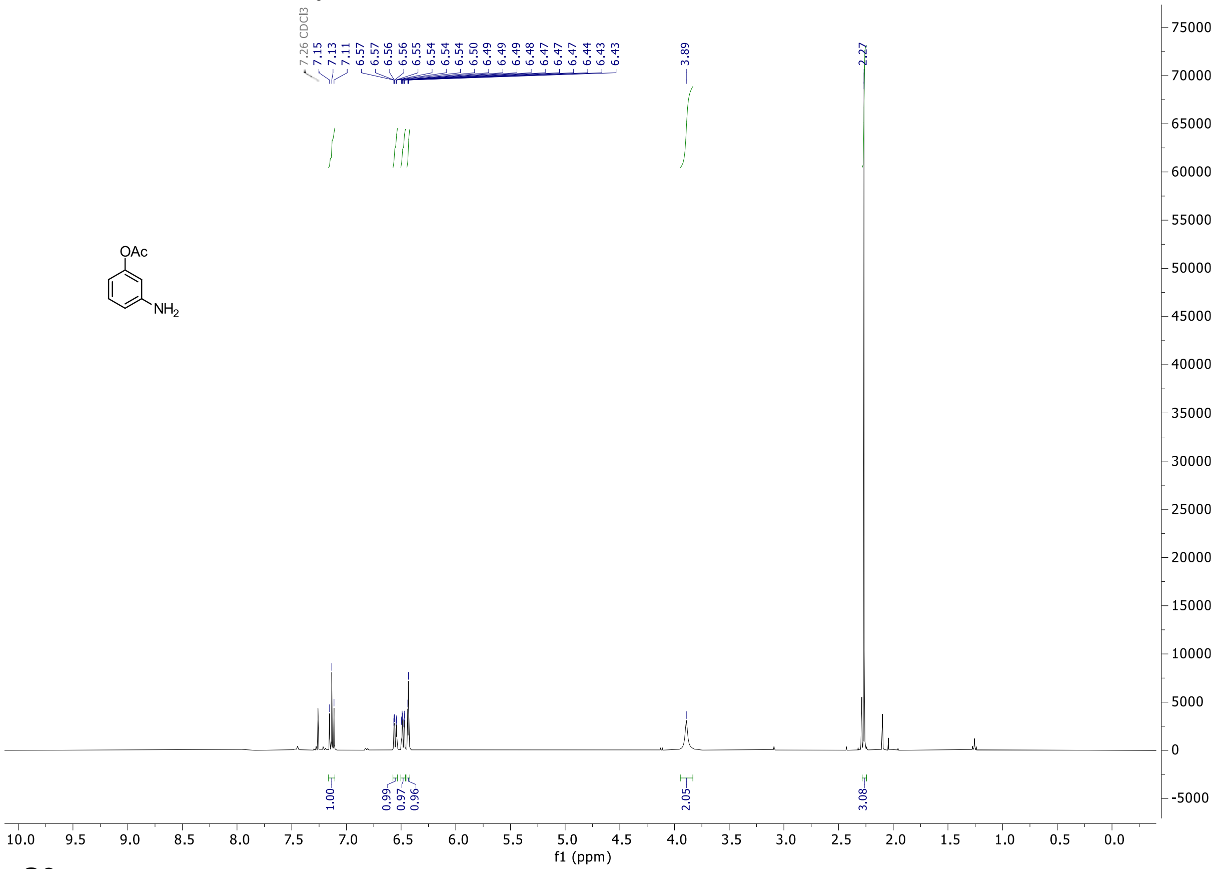

**S9**

**[1]H NMR** (400 MHz, $CDCl_3$)

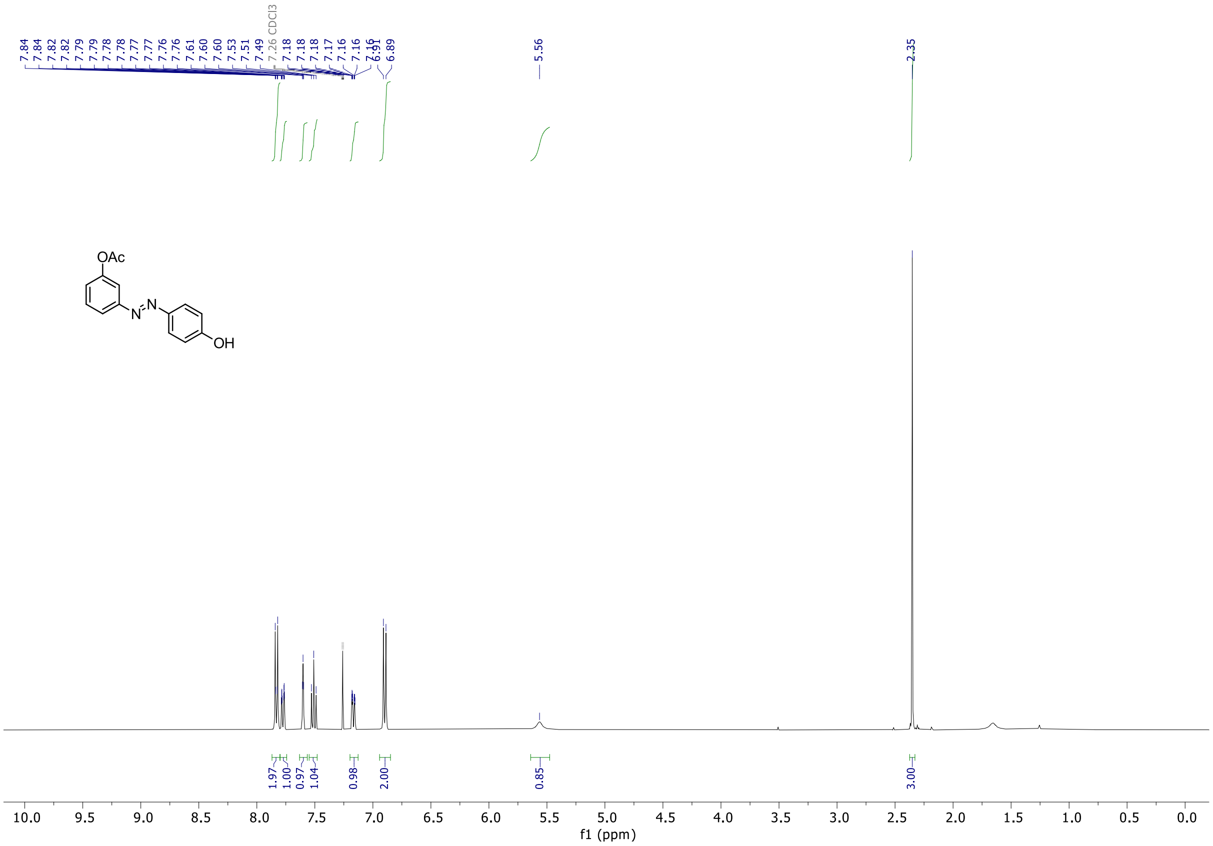

## S10

**$^1$H NMR** (400 MHz, $CDCl_3$)

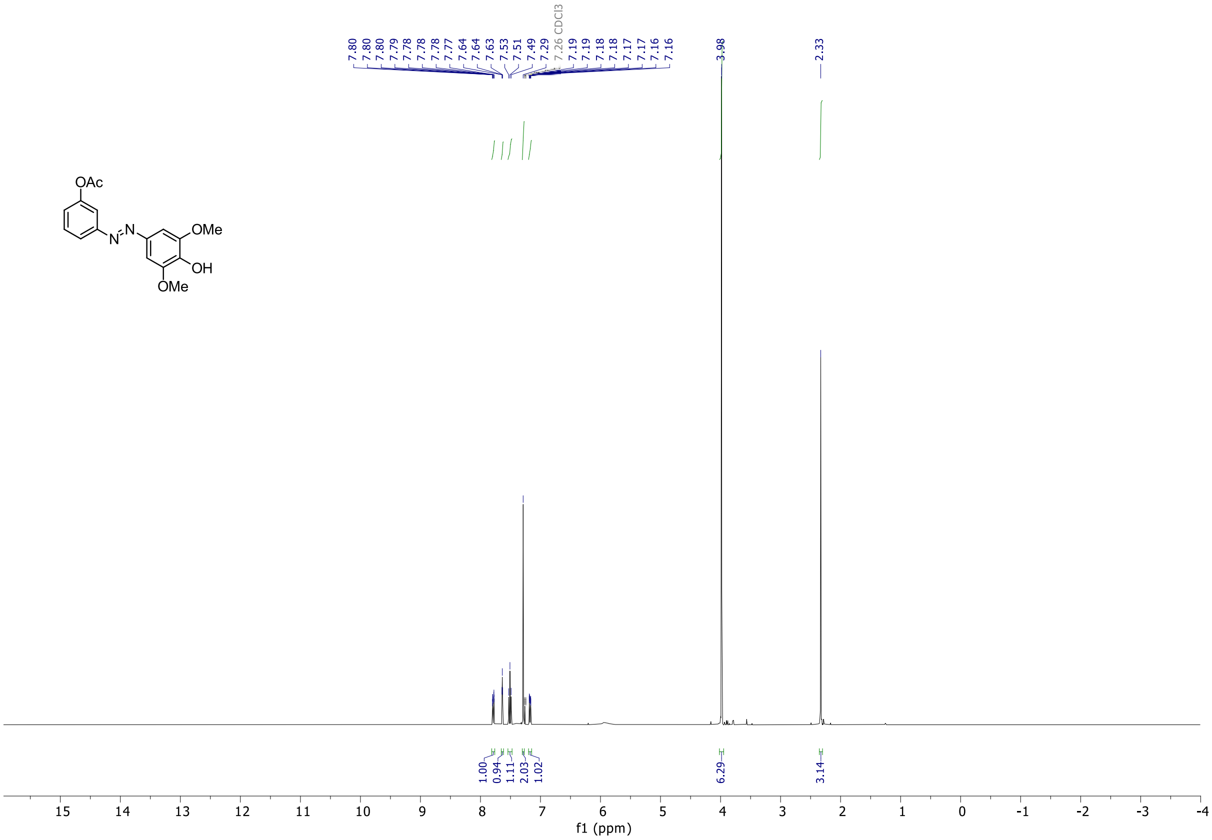

## S13

**$^1$H NMR** (400 MHz, $CDCl_3$)

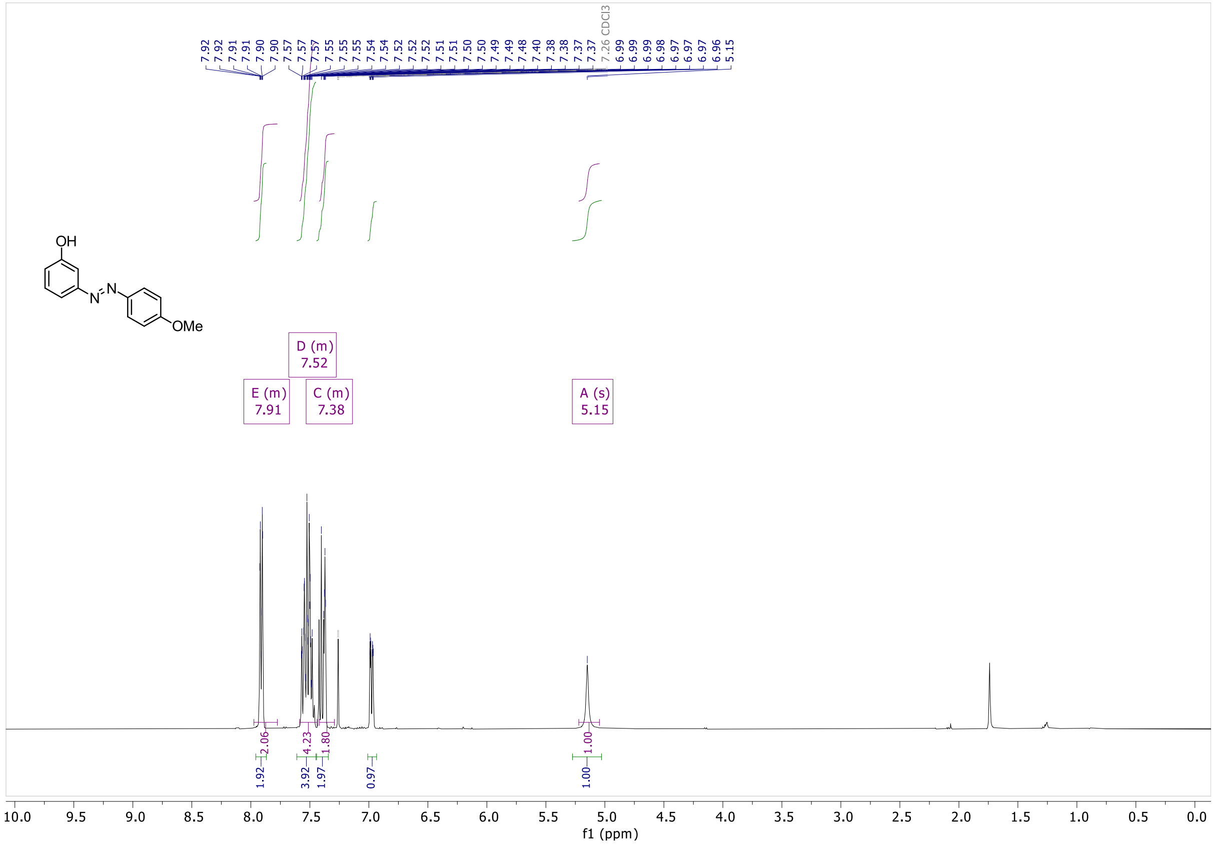

#### AzPico

**¹H NMR** (500 MHz, DMSO-*d6*)

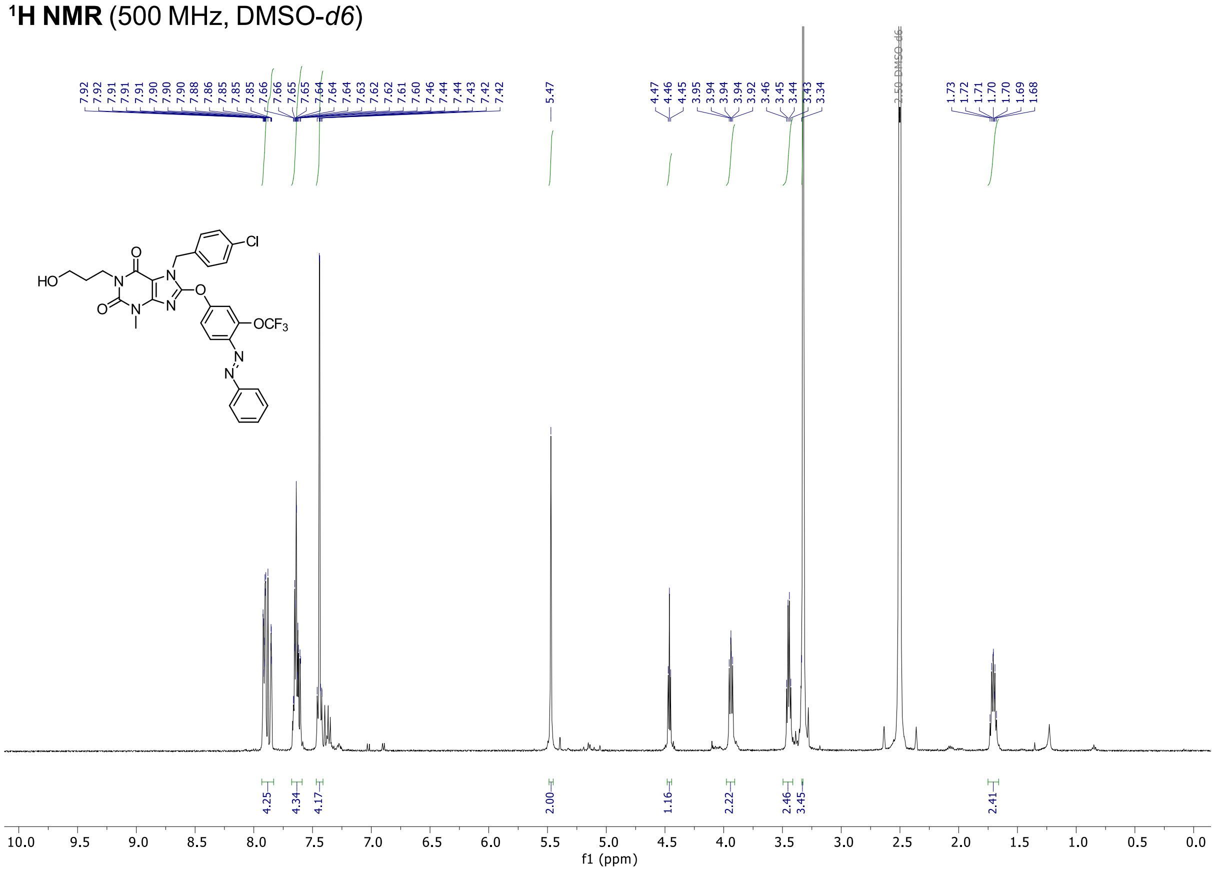

**¹³C NMR** (126 MHz, DMSO-*d6*)

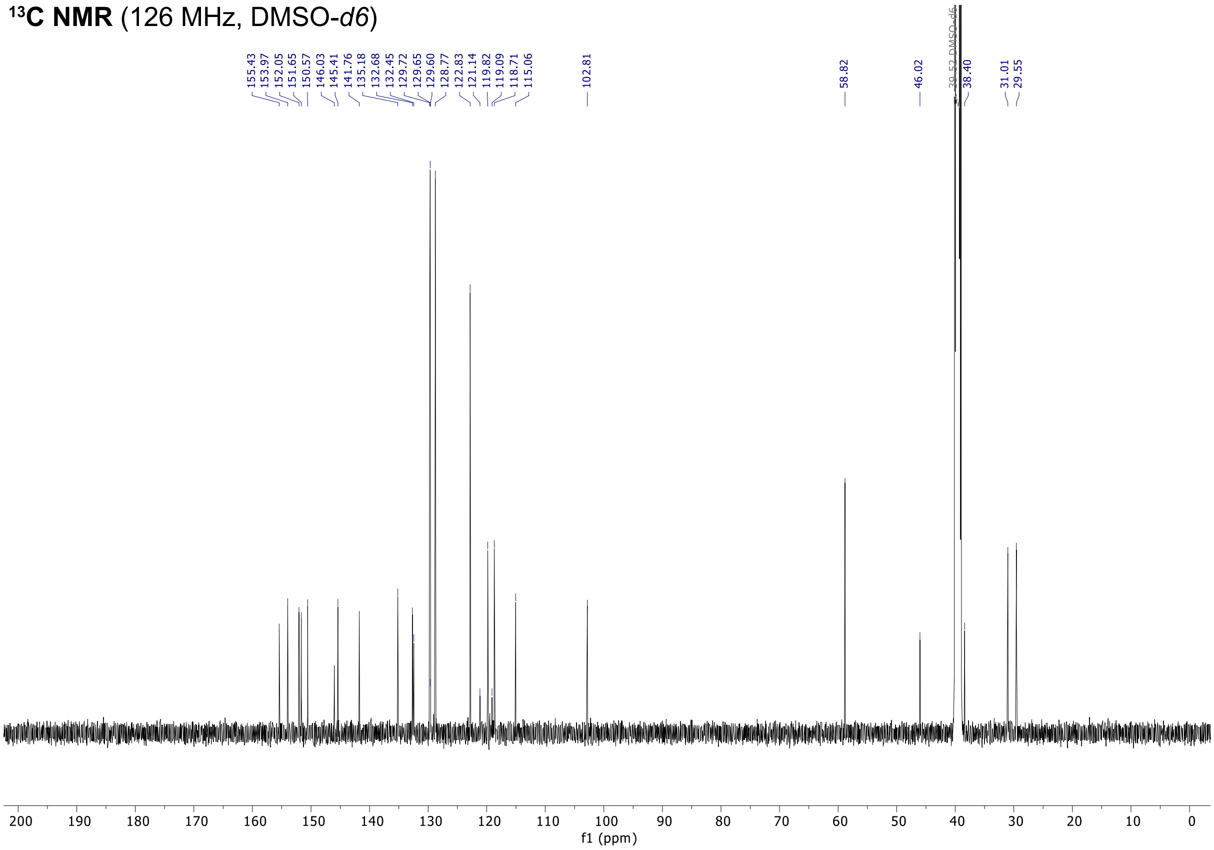

#### AzHC

**$^{1}$H NMR** (400 MHz, DMSO-*d6*)

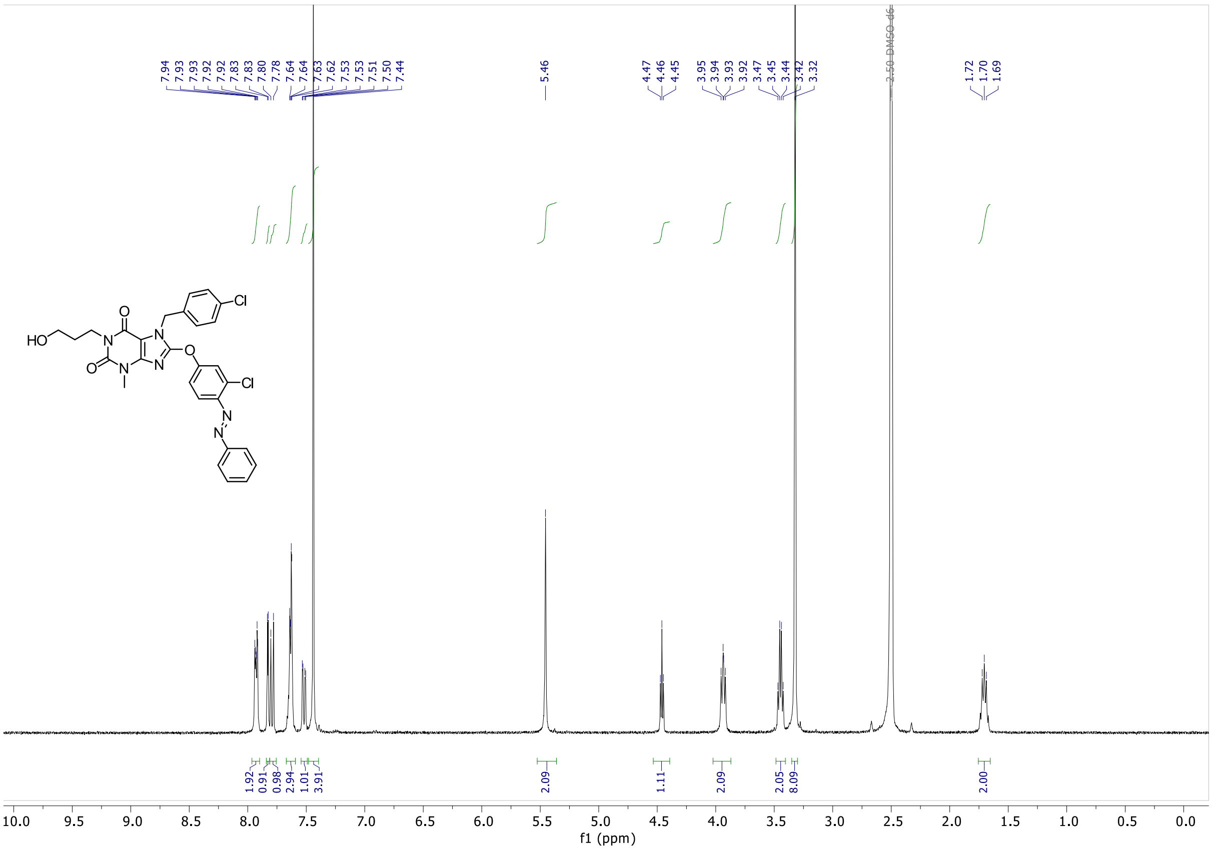

**$^{13}$C NMR** (126 MHz, DMSO-*d6*)

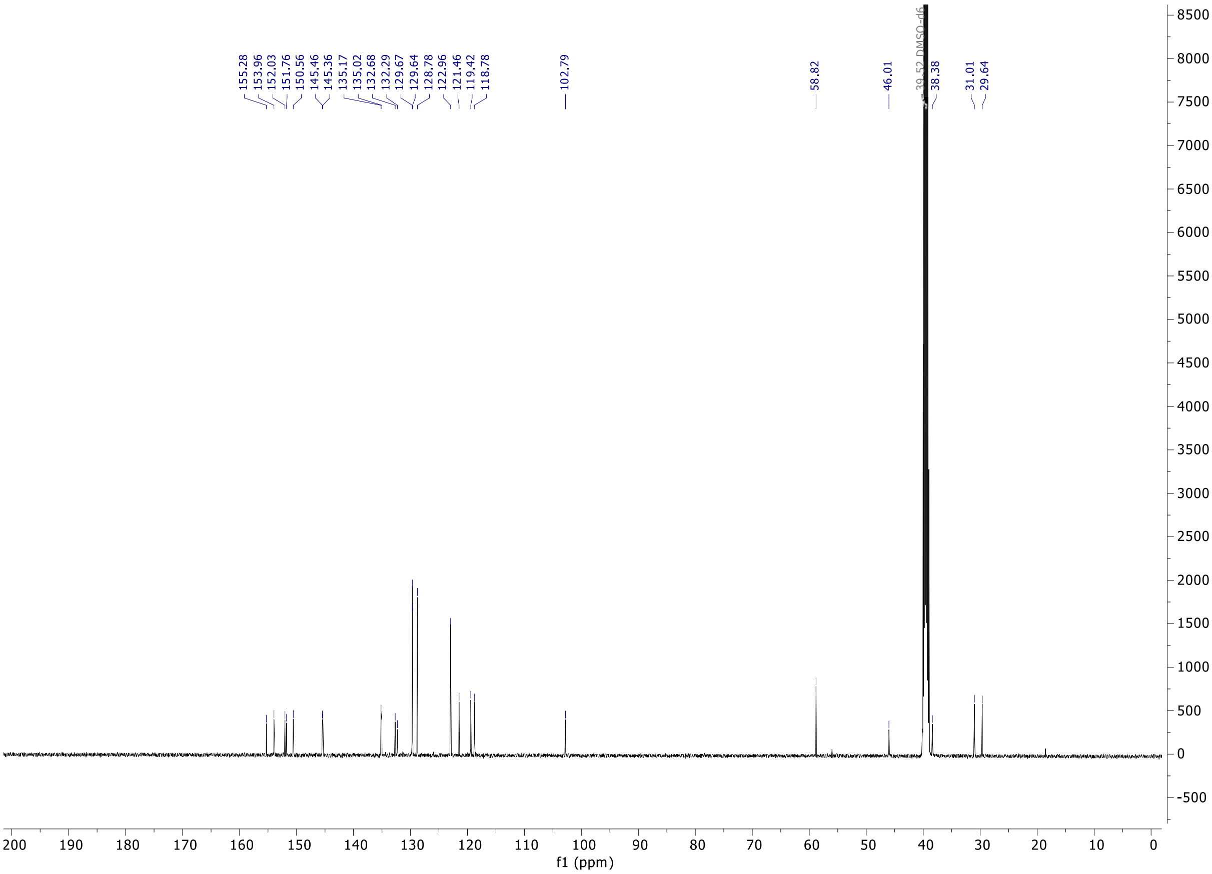

#### AzPico2

**$^1$H NMR** (500 MHz, DMSO-*d6*)

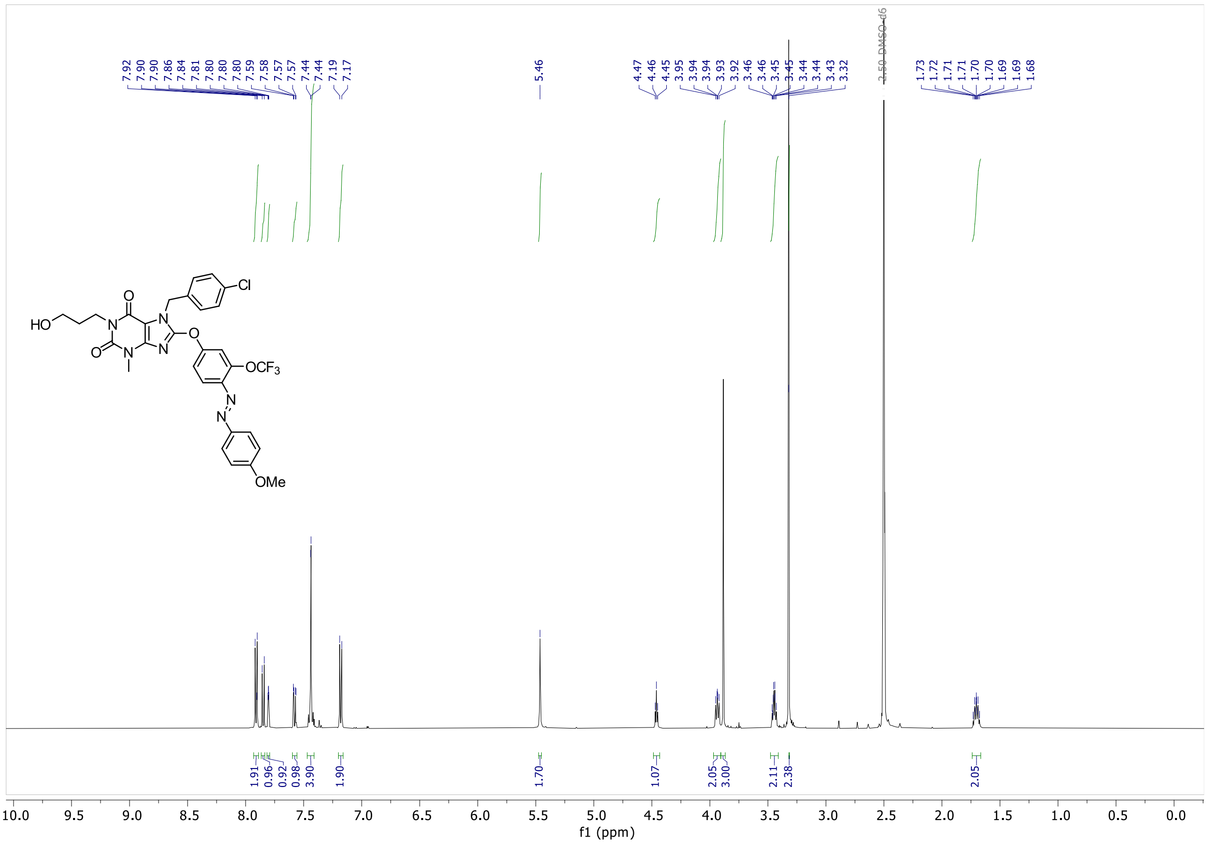

**$^{13}$C NMR** (126 MHz, DMSO-*d6*)

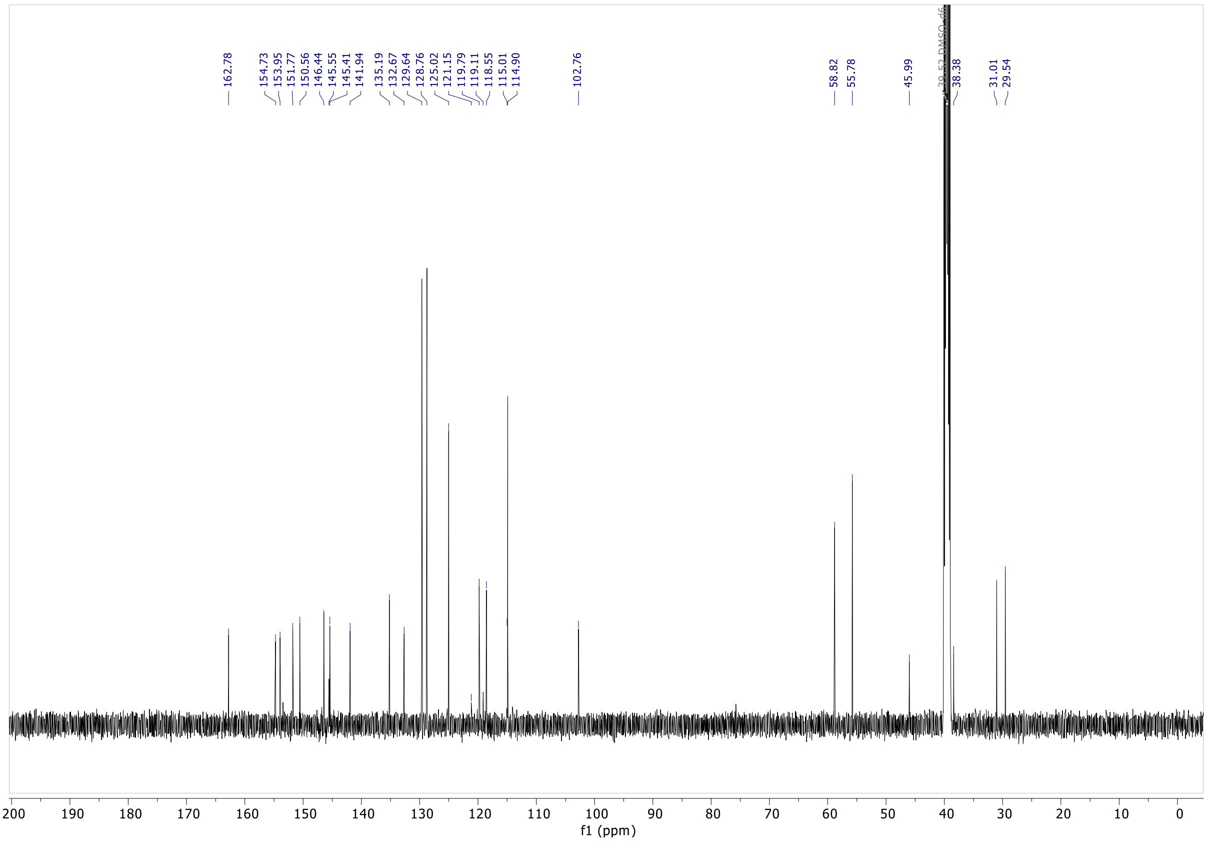

#### AzHC2

**¹H NMR** (500 MHz, DMSO-*d6*)

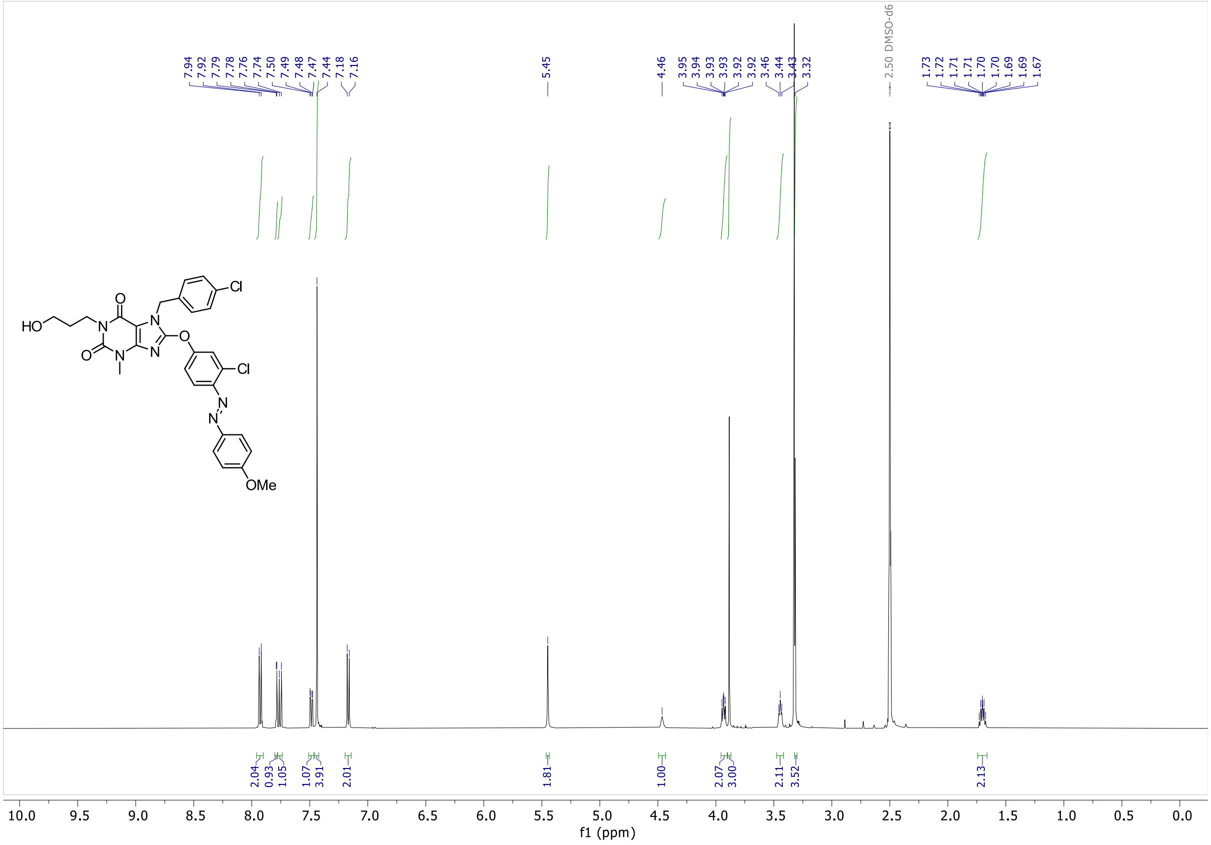

**¹³C NMR** (126 MHz, DMSO-*d6*)

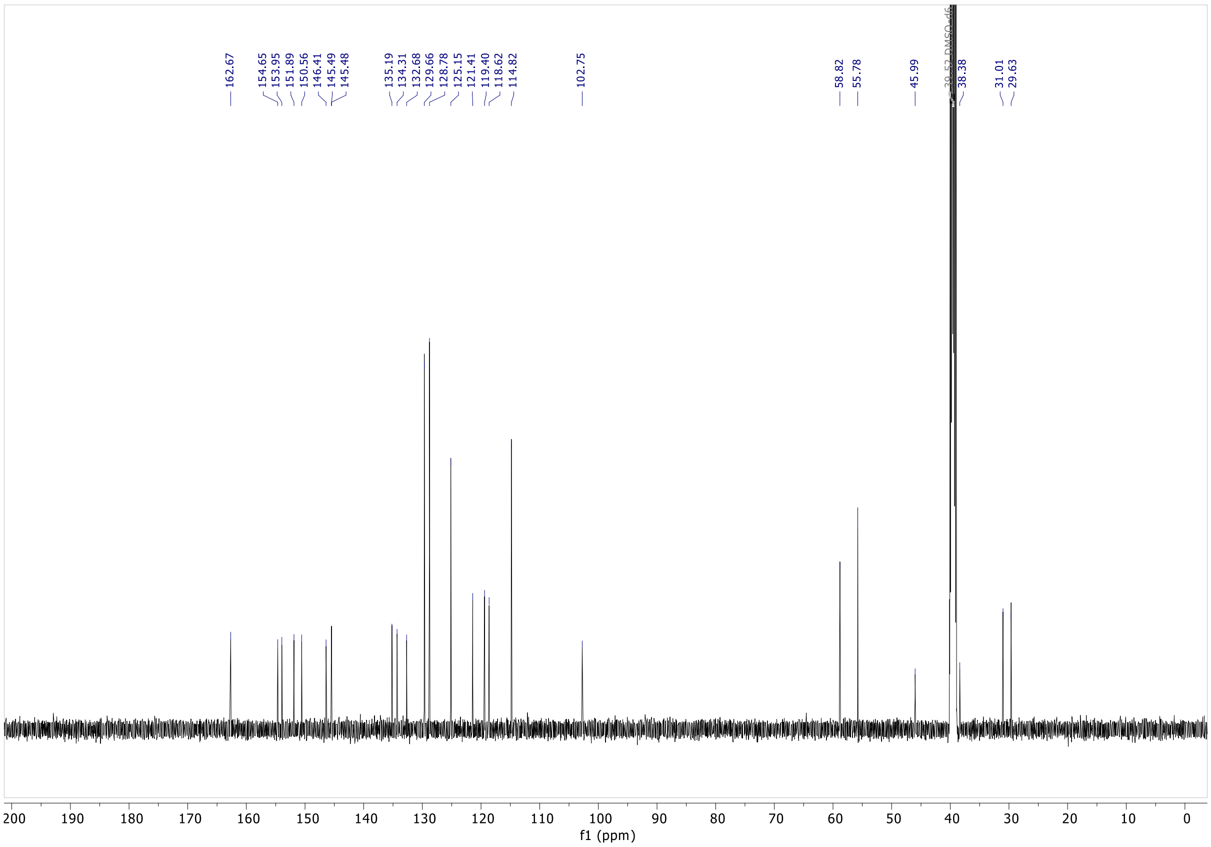

#### AzHC3

**¹H NMR** (400 MHz, DMSO-*d6*)

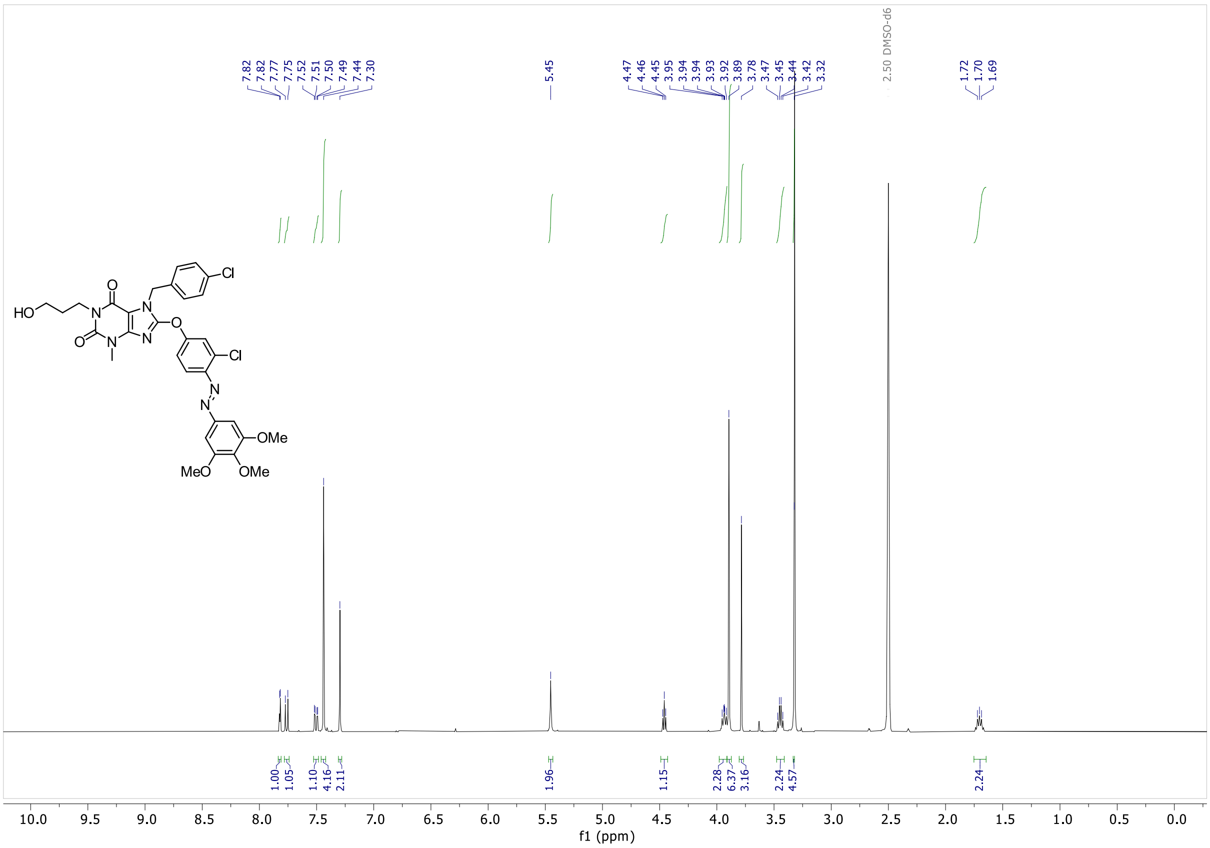

**¹³C NMR** (101 MHz, DMSO-*d6*)

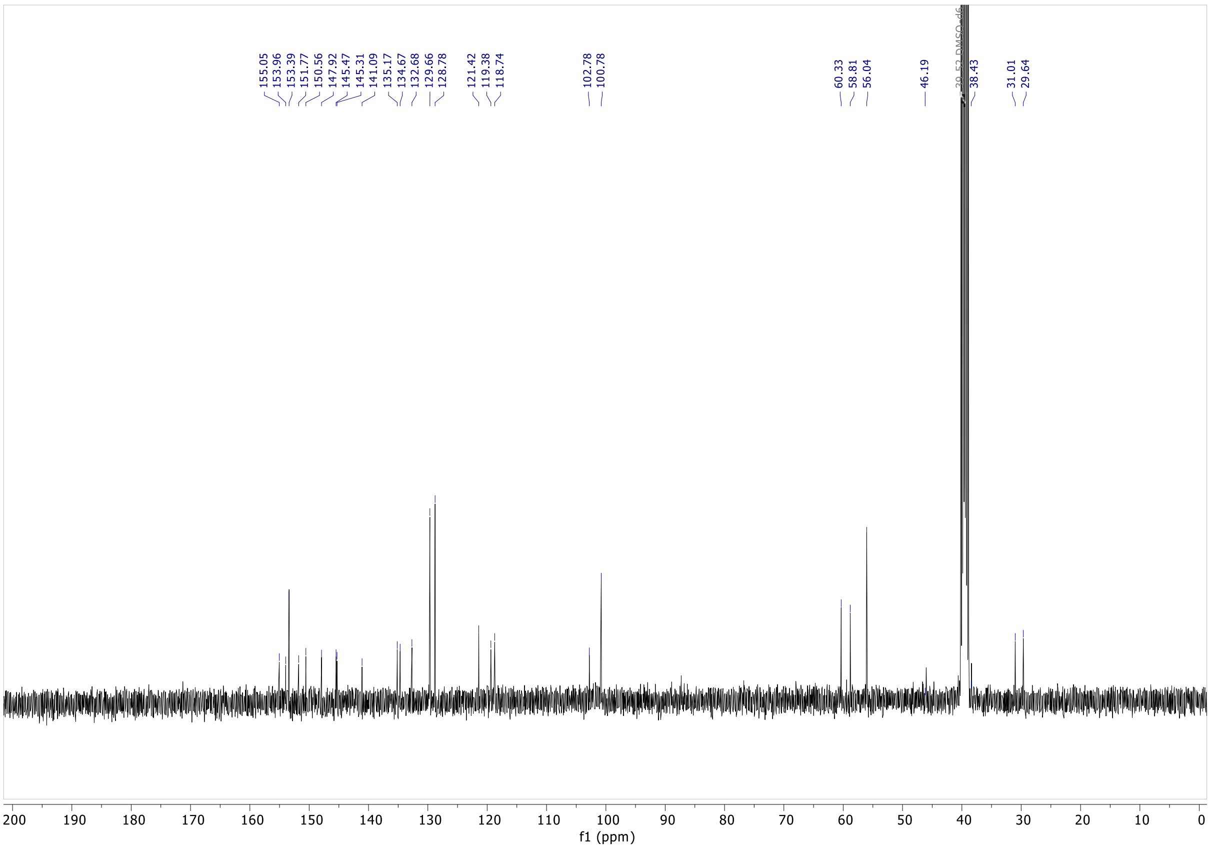

#### AzHC4

$^{1}$**H NMR** (400 MHz, DMSO-*d6*)

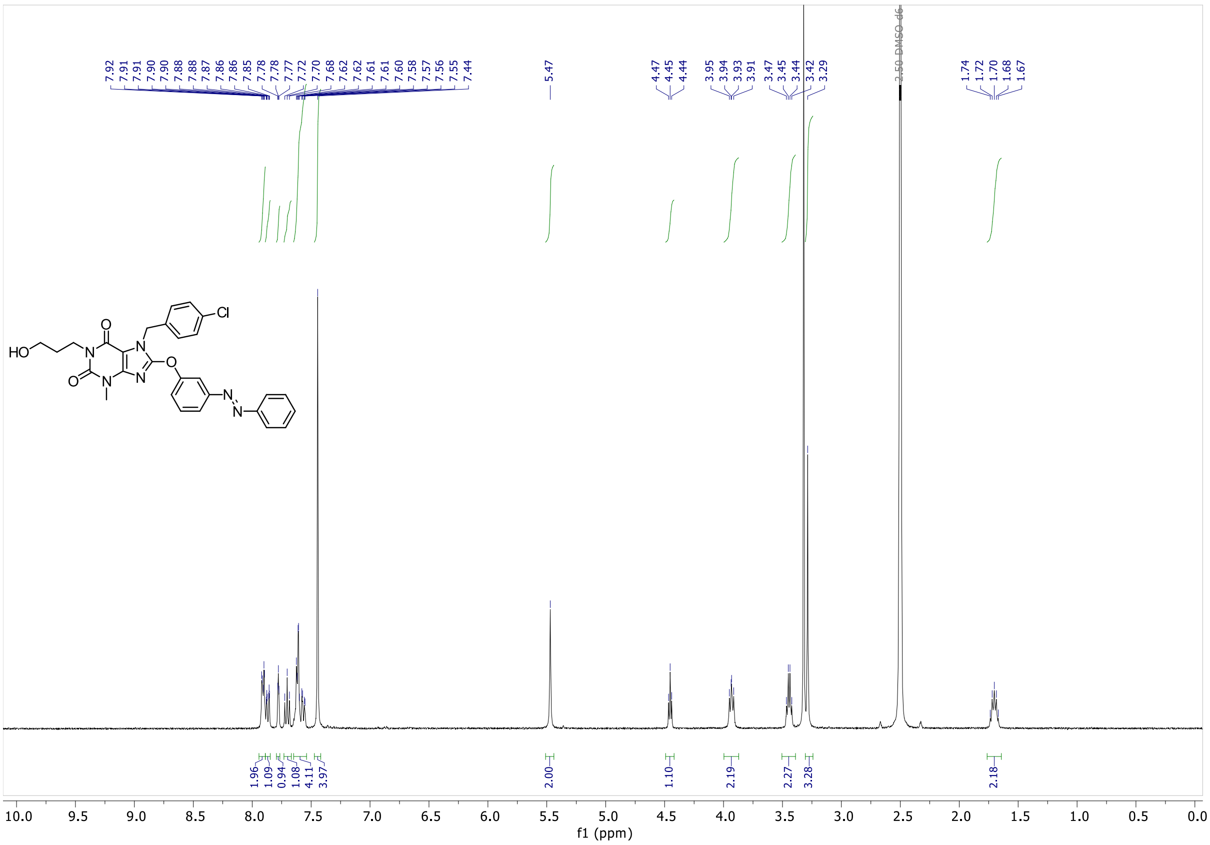

$^{13}$**C NMR** (101 MHz, DMSO-*d6*)

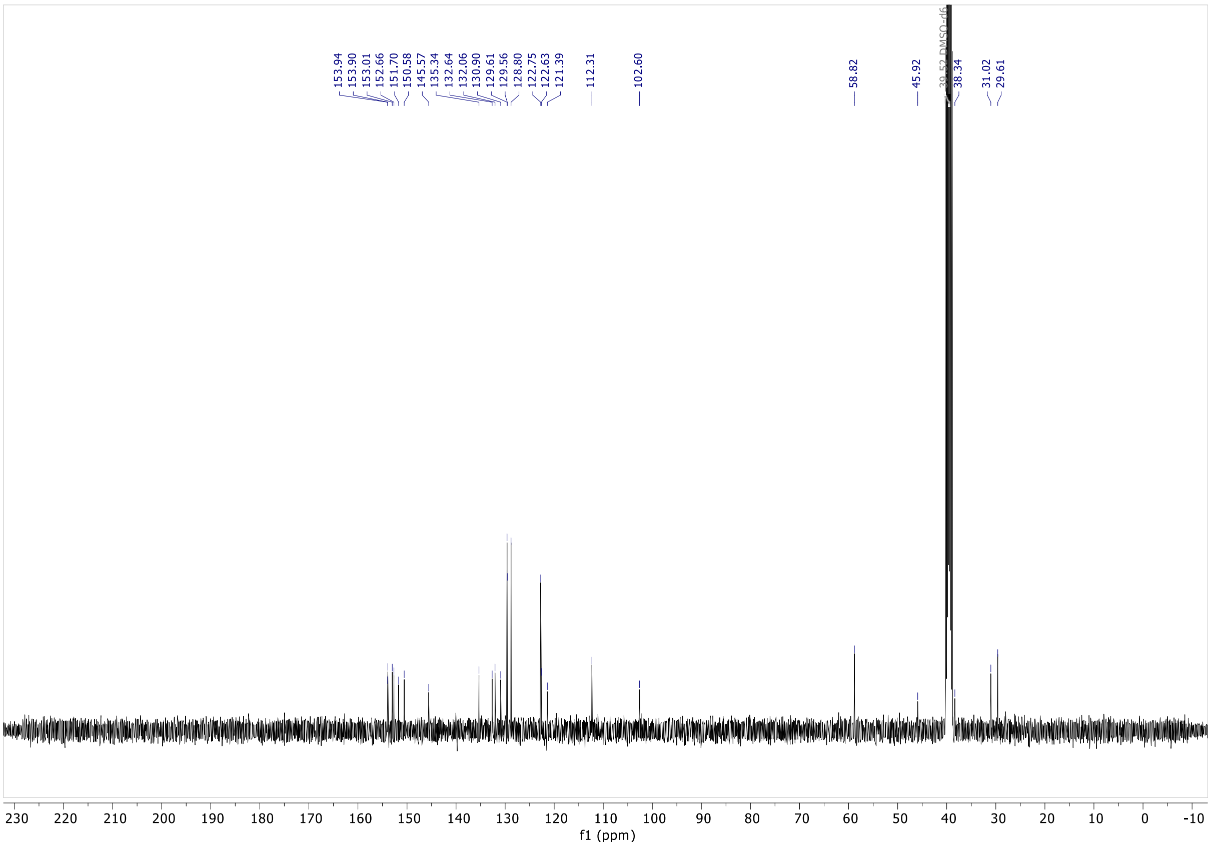

#### AzHC5

**$^{1}$H NMR** (400 MHz, DMSO-*d6*)

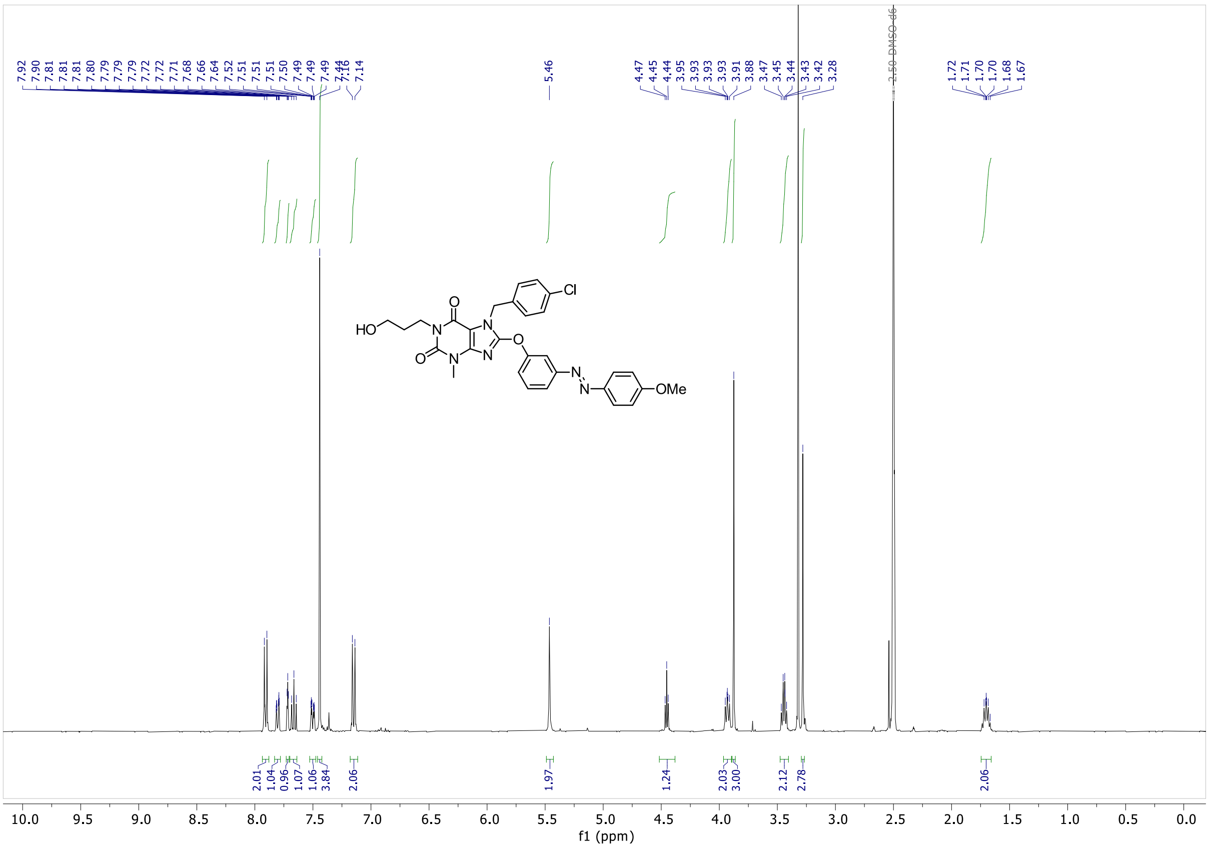

**$^{13}$C NMR** (101 MHz, DMSO-*d6*)

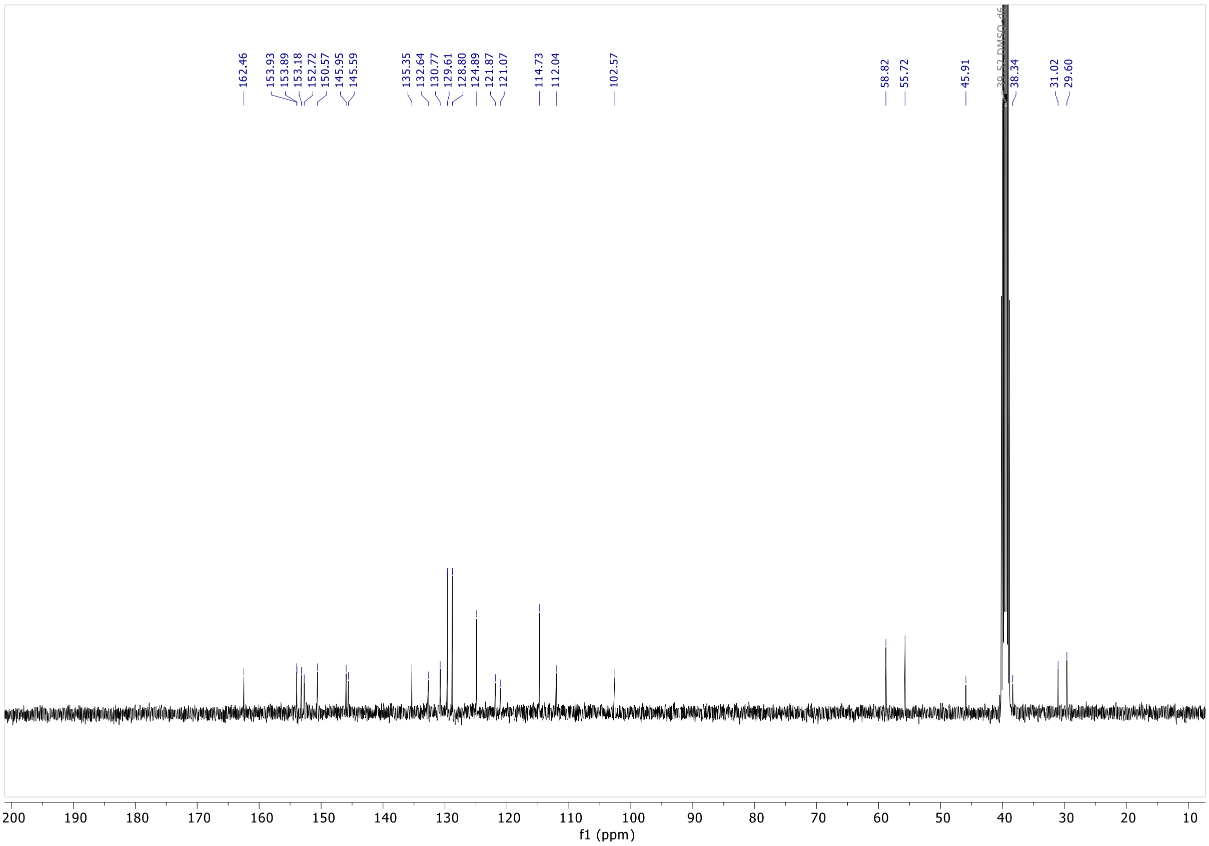

#### AzHC6

**$^{1}$H NMR** (400 MHz, DMSO-*d6*)

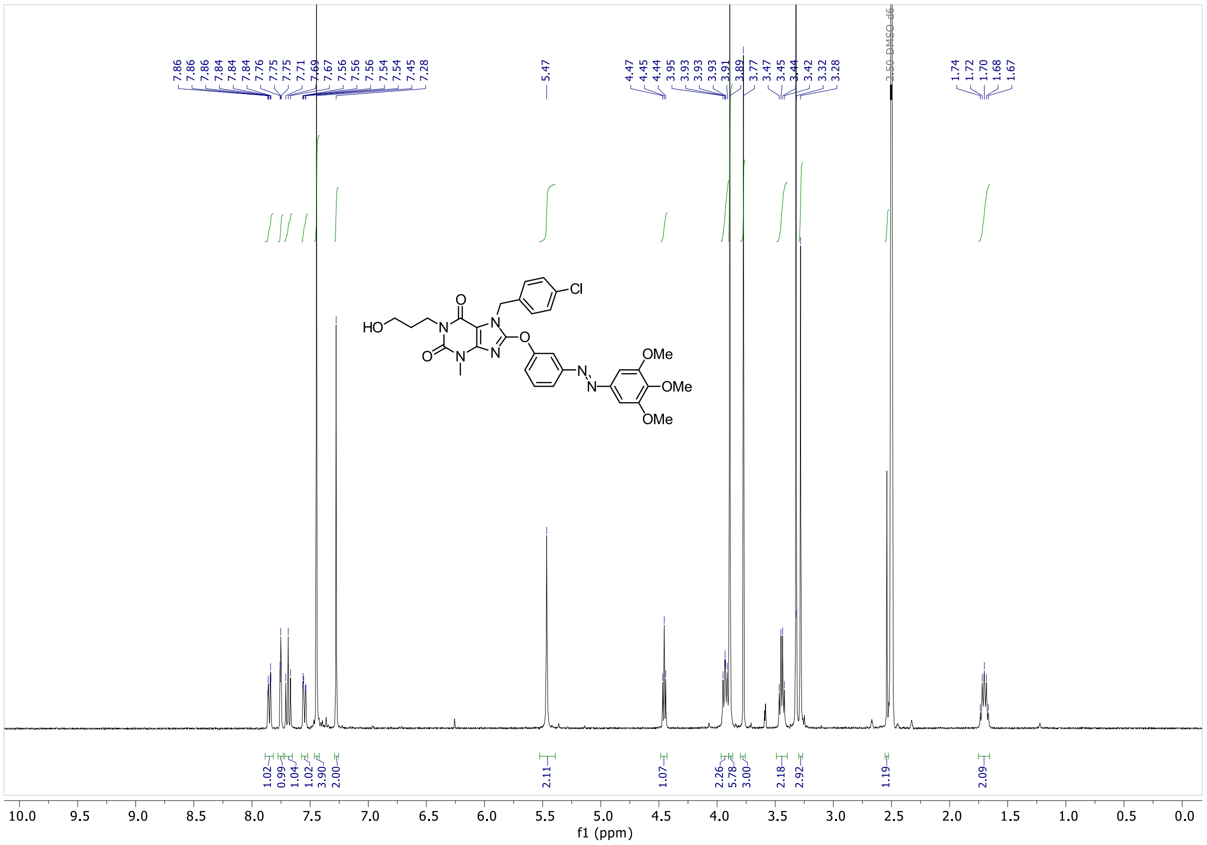

**$^{13}$C NMR** (101 MHz, DMSO-*d6*)

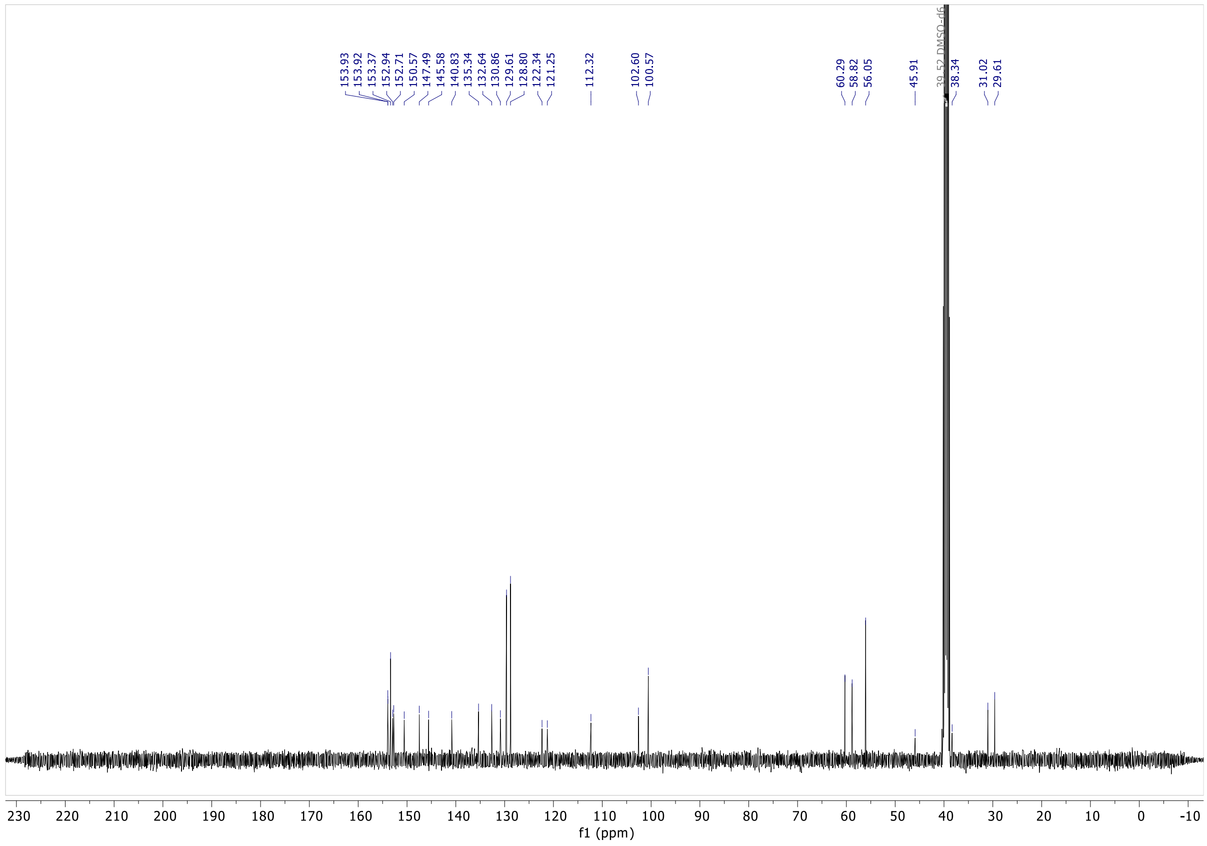

### 11 Copies of Main Figures With Full Legends

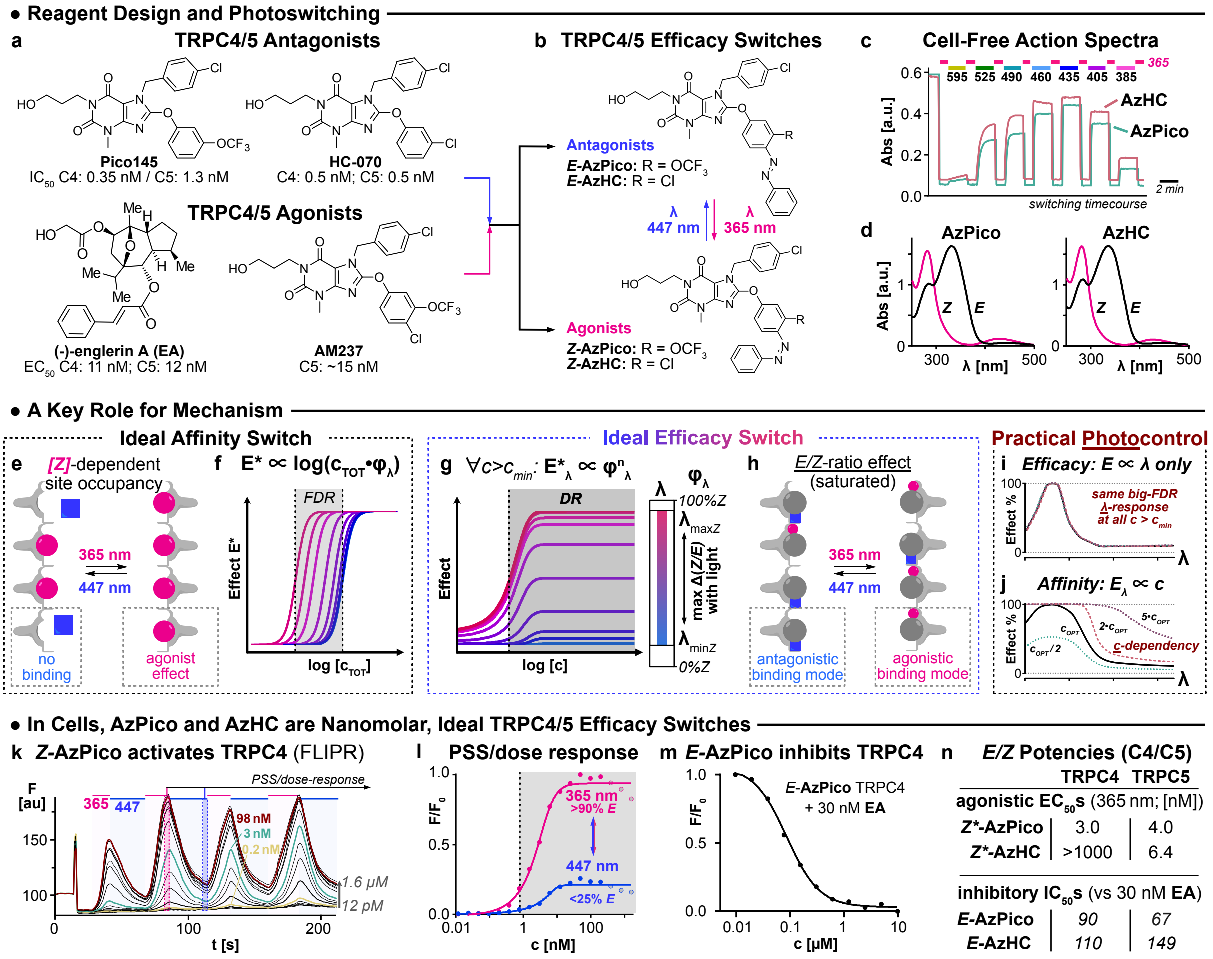

**Figure S23 (copy of main figure 1): Efficacy photoswitches for TRPC4/5. a**, Known TRPC4/5 modulators. **b**, Photoswitchable TRPC4/5 modulators **AzPico** and **AzHC**. **c-d**, Photoisomerisation action spectra and *E/Z* isomer absorption spectra of **AzPico** and **AzHC**. **e**,**f**, For an ideal affinity switch, only one isomer binds the target. Within the FDR window, binding site occupancy and thus biological effect **E*** depend on the total switch concentration $\mathbf{c_{TOT}}$ and the PSS fraction of active isomer $\boldsymbol{\varphi_\lambda}$. **g**,**h**, For an ideal efficacy switch, both isomers bind, with similar affinities, but with different efficacies. The dynamic range (DR) where the biological effect $E^*_\lambda$ is PSS-dependent but concentration-independent covers all $c > c_{min}$. **i**,**j**, Photocontrol in practice: for an efficacy switch, small variations in PSS(λ) sensitively control performance (whereas in affinity switches they are unimportant); but even large variations in concentration, which would ruin the performance of an affinity switch, are irrelevant. **k**,**l**, Reversible $Ca^{2+}$ influx modulation with **AzPico** under 365/447 nm cycles, as timecourse and peak amplitudes. **m**, *E*-**AzPico** binds competitively to **EA**. **n,** $EC_{50}$ and $IC_{50}$ values of *E/Z*-**AzPico** & **AzHC** on TRPC4 and TRPC5. (**k-n**, Fluo-4-loaded $HEK_{mTRPC4\beta}$ cells).

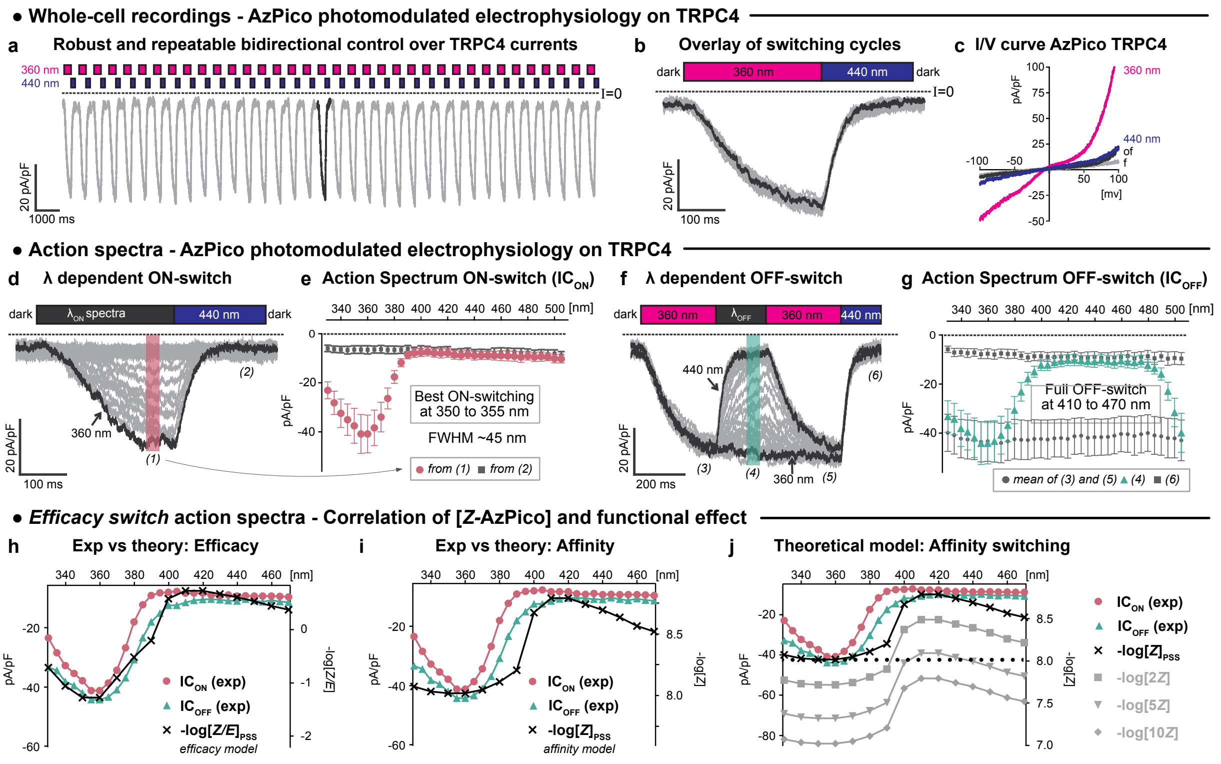

**Figure S24 (copy of main figure 2 with extended legend): AzPico-photocontrolled electrophysiology of TRPC4. a-g,** Electrophysiological whole-cell recordings of TRPC4 currents in voltage clamp mode (**a**,**b**,**d-g**: *V*h = -80 mV; **c**: *V*h scan) in HEK293 cells with 10 nM **AzPico** during photoswitching. **a-b**, Reproducibility of 36 consecutive photoswitching cycles of 360/440 nm (**a**: time-course; **b**: overlay of all cycles). **c**, *I*/*V* curves show that 440 nm drives almost full return to baseline currents throughout the applied voltage range. **d-g**, Spectral scans to extract the wavelength dependency of channel current photoswitch-on (**d-e**, cycles of $\lambda_{ON}$/440 nm) and photoswitch-off (**f-g**, 360 nm/$\lambda_{OFF}$). Panel **e** shows action spectra derived from **d** (plotted: inward currents at timepoints (1) for $\lambda_{ON}$, (2) for 440 nm); panel **g** shows action spectra derived from **f** (plotted: at timepoints (2) for $\lambda_{OFF}$, means of (1) and (3) for 360 nm, and (4) for 440 nm. In panel **f**, the reference cycle at 360/440 nm after each 360 nm/$\lambda_{OFF}$ scan cycle was incorporated so as to be able to exclude that artifacts from e.g. run-down or prolonged channel opening might be present. **h-j**: Ephys action spectra of **AzPico** match PSS-informed expectations for an efficacy switch (**h**), not an affinity switch (**i**). This is important, since an affinity switch would have severe concentration dependency (**j**).

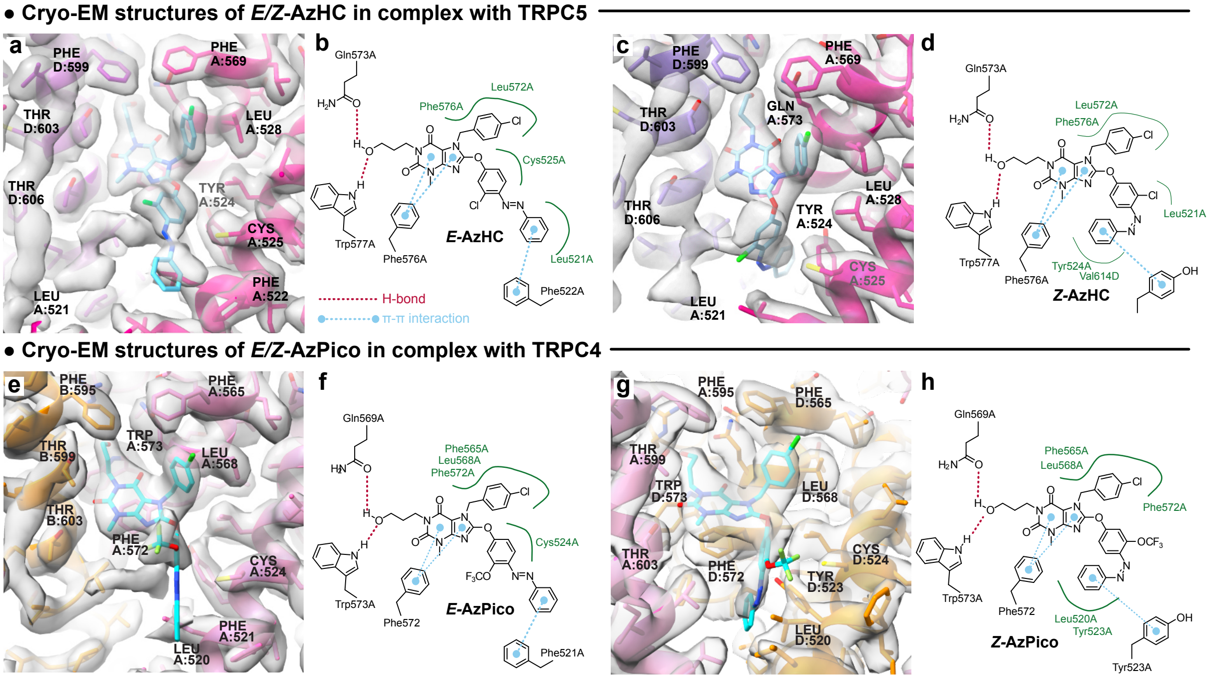

**[Figure S25](#) (copy of main figure 3 with more detailed legend): Structures of human TRPC4/5 in complex with *E/Z*-isomers of efficacy photoswitches.** (a) hTRPC5 in complex with (*E*)-**AzHC** (2.6 Å; C1 symmetry). The best-resolved xanthine binding site is highlighted. (b) 2D map of molecular interactions between (*E*)-**AzHC** and TRPC5 residues. (c) hTRPC5 in complex with (*Z*)-**AzHC** (3.1 Å; C1 symmetry). The best-resolved xanthine binding site is highlighted. (d) 2D map of molecular interactions between (*Z*)-**AzHC** and TRPC5 residues. (e-g) hTRPC4:*E/Z*-**AzPico** (*E:* xx Å; PDB: xxxx; *Z:* xx Å; PDB: xxxx; C1 symmetry (codes to be added upon paper acceptance)).

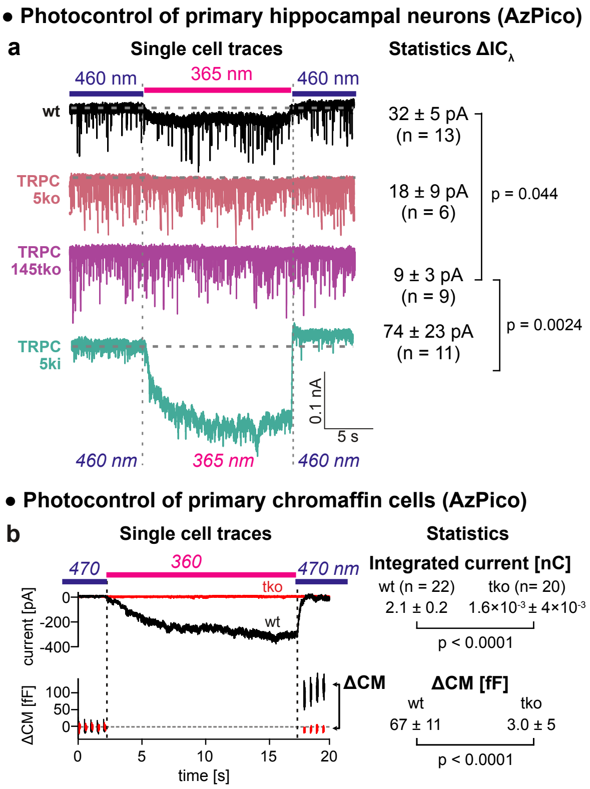

**[Figure S26](#) (copy of main figure 4): AzPico** (30 nM) photoswitchably evokes currents in primary neuronal and neuroendocrine cells with endogenous TRPC levels (single cell traces at left, group statistics at right). **a,** photoswitching-based current differentials **$\Delta IC_\lambda$** in hippocampal neurons (currents at 365 nm relative to 460 nm). **b,** photoswitch-based currents and net charge transfer (top) correlate to membrane capacitance changes **ΔCM** (bottom), indicating that phototriggering of TRPC[1]/4/5 leads to exocytosis (details at **Figure S20**).

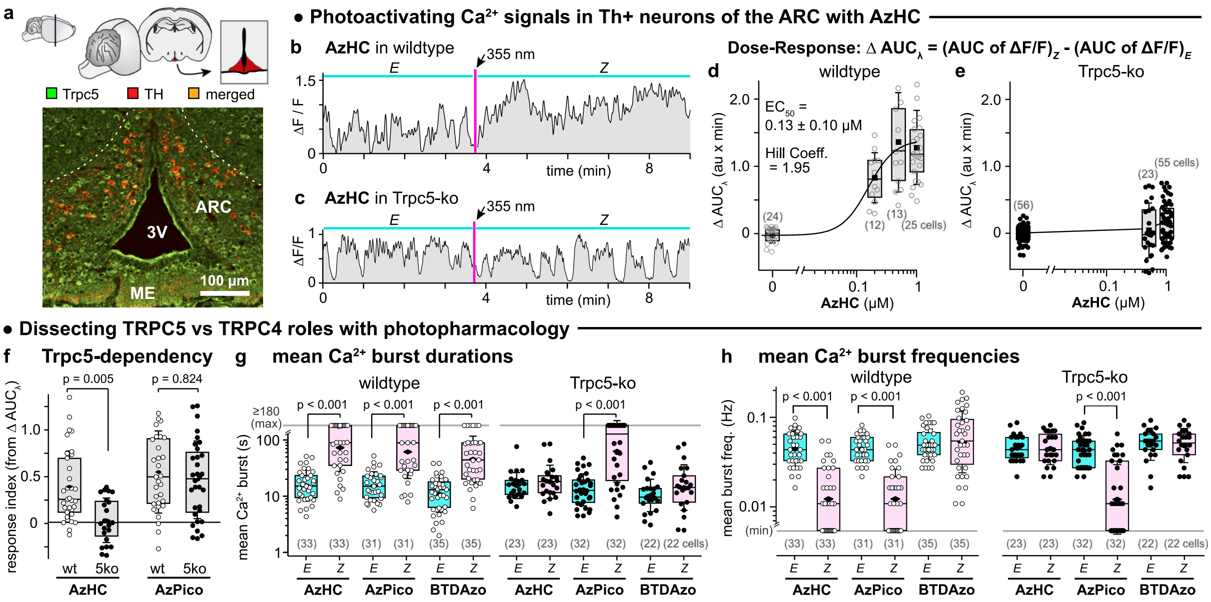

**Figure S27 (copy of main figure 5 with extended legend): AzHC and AzPico are potent photoswitchable activators of TRPC-dependent $Ca^{2+}$ responses in mouse hypothalamus. a,** immunohistochemical (IHC) image of TRPC5 (*green*) and Th+ neurons (*red,* tdTomato) in a brain slice of a Th-tdTomato mouse. 3V, third ventricle; ME, median eminence. Th+ neurons of the dorsomedial ARC express TRPC5 (*yellow*, merged). Cartoon of the coronal brain slice and region of the ARC (red) are indicated above the IHC image. **b-h,** $Ca^{2+}$ responses in Th+ neurons of Th-GCaMP6f (wildtype or wt) or Th-GCaMP6f-ΔTrpc5 (Trpc5-ko or 5ko) mice. **b,c,** Single-cell $Ca^{2+}$ traces before ("*E*") and after ("*Z*") a 355 nm pulse (default 67 ms long). **d,e,** $\Delta AUC_\lambda$: light-dependency of the area under the curves as acquired in (**b,c**). **f, AzHC** only photocontrols TRPC5-dependent $Ca^{2+}$ responses; **AzPico** can photocontrol $Ca^{2+}$ responses by another route (likely TRPC4) (the numbers of cells in each group are 33 & 23 (**AzHC** wt & 5ko), 31 & 32 (**AzPico** wt & 5ko)). **g,h,** mean $Ca^{2+}$ burst durations and frequencies in wt and 5ko upon *E*→*Z* photoswitching. The maximum measurement time window of 180 s results in **boundary values** e.g. when a cell has high signal throughout, its burst duration is capped at 180 s (frequency 0.006 Hz). For the sake of statistics, these values are taken literally in the calculation of the mean, median, SD, and statistical significance. Since boundary values only arise for *Z*-switches but not the *E*-switches to which their statistical significance is calculated, the p-values presented are true upper bound *p*-values (higher statistical significance would be found if measurement were continued longer). Since the absolute size as well as the statistical significance of the *EZ* differentials in the results are clear (and since these are the goal of the entire paper), we are satisfied with the very conservative underestimate that this data and analysis convey, about the power of the reagents' photocontrol in biology. Other significant p-values are: (i) *Z*-**BTDAzo** vs *Z*-**AzHC**, p < 0.001; (ii) *Z*-**BTDAzo** vs *Z*-**AzPico**, p < 0.027. (Default settings: **AzHC**/**AzPico** at 500 nM except in dose-response; **BTDazo** at 10 µM; **d-h**: each point is 1 cell, (n) cells per group; box plots show interquartile ranges, median (line), mean (black rhombus), and SD whiskers; **f-h**: Kruskal-Wallis ANOVA, Dunn's *p* values; for min/max values and other details, see **Figure S21**).

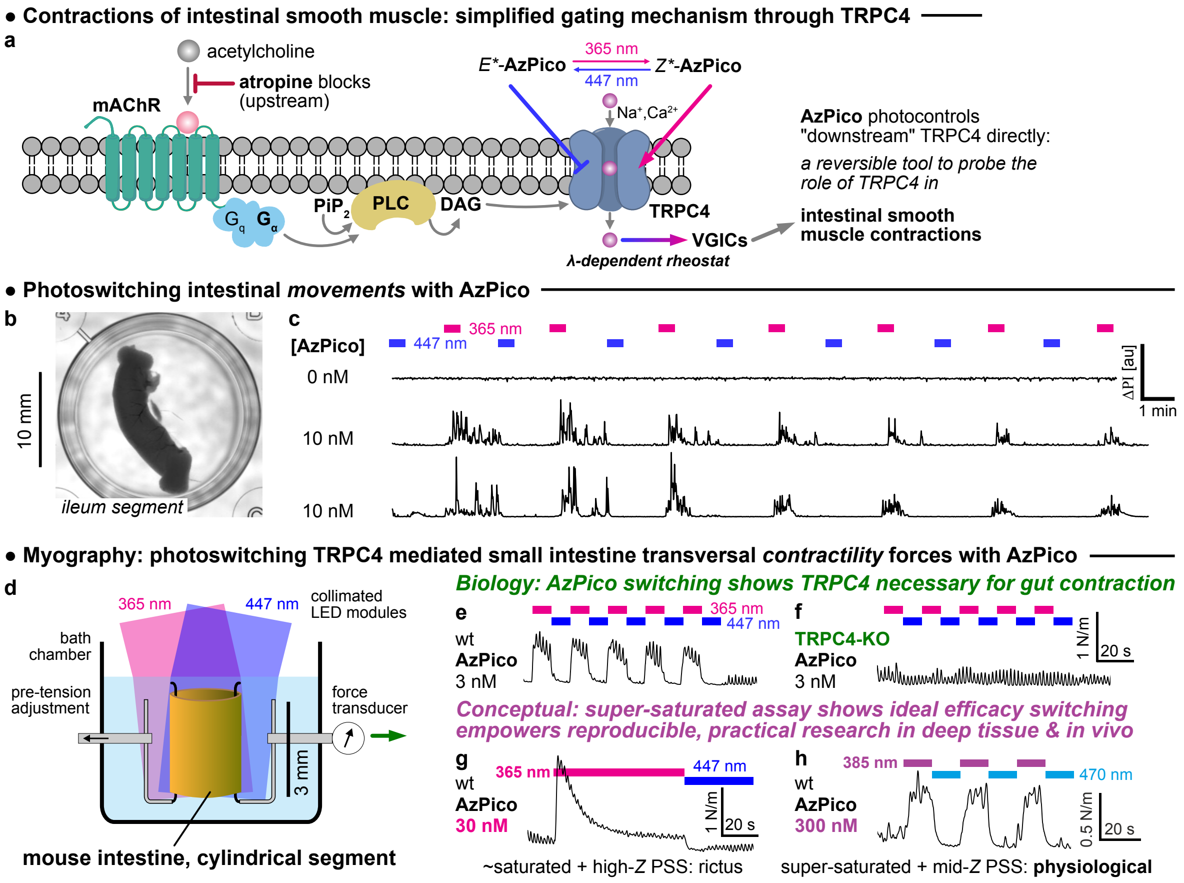

**Figure S28 (copy of main figure 6 with extended legend): AzPico photocontrol of macroscopic movements and coordinated contractility in intestine shows the key role of TRPC4 and the power of ideal efficacy switching**. **a,** simplified molecular mechanism for TRPC4-dependent intestinal contractility[81], that was now directly testable using **AzPico** (*E*/Z** indicate *E/Z*-rich-PSSs). **b**,**c** (see also **Movie S1**), sizeable intestinal segments (mouse ileum) whose motility was blocked by atropine (300 nM) but were treated with **AzPico** were driven into phases of fast macroscopic motions by 365 nm UV photoswitching (with longitudinal as well as ring muscle contractility), then returned to immobility by 447 nm blue light, reversibly over many cycles (narrow-band LED sources used, bandwidth FWHM ca. 30 nm). The metric used for assessment here is "ΔPI" i.e. the absolute value of the well-average instantaneous change of pixel intensity in the transmitted light image at time *t* compared to the previous frame (i.e. | I(t) – I(t-1) |, arbitrary units) which serves as a bias-free measure of gross displacement motion (an underestimate of total motion, since only covering or uncovering new pixels leads to large values). In the absence of **AzPico**, no photoresponse is detected. **Figure S22** shows additional traces and controls. **d-f**, physiological-like ring contractility is reversibly stimulated and suppressed by alternating 365 nm / blue illumination of segments treated with sub-saturating **AzPico**, in a TRPC4-dependent manner (knockout not photoresponsive). **g,h**, the ideal efficacy switch paradigm allows fully reproducible control of deep tissue bioactivity, by leveraging the *saturation* of dose-and-photon-flux (hard to titrate) but the *selection* of wavelength (easy to choose). Whereas roughly saturating concentrations of **AzPico** lead to over-stimulation under optimal-*Z* illumination at 365 nm, highly saturated **AzPico** can instead be used without ovestimulation just by dialling in a PSS with slightly lower *Z* content (385 nm).
